# Identifying latent behavioral states in animal movement with M4, a non-parametric Bayesian method

**DOI:** 10.1101/2020.11.05.369702

**Authors:** Joshua A Cullen, Caroline L Poli, Robert J Fletcher, Denis Valle

**Author notes:** Corresponding Author: 136 Newins-Ziegler Hall, Gainesville, FL 32611. PO BOX 110410.

## Abstract

1. Understanding animal movement often relies upon telemetry and biologging devices. These data are frequently used to estimate latent behavioral states to help understand why animals move across the landscape. While there are a variety of methods that make behavioral inferences from biotelemetry data, some features of these methods (e.g., analysis of a single data stream, use of parametric distributions) may limit their generality to reliably discriminate among behavioral states.
2. To address some of the limitations of existing behavioral state estimation models, we introduce a non-parametric Bayesian framework called the mixed-membership method for movement (M4), which is available within the open-source `bayesmove` R package. This framework can analyze multiple data streams (e.g., step length, turning angle, acceleration) without relying on parametric distributions, which may capture complex behaviors more successfully than current methods. We tested our Bayesian framework using simulated trajectories and compared model performance against two segmentation methods (behavioral change point analysis (BCPA) and segclust2d), one machine learning method (expectation-maximization binary clustering (EMbC)), and one type of state-space model (hidden Markov model (HMM)). We also illustrated this Bayesian framework using movements of juvenile snail kites (*Rostrhamus sociabilis*) in Florida, USA.
3. The Bayesian framework estimated breakpoints more accurately than the other segmentation methods for tracks of different lengths. Likewise, the Bayesian framework provided more accurate estimates of behavior than the other state estimation methods when simulations were generated from less frequently considered distributions (e.g., truncated normal, beta, uniform). Three behavioral states were estimated from snail kite movements, which were labeled as ‘encamped’, ‘area-restricted search’, and ‘transit’. Changes in these behaviors over time were associated with known dispersal events from the nest site, as well as movements to and from possible breeding locations.
4. Our non-parametric Bayesian framework estimated behavioral states with comparable or superior accuracy compared to the other methods when step lengths and turning angles of simulations were generated from less frequently considered distributions. Since the most appropriate parametric distributions may not be obvious *a priori*, methods (such as M4) that are agnostic to the underlying distributions can provide powerful alternatives to address questions in movement ecology.

## 1. Introduction

Our understanding of animal movement has advanced considerably in recent decades with the emergence of the field of movement ecology (Fraser et al. 2018; Joo, Picardi, et al. 2020), which focuses on understanding where animals go, what they are doing, and how they are influenced by their surrounding environment (Nathan et al. 2008). As telemetry and biologging devices continue to increase in their battery life, data resolution, and affordability (Hussey et al. 2015; Kays et al. 2015), statistical methods that can efficiently analyze these large datasets will become ever more important (Patterson et al. 2017; Potts et al. 2018). To fully understand animal movement, it is necessary to account for behavior since space and resource use are directly linked to an animal’s internal state (Nathan et al. 2008; Gurarie et al. 2016).

Since the direct observation of animal behavior can be challenging in many situations, recorded tracks from biologging devices are increasingly used to infer potential behavior by estimating latent states. These latent states can be estimated from a variety of data streams (i.e., time series of variables) such as step lengths, turning angles, ambient temperature, and acceleration, among others (Edelhoff et al. 2016). State estimation is often performed using segmentation and clustering methods, as well as state-space models (SSMs). Segmentation methods partition tracks into segments by detecting shifts in the data stream(s), whereas clustering methods classify these segments (or the observations directly) into discrete states. Alternatively, SSMs estimate latent states per observation based on the transition probabilities among a given number of states (Edelhoff et al. 2016; Gurarie et al. 2016). While existing state estimation methods provide fast or powerful predictive capacity (Edelhoff et al. 2016; Patterson et al. 2017), they possess a number of limitations that can impact the inference made on behavioral states.

For instance, segmentation methods commonly infer behavior using only a single data stream such as persistence velocity or speed (Edelhoff et al. 2016; but see Patin et al. 2020). This can be problematic when underlying behaviors are complex and not well represented by a single metric alone. Additionally, many segmentation methods, clustering methods, and SSMs typically estimate behavioral states by fitting the data streams to parametric probability distributions (e.g., Edelhoff et al. 2016; Patterson et al. 2017; Joo, Boone, et al. 2020), such as Gaussian, gamma, or wrapped Cauchy distributions. When the structure in the data streams is not well captured by parametric distributions, this can often result in overestimation of the true number of states when information criteria are used due to model misspecification (Gurarie et al. 2016; Pohle et al. 2017). Furthermore, running SSMs and some clustering methods can be computationally costly: model runtime can take minutes to days depending on the type of model, sample size, number of estimated states, and computer hardware. This is further exacerbated when model selection (e.g., determining the likely number of groups by fitting models with different numbers of groups) and multi-model inference are performed.

Given the limitations posed by existing state estimation methods, there is a need to develop a framework that is based on as few parametric assumptions as possible while also being fast and flexible. Here, we introduce a new two-stage modeling framework called the mixed-membership method for movement (M4) that implements non-parametric Bayesian methods to: 1) segment multiple data streams into relatively homogeneous units of behaviors; and 2) subsequently determine the likely number of behavioral states using a mixed-membership method where segments can be comprised of more than one behavioral state. Latent behavioral states are estimated for entire track segments (as opposed to individual observations) since this reflects our understanding that behavior is inherently autocorrelated, especially when observations are sampled at short time intervals (Pohle et al. 2017; Potts et al. 2018). Additionally, track segments are expected to be characterized by multiple states (Pohle et al. 2017; Patin et al. 2020). This M4 model framework is available within the open-source R package `bayesmove` available on CRAN (Cullen and Valle 2021). In this article, we describe the model structure and the Bayesian sampling methods used to estimate from the posterior distribution. We then demonstrate that M4 can successfully recover breakpoints and behavioral states based on simulated trajectories and compare our model’s performance against two common segmentation methods (behavioral change point analysis (BCPA), Gurarie et al. 2009; segclust2d, Patin et al. 2020), one machine learning method (expectation-maximization binary clustering (EMbC), Garriga et al. 2016), and one type of SSM, a hidden Markov model (HMM). Finally, we illustrate our novel approach on the movements of an endangered raptor species, the Everglade snail kite (*Rostrhamus sociabilis*), and interpret the results within the context of natal and breeding dispersal events.

## 2. Materials and Methods

### 2.1 Model Structure

Most existing segmentation methods (e.g., BCPA, segclust2d, behavioral movement segmentation), some machine learning methods (e.g., EMbC), and most SSMs (e.g., HMMs, multistate random walks) experience one or more common limitations to behavioral state estimation. These limitations include the reliance on parametric distributions, analysis of only a single data stream, as well as reliance on information criteria to determine the most likely number of states.

#### 2.1.1 Discretization of data streams

We address the problem of parametric distributions by providing an approach that relaxes parametric assumptions through the discretization of data streams (Figs 1a, b, c). Although data streams (e.g., step lengths and turning angles) are not typically discretized into bins, we expect that this may lead to more robust estimates in the face of parametric distribution uncertainty. This is because bins are estimated independently of one another and extreme values lose their influence when added to the first or last bins with the rest of the data. Therefore, the discretization of data streams is expected to increase model flexibility (Kitagawa 1987; John and Langley 1995).

**Figure 1.**
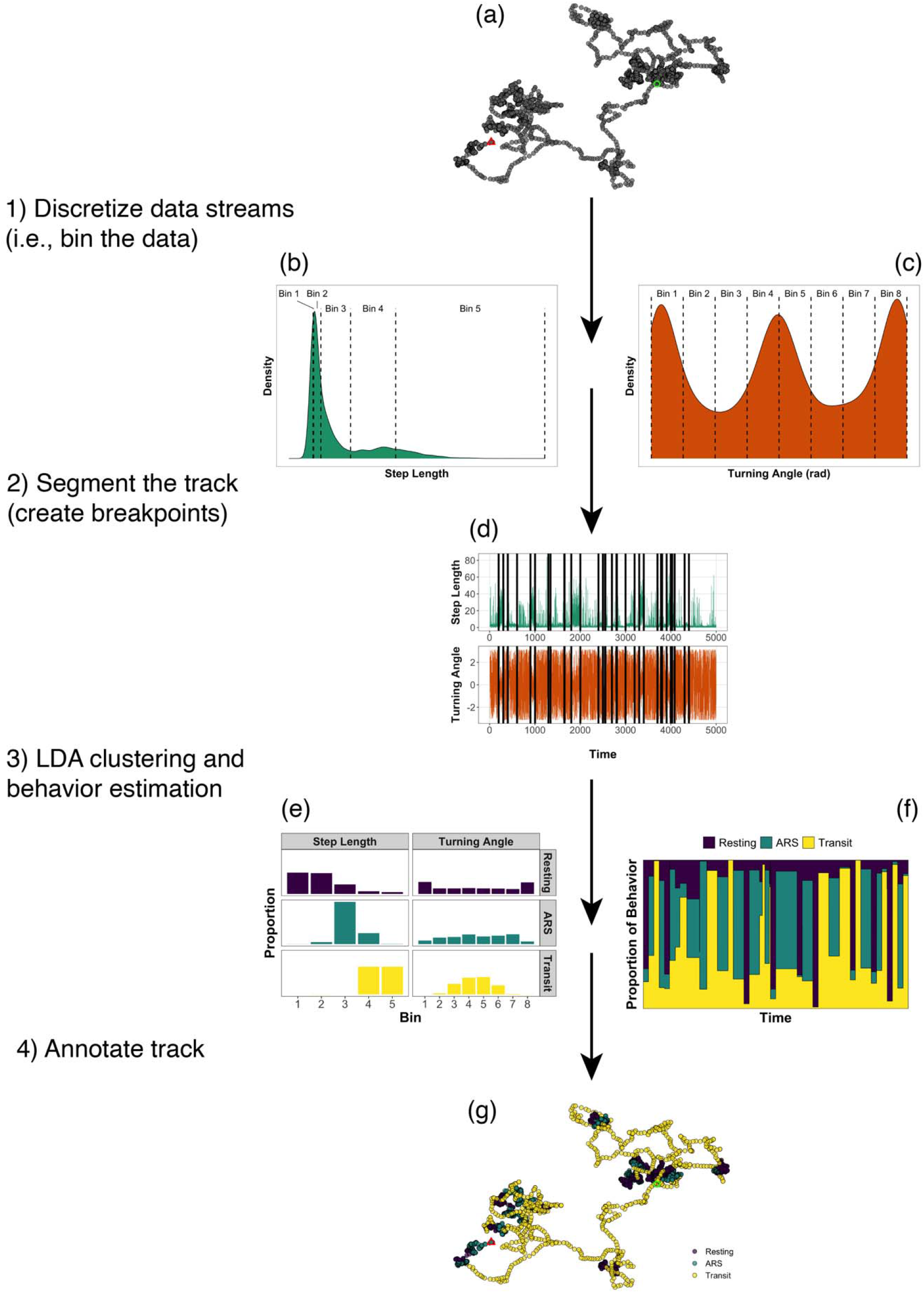
General workflow to analyze animal trajectories from telemetry data using M4. Steps from this analysis include: (a) starting with a track of coordinates that are used to calculate step lengths and turning angles, (b) discretizing step lengths and (c) turning angles into sets of bins, (d) performing segmentation on the joint time series of step lengths and turning angles, (e) cluster the track segments into behavioral states by Latent Dirichlet Allocation (LDA) (matrix Φ) and (f) evaluate time series of behavior proportions for each individual (matrix Θ), and (g) visualize the annotated tracks by displaying the dominant behavior of each time series. Note that continuous time series are only displayed with breakpoints (black lines) in (d) to improve interpretability of segments.

Selecting the number of bins and the binning method is relatively subjective and therefore it is important that prior biological reasoning be used to inform these decisions. For example, discretization methods may include the use of equal bin widths or quantiles. However, the number of bins should be sufficient to characterize the shape of the density distribution. These assumptions during the discretization process are not unlike assumptions made for HMMs when selecting probability distributions to fit data streams, but require practitioners to make more decisions up front. Based on a sensitivity analysis of binning methods used on a right-skewed data stream (i.e., step lengths), the use of quantiles resulted in greater discrimination of behavioral states than bins of equal widths (Appendix S1). However, data streams with circular distributions (e.g., turning angles) will likely be more interpretable when using bins of equal widths. At a minimum, discretized values for each data stream and the associated track IDs are required to begin analyzing the data.

#### 2.1.2 Segmentation

Similar to other segmentation methods, our model aims to divide tracks into segments by estimating a set of breakpoints. Through the use of joint probabilities within a Bayesian framework, we circumvent the limitation of analyzing only a single data stream. Moreover, our model estimates the location and number of unknown breakpoints by implementing a Gibbs sampler within a reversible-jump Markov chain Monte Carlo (RJMCMC) algorithm. RJMCMC is a trans-dimensional algorithm that serves as a model-based approach to model selection by providing simultaneous inference on parameter values given a particular model, as well as model space (i.e., the collection of all possible models) (Green 1995). In particular, we use a birth-death RJMCMC that allows the addition (i.e., birth), removal (i.e., death), or swap of proposed breakpoints where model parameters are updated from the known posterior distribution using a Gibbs sampler (see Appendix S2 for more details). We adopt this approach to perform unsupervised segmentation on each individual trajectory. In our framework, each potential model *M_k_* is characterized by a set of *P* breakpoints {*b*_1*k*_,…, *b_Pk_*}, where *k* is the model number. Each breakpoint is restricted to being an integer between 2 and *T*_*i*−1_ across all observations, where *T_i_* is the total number of observations for individual *i*. Given a particular model *M_k_*, its breakpoints define track segments. We assume that for any given track segment *c*:

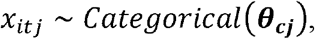

where *x_it j_* is the bin label for individual *i* at time *t* for data stream *j* and ***θ_cj_*** is a vector of probabilities that sum to one. The vector ***θ_cj_*** indicates the probability that observations within segment *c* are assigned to one of *L* bins. Overall, the model is seeking breakpoints that define relatively homogeneous track segments. The use of a categorical distribution to characterize track segments (as opposed to continuous distributions) is what makes this framework non-parametric. Our prior is given by:

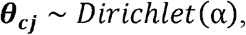

where α > 0 is equal across all bins. We integrate over the latent parameter ***θ_cj_*** to enable the algorithm to visit multiple models and increase computational efficiency. Since there are no longer any remaining parameters on which to assess model convergence, we do so by evaluating trace-plots of the log marginal likelihood (Denison et al. 2002). Details for the derivation of the full conditional distributions can be found in Appendix S3.

Since the posterior distribution of models *M_k_* can vary greatly in the number and position of breakpoints, but only a single set of breakpoints can be used to define track segments, we select the Maximum a Posteriori (MAP) estimate (i.e., the breakpoints of the model with the greatest log marginal likelihood) (Fig. 1d). Although the MAP estimate does not account for uncertainty in breakpoint number and position, it appears to be in good agreement with estimates from the entire posterior distribution as described in Appendix S4. These MAP breakpoints are then used to define segments per individual track, which are subsequently clustered into latent behavioral states by a mixed-membership model.

#### 2.1.3 Mixed-membership clustering

Although most existing state estimation methods assign a single discrete state to observations or track segments (e.g., Garriga et al. 2016; McClintock and Michelot 2018; Patin et al. 2020; but see Jonsen et al. 2019), animal movement may not be entirely comprised of a single behavior over a given sampling interval (Pohle et al. 2017; Patin et al. 2020). Latent Dirichlet Allocation (LDA), a mixed-membership clustering method, can be used to classify each track segment as a mixture of multiple states (Valle et al. 2014; Hudon et al. 2021). For example, a proportion of observations within a given track segment might belong to state 1 while another proportion might belong to state 2 and so on.

LDA is used to characterize track segments in terms of their behavioral state components, where each state corresponds to a distribution of discretized data streams. To do so, the model estimates the probability of observations from each track segment (rows) belonging to each latent state (columns) in matrix Θ (Fig. 1f). Additionally, the model characterizes the latent states (rows) with the probability of observations belonging to each bin per discretized data stream (columns) in matrix **Φ** (Fig. 1e). The track segments from all individual animals are analyzed together since we assume that there is a common set of behaviors exhibited across the population. Although there may be some individual heterogeneity in movement patterns, the pooling of all individuals ensures that behavioral states are directly comparable and improves the inference on individuals with fewer observations (Jonsen 2016). In this model, we assume that:

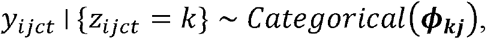

where denotes *y_ijct_* the bin for time *t* of track segment *c* from data stream *j* for individual *i*. Additionally, *z_ijct_* is the latent behavioral state membership associated with *y_ijct_* and ***ϕ_kj_*** is a vector of probabilities for each behavior and data stream. Notice that *z_ijct_* influences the distribution for *y_ijct_* by determining the subscript *k* for the vector ***ϕ_kj_***. We assume that the latent state membership is given by:

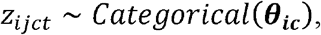

where ***θ_ic_*** is a vector of probabilities of size *K* (i.e., the number of clusters or states) that sum to one and indicates the likelihood of assigning an observation at time *t* of track segment *c* for individual *i* to each behavioral state *k*. This formulation assumes that each observation within a particular track segment must belong to a single behavioral state, but that track segments are comprised of multiple states. For our priors, we assume that:

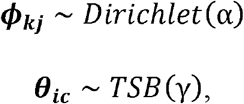

where *TSB*(*γ*) represents the truncated stick-breaking prior from Bayesian non-parametrics. This prior is given by:

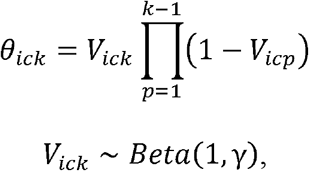

where *V_K_* = 1 and γ > 0. By setting 0 < *γ* < 1, we can shrink the probability of assigning state *k* to track segment *c* (i.e., *θ_ick_*) to approximately zero as *k* approaches *K*. As a result, fewer and fewer observations will be assigned to states with large values of *k*, enabling the model to identify the most likely number of behavioral states (Valle et al. In Review; Valle et al. 2017). This is an improvement on existing state estimation methods in the sense that our model only needs to be run once, whereas several other common methods (e.g., HMMs, segclust2d, and other clustering methods) are typically run multiple times with varying numbers of behavioral states to then determine the best model via information criteria (e.g., AIC or BIC).

This LDA model is fitted using a Gibbs sampler and a complete description of the full conditional distribution can be found in Appendix S5. Similar to the segmentation model, convergence was assessed by inspecting trace-plots of the log-likelihood. The posterior mean for all ***θ_ic_*** was then used to identify the most likely number of behaviors. The estimated statedependent distributions for each data stream (from ***ϕ_kj_***) were evaluated and used to corroborate the findings based on the posterior means from all ***θ_ic_***’s by determining whether the distributions were biologically relevant (Figs 1e, f, g). This combination of results provides a straightforward approach to selecting the most likely number of behavioral states. A list of the primary functions to analyze data using the M4 framework within the `bayesmove` R package is included in Appendix S6.

### 2.2 Simulation Study

We assessed the performance of M4 compared to other methods via simulations. We first evaluated the ability of our track segmentation method to detect true breakpoints and compared its results to those obtained by two segmentation methods (i.e., BCPA and segclust2d). We then evaluated the ability of our clustering method to estimate the true number of behavioral states and to properly assign behavior proportions to track segments. For this component, we compared the results of our model to those obtained by a HMM (McClintock and Michelot 2018) and two additional clustering methods (i.e., segclust2d and EMbC).

#### 2.2.1 Generating simulated trajectories

We generated multiple three-state trajectories from a correlated random walk at regular time intervals, where five tracks were simulated at each of four durations (1000, 5000, 10000, 50000 observations), resulting in a total of 20 tracks. Each track was comprised of 10, 50, 100, or 500 segments that each included 100 observations. Each of these segments included a dominant behavioral state (80% of observations), which was randomly assigned to each segment. The three behavioral states were parameterized to represent (1) little to no movement (‘encamped’), (2) slow and tortuous movement (‘area-restricted search’ or ARS), as well as (3) fast and directed movement (‘transit’). To assess model performance on tracks generated from different types of distributions, we generated two sets of tracks with 20 simulations in each. In the first set of simulations, step lengths for each behavior were drawn from a truncated normal distribution and turning angles were drawn from either a beta, uniform, or truncated normal distribution (hereafter referred to as ‘uncommon distributions’) (Fig. 2a). For the second set of simulations, step lengths for each behavioral state were generated from a gamma distribution and turning angles were drawn from a wrapped Cauchy distribution (hereafter referred to as ‘common distributions’) (Fig. 2b). Both sets of simulations were designed to generally resemble each other in their step length and turning angle distributions.

**Figure 2.**
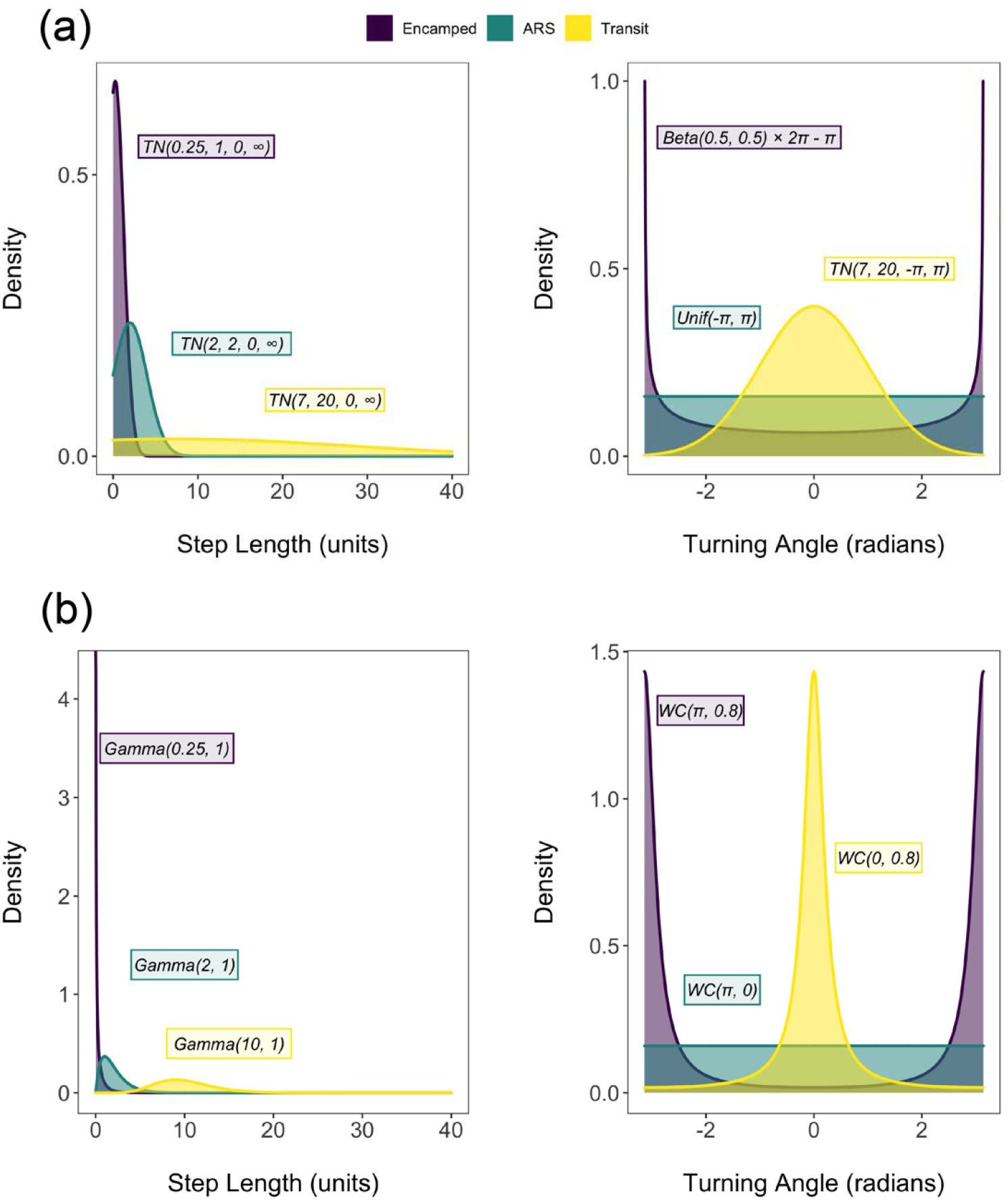
Uncommon (a) and common (b) distributions used to generate step lengths and turning angles for each simulated state. (a) Step lengths are generated from a truncated normal (TN) and turning angles are generated from beta, uniform, and truncated normal distributions. TN (a,b,c,d) denotes a truncated normal distribution with mean a, standard deviation b, lower bound c, and upper bound d. (b) Step lengths are generated from a gamma distribution and turning angles are generated from a wrapped Cauchy (WC) distribution.

#### 2.2.2 Implementation of Bayesian M4 framework

Step lengths and turning angles were the data streams used to make inference on latent behavioral states. Step lengths were separated into five bins using the 25^th^, 50^th^, 75^th^, 90^th^, and 100^th^ quantiles as upper limits. Quantiles were used to discretize highly right-skewed step lengths as suggested by our sensitivity analysis (Appendix S1). Turning angles were discretized into eight bins from *−π* to *π* using equal widths 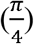 since the distribution of this variable was relatively balanced and bounded by lower and upper limits.

Each simulated track was analyzed by the M4 segmentation model using a vague prior where the hyperparameter α was set to 1. Trace-plots of the log marginal likelihood indicated that the model reached convergence for each simulation, where 40,000 MCMC iterations were used for all but the longest tracks, which used 60,000 iterations (Appendix S7). We then assessed how well our model identified the true breakpoints for each simulation, where a threshold of ±10 observations was used to distinguish an accurate from an inaccurate estimated breakpoint. If no breakpoints were estimated within ±30 observations of a true breakpoint, then the model was considered to have ‘missed’ that breakpoint. Other thresholds were tested and all resulted in the same relative pattern of accuracy (Appendix S1).

Estimated track segments were used as input for the LDA model of M4, which was run using 1000 MCMC iterations, a burn-in of 500 iterations, and vague priors where hyperparameters were set to γ = 0.1 and α = 0.1. We set the maximum number of behavioral states to seven since this was expected to include the true number of states, estimated via the truncated stick-breaking prior of the LDA. Trace-plots of the log-likelihood indicated that the model reached convergence for each simulation (Appendix S7). The true number of states was estimated by calculating the arithmetic mean of behavior proportions across all track segments and selecting the set of states that together represented ≥90% of all observations on average. Additionally, state-dependent distributions of step lengths and turning angles were inspected so that we only selected states that were also biologically interpretable. Since the LDA treats track segments as a combination of behavioral states, proportions of each state were estimated per track segment. Accuracy of state estimates were evaluated by two methods: 1) we calculated the percentage of observations where the dominant behavior of each track segment was accurately classified, and 2) we calculated the root mean square error (RMSE) of the estimated behavior proportions compared to the true behavior proportions over all states and track segments.

#### 2.2.3 Method comparison

We compared the performance of M4 on the simulated trajectories against BCPA, EMbC, HMM, and segclust2d (see Appendix S8 for details regarding model properties and assumptions). All models were run using a 2.6 GHz i7 CPU with 16 GB RAM.

##### 2.2.3.1 Segmentation models

The BCPA model performed segmentation based on persistence velocity (PV), which is a combination of velocity (*V*) and turning angle (θ) (i.e., *PV* = *Vcos*(θ)), using the R package `bcpa` v1.1 (Gurarie 2014). Parameters for BCPA were tuned to provide a close approximation of the true number and location of simulated breakpoints with window size set to 80, sensitivity set to 2, and clusterwidth set to 30. Breakpoint accuracy was evaluated using the same method as for M4.

The segclust2d model performed segmentation on step lengths and the absolute value of turning angles using the R package `segclust2d` v0.2.0 (Patin et al. 2019). This method models each data stream using a Gaussian distribution, so the absolute value of turning angles was used to accommodate this unimodal assumption. Tuning parameters were chosen within the bounds of the simulated tracks, such that the maximum number of segments was set to 1.5 × *s* where *s* is the true number of segments, the minimum observations per segment was set to 50, and the number of potential clusters (i.e., states) ranged from 2 to 4. Since the model was still analyzing the longest simulations (50000 observations) after two days, these tracks were omitted from the reported results for segclust2d. Breakpoint accuracy was assessed in the same manner as for M4.

##### 2.2.3.2 Clustering models

The EMbC model was fitted to step lengths and the absolute value of turning angles using the R package `EMbC` v2.0.3 (Garriga et al. 2019). The absolute value of turning angles was used to achieve better discrimination among states given the use of a unimodal distribution like the Gaussian distribution. This model uses binary clustering to partition each of *n* data streams into a ‘low’ and ‘high’ class, resulting in a total of 2^n^ possible states. For our analysis, this resulted in four states estimated from a bivariate Gaussian distribution. To make these results comparable to the other models, both states with ‘high’ step lengths (and ‘low’ or ‘high’ turning angles) were merged into a single state to produce three states overall. State classification accuracy was assessed at the segment-level so that results were directly comparable with the Bayesian M4 model. This was achieved by using true breakpoints to segment the time series of states estimated by the EMbC model and then calculating the proportion of these behaviors within each track segment. Additionally, the resulting state-dependent distributions of step lengths and turning angles were discretized using the bin limits defined for M4 to compare the accuracy of distribution shapes. Accuracy was measured by RMSE across bins of all states and data streams per simulation.

A discrete-time HMM was also fitted to each of the simulated trajectories using the R package `momentuHMM` v1.5 (McClintock and Michelot 2018). Step lengths were modeled using a gamma distribution and turning angles were assumed to arise from a wrapped Cauchy distribution. The HMMs for each simulation were run using a range of 2 to 4 possible behavioral states (*K*) and each *K*-state model was run 30 times using different starting values to increase the chance of finding the global (as opposed to local) maximum of the likelihood. The selection of “good” starting values is critical since it can affect computation time and the ability of the model to identify the global maximum of the likelihood (Michelot et al. 2016; McClintock and Michelot 2018). The most likely number of states was selected using a combination of AIC and BIC, where the model with the lowest value was considered to be most likely. However, if the difference in AIC or BIC (ΔAIC/BIC) of the next best model was < 10 (Burnham and Anderson 2002), the more parsimonious model was chosen. Behavior classification accuracy was assessed in the same manner as for EMbC.

The segclust2d model clustered segments previously estimated by this method into *K* states. The number of likely states was estimated using BIC in the same manner as for HMM. The likely number of states (and associated breakpoints) were used to assign behavioral states to track segments, which were then compared to the other methods using the proportion of each state per estimated segment (which were all either 0 or 1). Additionally, the accuracy of the state-dependent distributions were evaluated in the same manner as for EMbC and HMM.

### 2.3 Snail kite case study

As part of a larger investigation on the effects of wetland management on wildlife, solar-powered GPS-GSM transmitters (Ecotone Telemetry) were attached to juvenile snail kites (n=26) prior to fledging at Lakes Tohopekaliga, East Tohopekaliga, and Kissimmee in central Florida during 2018 and 2019. Subsequent movement of each individual resulted in a total of 40,720 observations (Fig. 3). Locations were collected once per hour only during daylight at an accuracy of ±30 m. As a result of the programmed duty cycle and time periods where GPS tags failed to transmit data, track time intervals were irregular. To ensure comparable step lengths and turning angles, we filtered our data to the most common time interval (i.e., 1 h). We chose to omit all other observations since imputation procedures for long time gaps would increase the number of artificial data and the use of linear interpolation would artificially inflate the number of turning angles at zero radians.

**Figure 3.**
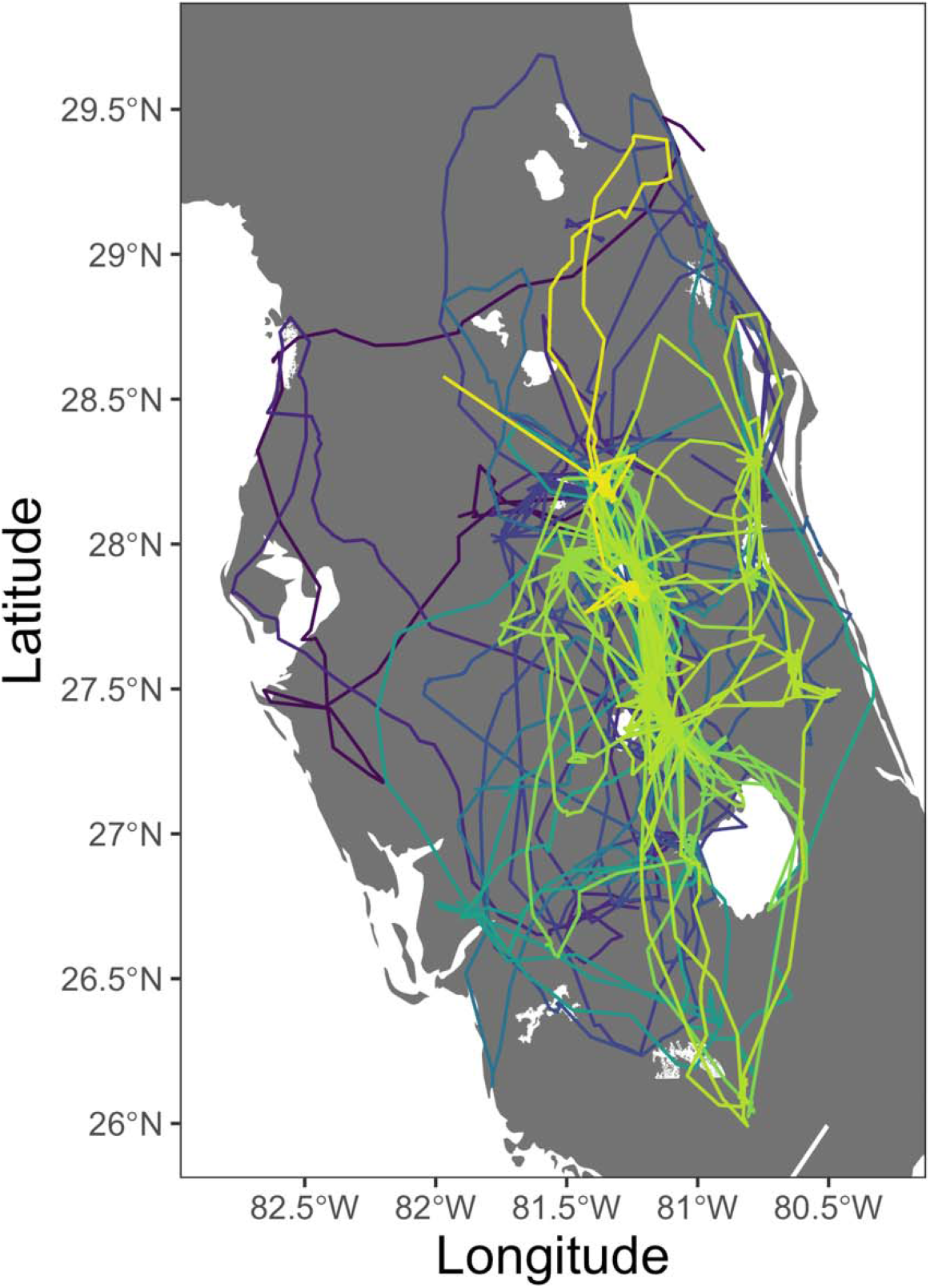
Tracks from 26 GPS-tagged snail kites in central Florida (2018-2019), where colors indicate different individuals.

Step lengths and turning angles were used to estimate latent behavioral states. As was performed on the simulated tracks, step lengths for the empirical data were discretized into five bins using the 25^th^, 50^th^, 75^th^, 90^th^, and 100^th^ quantiles as upper limits. This resulted in bin limits at 0.00, 0.03, 0.09, 0.32, 1.63, and 72.56 km. Turning angles were discretized into eight bins from −*π* to *π* using equal bin widths, resulting in bin limits at 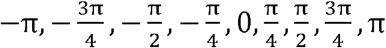.

Step lengths and turning angles for each of the 26 snail kites were analyzed by the M4 segmentation model using 80,000 iterations, a burn-in of 40,000 iterations, and hyperparameter *α* = 1. The MAP estimates of breakpoints for each snail kite were used to define track segments per individual. Subsequently, track segments were analyzed across all individuals via LDA to estimate the most likely number of states, to define state-dependent distributions, and to estimate the proportion of each state that characterized each track segment. This was performed using 1000 MCMC iterations with a burn-in of 500 iterations and vague priors were used with hyperparameters set at γ = 0.1 and α = 0.1 for a maximum possible number of seven states. Trace-plots of the segmentation and LDA models from M4 indicated that all had reached convergence (Appendix S7). Proportions of behavioral states were evaluated over time in relation to emigration from natal sites and peak breeding season of snail kites in Florida (March 1 – June 30; Reichert et al. 2020) to discern any patterns associated with these events.

## 3. Results

### 3.1 Segmentation model comparison

The M4 segmentation model successfully recovered breakpoints from the simulations and outperformed both BCPA and segclust2d. Among the three methods, the segclust2d model took much longer to run (0.46 to 418 min) compared to M4 (1.98 to 227 min) and BCPA models (0.25 to 21 min), particularly for longer tracks (Figs. 4a, 5a). While all three models exhibited similar accuracy on the shortest simulations, M4 was much more accurate on all larger simulations. For these large simulations, accuracy of the M4 segmentation model was >80% on average when simulations were generated from uncommon distributions and >90% on average when generated from common distributions (Figs. 4b, 5b, 6a). Additionally, M4 missed the lowest proportion of true breakpoints (uncommon: 21%; common: 0.3%) compared to BCPA (uncommon: 67%; common: 66%) and segclust2d (uncommon: 26%; common: 30%) across simulations of all analyzed track lengths.

**Figure 4.**
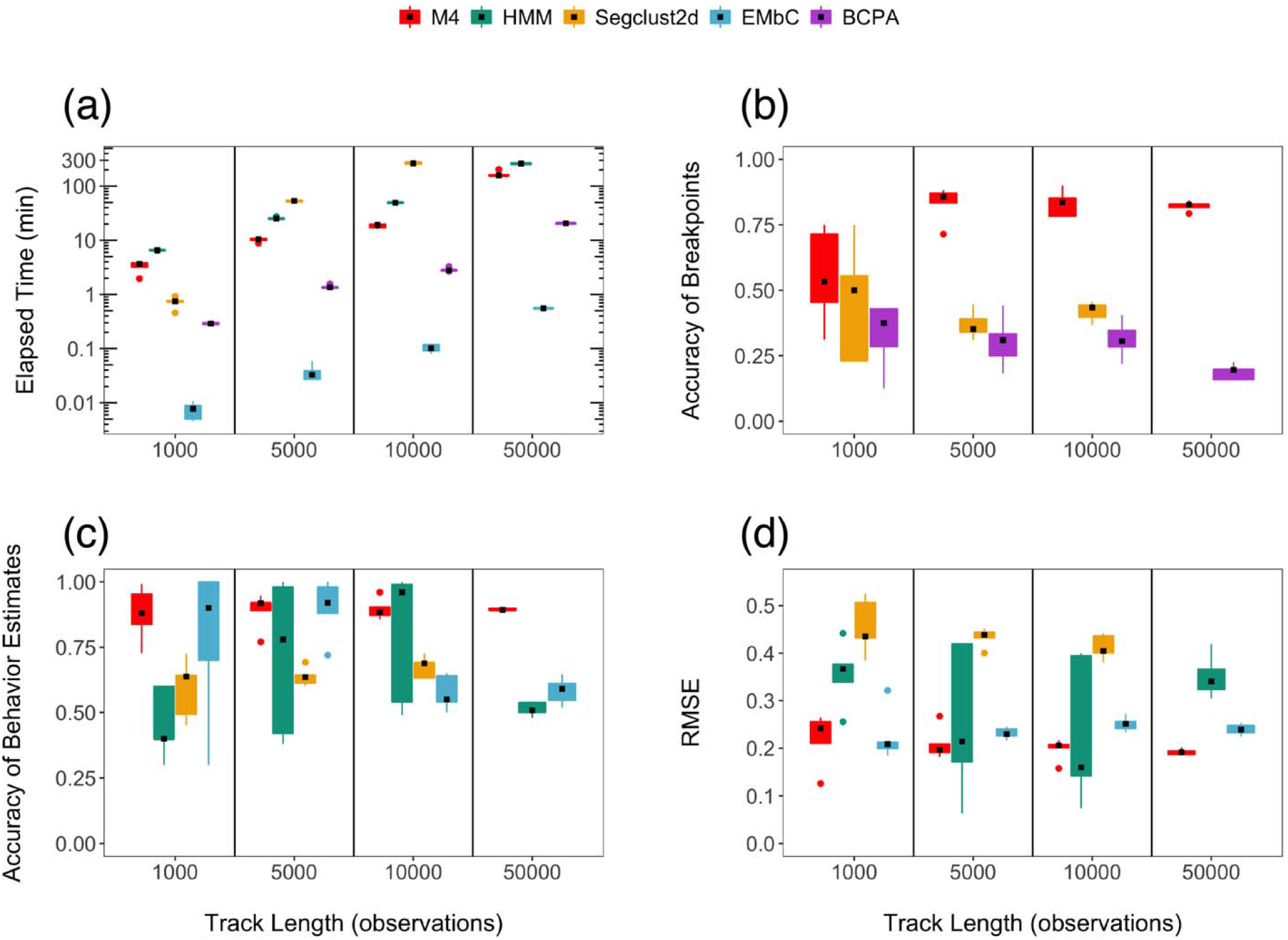
Comparison of the performance of different methods on tracks simulated from uncommon distributions, where black points indicate the median for each boxplot. (a) The elapsed time to analyze each of the simulations is shown for the different methods on a logarithmic scale, where the measure for M4 reflects the sum of elapsed times from both the segmentation and LDA models. (b) Accuracy of breakpoint estimates are compared among the M4, segclust2d, and BCPA models. (c) Accuracy of the estimates for the dominant behavior of each track segment is shown for the M4, HMM, segclust2d, and EMbC models. (d) Accuracy of behavior proportion estimates per track segment are compared among the M4, HMM, segclust2d, and EMbC methods.

**Figure 5.**
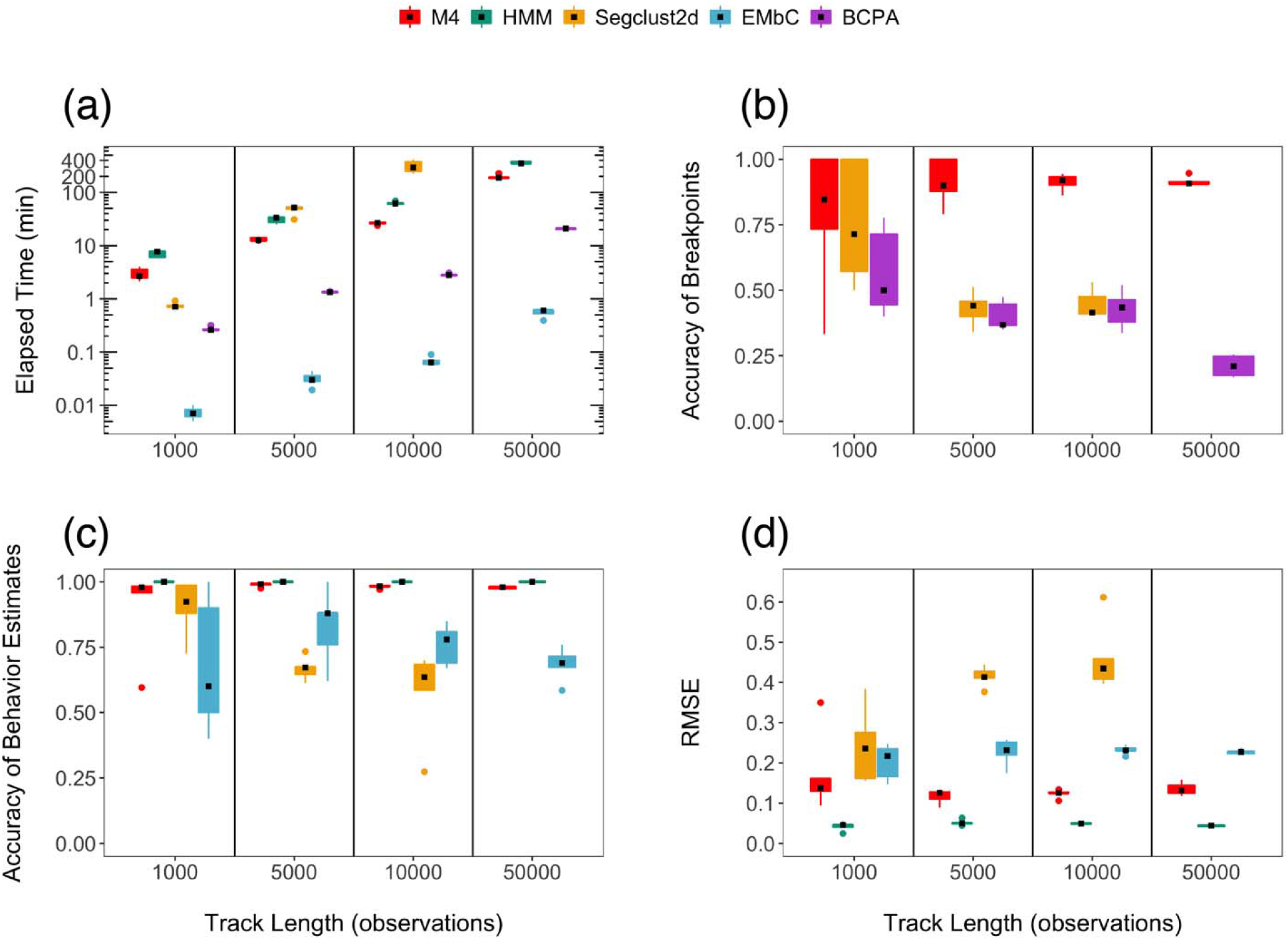
Comparison of the performance of different methods on tracks simulated from common distributions, where black points indicate the median for each boxplot. (a) The elapsed time to analyze each of the simulations is shown for the different methods on a logarithmic scale, where the measure for M4 reflects the sum of elapsed times from both the segmentation and LDA models. (b) Accuracy of breakpoint estimates are compared among the M4, segclust2d, and BCPA models. (c) Accuracy of the estimates for the dominant behavior of each track segment is shown for the M4, HMM, segclust2d, and EMbC models. (d) Accuracy of behavior proportion estimates per track segment are compared among the M4, HMM, segclust2d, and EMbC methods.

**Figure 6.**
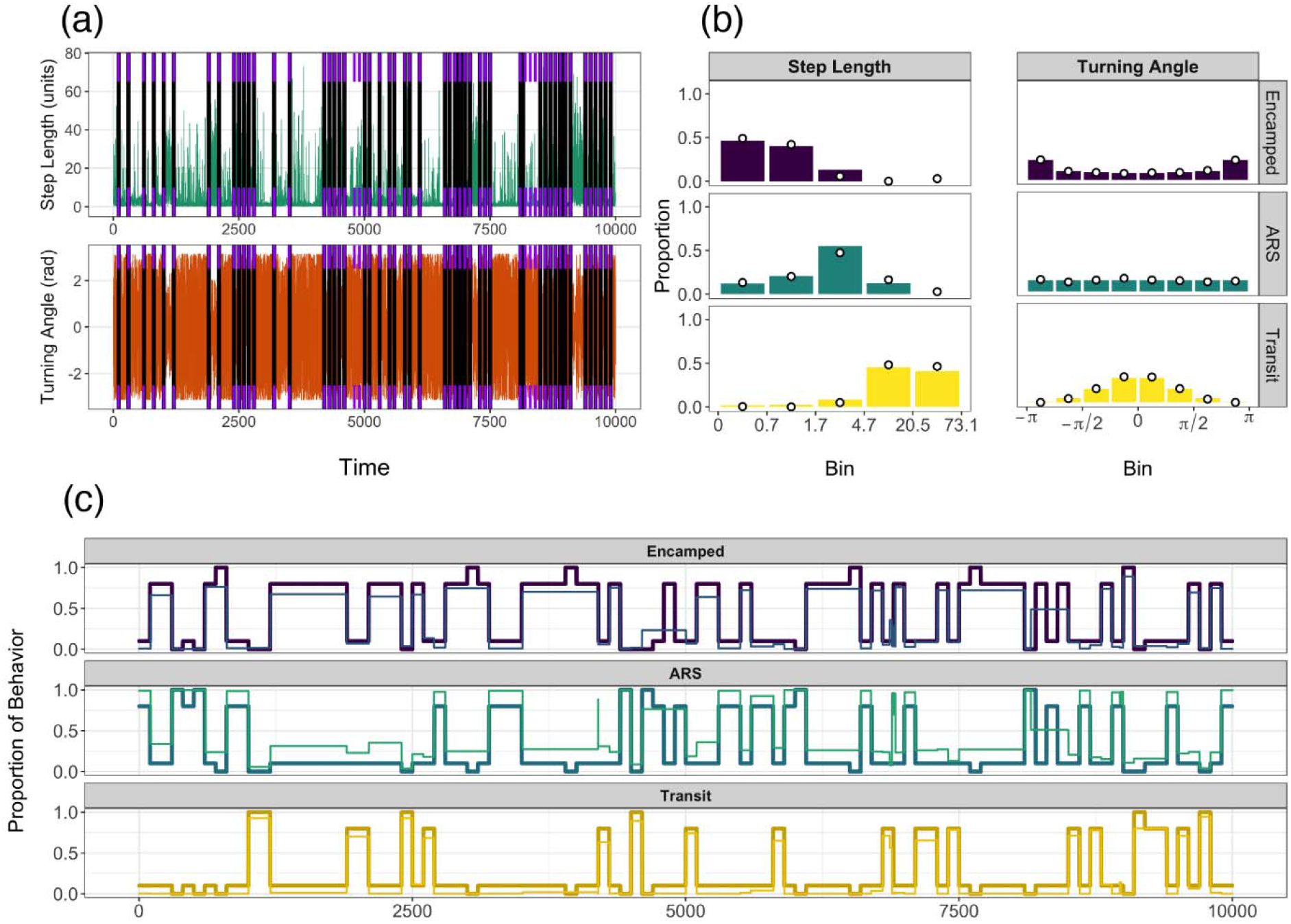
Evaluation of model performance for a single simulation, as an example, for (a) breakpoint estimation, (b) determination of the shape of behavioral states, and (c) estimating behavior proportions over time. (a) A time series of true (purple tick marks) and estimated breakpoints (black lines) are shown for the segmentation of both data streams. (b) Discretized distributions are shown for each data stream per behavioral state where three states were determined to be most likely. Bars denote true proportions for each of the bins whereas points indicate model estimates. (c) Time series of proportions for each behavior are shown; thick, dark lines indicate true behavior proportions whereas thin, light lines indicate estimated proportions of behaviors.

### 3.2 Clustering model comparison

When estimating the true number of states on both sets of simulations, M4 correctly determined the number of true states more frequently than the other methods and exhibited greater computational efficiency over all other clustering methods besides EMbC. The Bayesian LDA model took 2 – 23 s to run, highlighting the computational efficiency of this particular model. When added to the duration of the segmentation model, the proposed method ran faster than the HMM and segclust2d at all track lengths despite these models being fitted with only two to four states whereas our method allows for up to seven states (Figs. 4a, 5a). However, the time to run each EMbC model increased very little with increases in track length, but also automatically assumed four states were present. The LDA model from M4 correctly suggested three states as most likely for 18 of the 20 simulations generated from uncommon distributions and 19 of 20 simulations generated from common distributions (Fig. 6b, Appendix S9). By comparison, the HMM suggested (via AIC and BIC) that three states were most likely in 17 and 16 of the 20 analyzed simulations generated by uncommon and common distributions, respectively. The segclust2d model suggested that three states were most likely in only 6 and 5 of the 15 analyzed simulations generated from uncommon and common distributions, respectively, based on BIC.

To enable direct comparisons among all four models that estimated behavioral states, we assumed three states were most likely for all 20 simulations when calculating model accuracy. Using this assumption, we find that M4 demonstrated high accuracy in behavioral state estimation for both sets of simulations, often equivalent or superior to the other clustering methods. When analyzing simulations generated from uncommon distributions, mean accuracy of M4 to classify the dominant state within each segment was greater than that of the HMM and segclust2d models at all track lengths (Fig. 4c). However, mean accuracy of the EMbC model was slightly greater than M4 on this set of simulations at a track length of 5000 observations. When analyzing simulations generated from common distributions, mean accuracy of M4 was slightly below that of the HMM, but greater than the mean accuracy of the EMbC and segclust2d models at all track lengths (Fig. 5c). Additionally, accuracy measures displayed little variability in M4 across tracks of different lengths and on each set of simulations, highlighting the increased stability of this framework. Similar to the pattern found for estimates of dominant behavioral states, accuracy of behavioral state proportions were higher in M4 for all but the HMM on simulations generated from common distributions, as denoted by low RMSE (Figs. 4d, 5d, 6c).

The accuracy of the estimated step lengths and turning angles distributions were relatively consistent across each set of simulations. For tracks generated from uncommon distributions, M4 was slightly more accurate than the HMM, but much more accurate than EMbC and segclust2d across all track lengths (Appendix S9). However, HMM estimates were slightly more accurate than the Bayesian model on tracks of all lengths when generated from common distributions (Appendix S9). When viewed as continuous distributions, it is clear that the HMM, EMbC, and segclust2d models had difficulty estimating the true distributions of step lengths and turning angles regardless of track length for the simulation with uncommon distributions (Appendix S10). On the other hand, the HMM was able to perfectly estimate the state-dependent distributions of the simulations generated from common distributions (Appendix S10).

### 3.3 Snail kite analysis

The segmentation of 26 snail kite trajectories using M4 took approximately 61 min to run and estimated 1 to 64 breakpoints for these individuals. Breakpoints were then used to define 444 track segments from all individuals (Fig. 7a). These segments were clustered into states using M4, which took approximately 27 s to run. It appeared that there were likely three behavioral states, which comprised 91.6% of all state assignments on average (Fig. 7b). To ensure that these three states were biologically interpretable, distributions of step lengths and turning angles were also evaluated (Fig. 7c). The distributions showed: 1) a slow and tortuous behavior; 2) a tortuous behavior with intermediate speed; and 3) a fast and directed behavior. For this reason, these behaviors were labelled ‘encamped, ‘ARS’, and ‘transit’, respectively.

**Figure 7.**
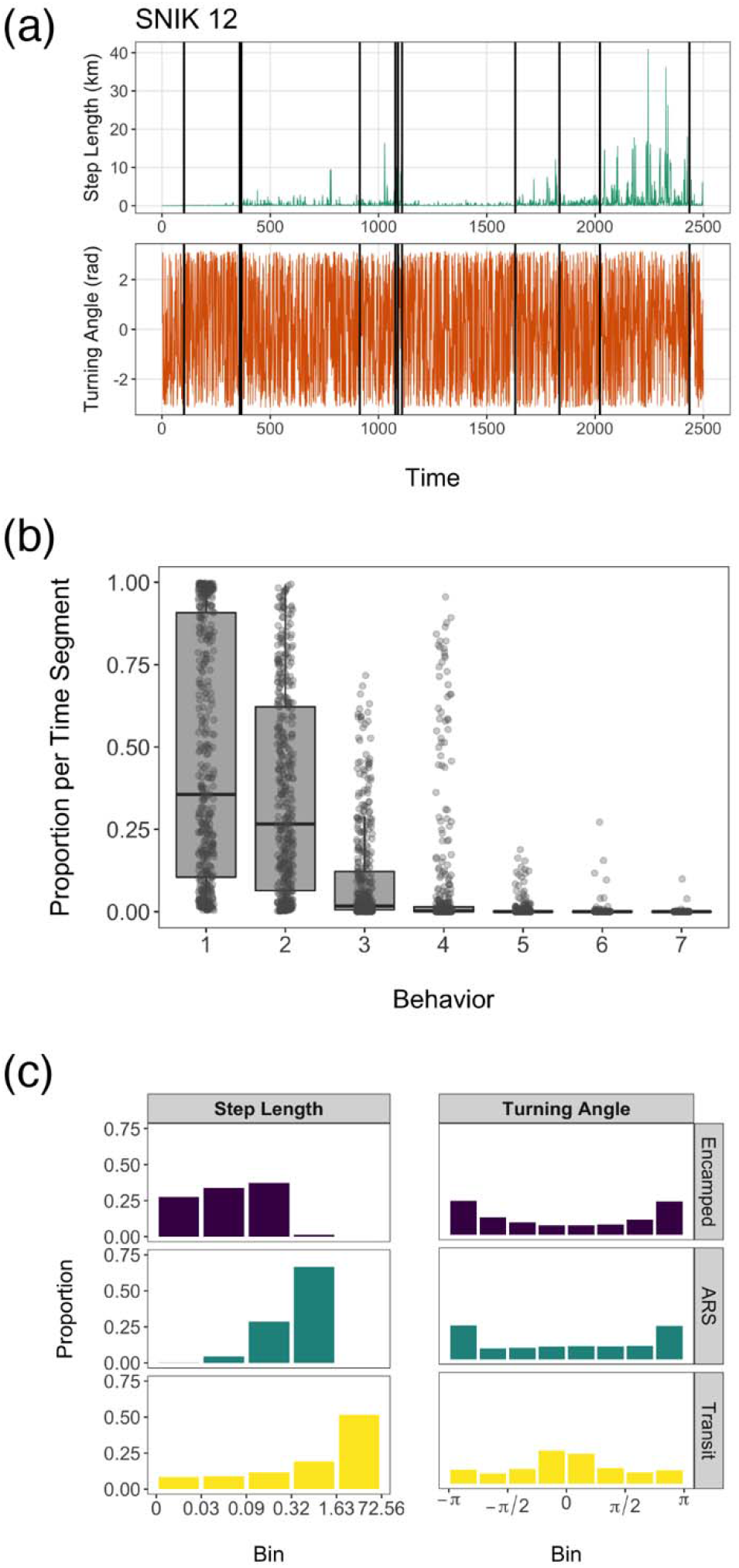
Snail kite results from track segmentation and determination of the most likely number of behavioral states. (a) A subset of a time series from a single individual (SNIK 12) that displays estimated breakpoints (black lines) overlaying each data stream. Time series of step lengths and turning angles are shown on a continuous scale in relation to estimated breakpoints for ease of interpretation. (b) Boxplot showing the estimated proportions for each of the seven possible behaviors from all 444 track segments analyzed. (c) Distributions of step lengths (km) and turning angles (rad) from each of the three retained behavioral states, ordered from slowest to fastest.

Some individuals were only tracked for a short period of time and did not leave the natal area. However, 17 birds did emigrate from their natal site. Dispersal events were typically denoted by a brief period of ARS or transit behavior (Figs. 8a, b, Appendix S9). The three longest tracks, which belonged to snail kites tracked for more than a year (SNIK 12, SNIK 14, and SNIK 15), displayed relatively synchronous behavior before, during, and after their first breeding season. Two brief periods of high activity behavior that occurred during and immediately following peak breeding season in 2019 may potentially represent pre- and post-breeding dispersal events (Figs. 8a, c, d).

**Figure 8.**
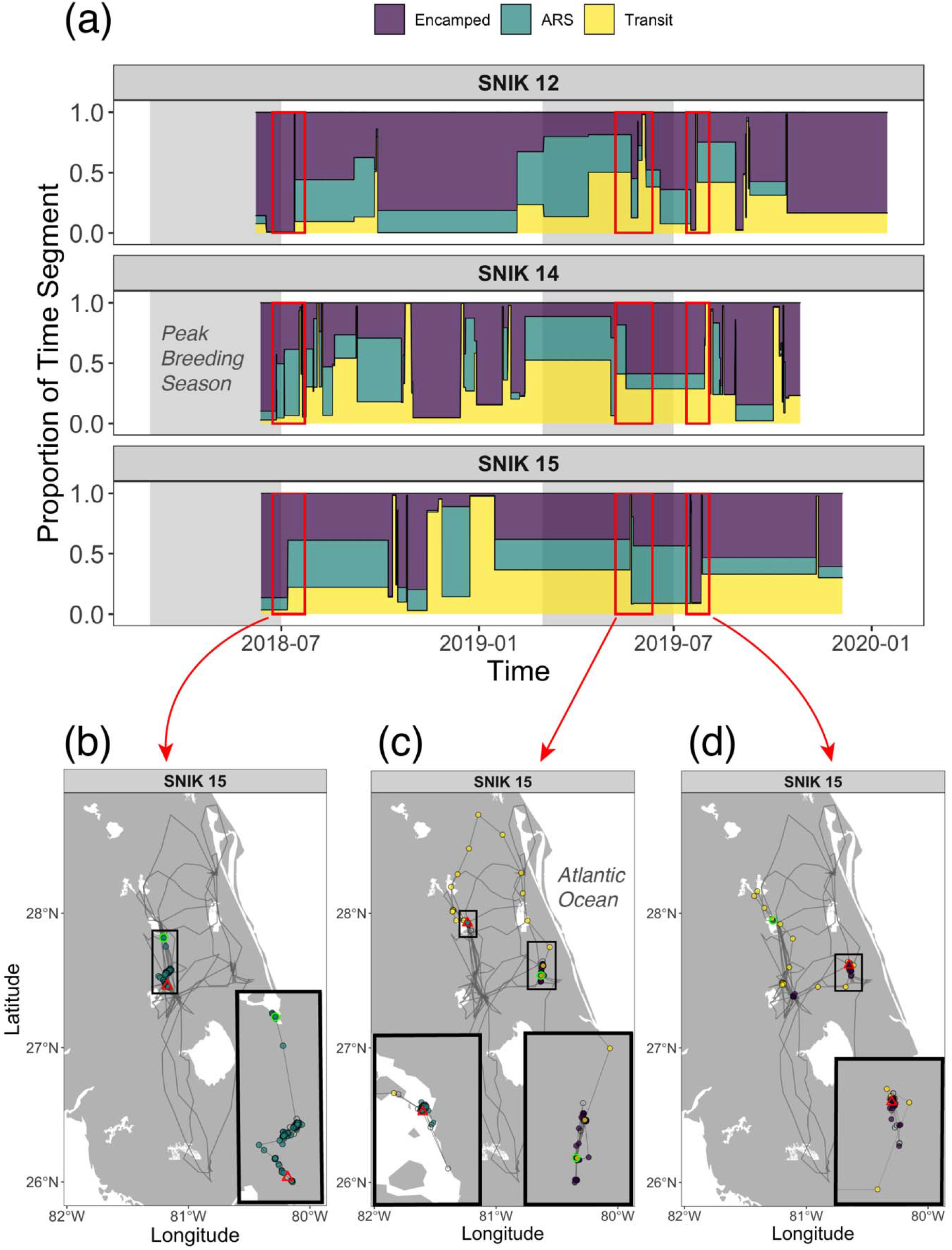
Patterns of movement behavior are shown over time for the three snail kites that were tagged over the longest durations, with particular focus on SNIK 15. (a) Time series of behavior proportions are shown with respect to peak breeding season (grey panels) for each of the three snail kites, with possible natal, pre-breeding, and post-breeding dispersal events denoted by red boxes. Maps of SNIK 15 movement depict the starting (green circle) and ending locations (red triangle) for (b) natal, (c) pre-breeding, and (d) post-breeding season dispersal events, as well as the dominant behavior associated with each track segment. Observations without behavior estimates (i.e., observations not recorded at 1-h time intervals) are shown as open points.

## 4. Discussion

We demonstrated that our Bayesian M4 framework (available within the bayesmove R package) can accurately identify changes in behavioral states, reliably estimate the most likely number of behavioral states, and properly characterize the state-dependent distributions of data streams. This two-stage model treats track segments as the unit of interest (as opposed to observations) and relies on the discretization of data streams to avoid the need to specify parametric probability distributions. Importantly, the proposed method is relatively computationally efficient, a key characteristic given the ever-increasing storage capacities of modern sensors and their ability to measure a growing number of intrinsic and environmental variables (Whitford and Klimley 2019; Williams et al. 2020). A comparison of model performance in addition to the analysis of an empirical dataset highlight the utility of the M4 framework.

### 4.1 Method Comparison

Although BCPA was notably faster at track segmentation than M4, accuracy of the estimated breakpoints was much higher in the latter. Additionally, M4 was faster and exhibited greater accuracy of breakpoint estimates than the segclust2d method, which was not able to successfully analyze simulated tracks of 50000 observations. Since the accuracy of the segclust2d method was not much greater than the BCPA for either set of simulations (Figs. 4b, 5b), it appears that BCPA’s reliance on a single derived variable (i.e., persistence velocity) instead of separate data streams was not as limiting as was initially expected.

While HMMs are powerful methods that can incorporate individual-level random effects and account for cyclical patterns (Patterson et al. 2017; McClintock and Michelot 2018), they can also be restrictive in some of their assumptions. Standard forms of HMMs require the use of parametric distributions, which may not fit the data streams well (Appendix S9; Langrock et al. 2018). While HMMs displayed better performance than M4 when the selected parametric distributions matched the true underlying distributions (Fig. 5c), we find that the proposed methodology performed better than HMMs when the selected parametric distributions did not match the true underlying distribution. By comparison, the segclust2d and EMbC methods are straightforward to apply when estimating latent behavioral states from a set of tracks, but appear limited by their assumption of Gaussian distributions when partitioning observations into segments or into states, respectively. Since the most common data streams (i.e., step lengths and turning angles) are not typically modeled with a Gaussian distribution (McClintock et al. 2020), this likely contributes to the lower accuracy of these models.

The determination of the most likely number of states is another issue when fitting clustering models and HMMs since this is typically unknown *a priori* and is directly impacted by how well the selected parametric distributions characterize the states (Pohle et al. 2017). Unfortunately, HMMs often require multiple models to be fit and compared using information theoretic approaches, which tend to favor a greater number of states than are truly present and come at a high computational cost (Li and Bolker 2017; Pohle et al. 2017). Importantly, while M4 allows for up to 7 behavioral states, we only attempted to fit HMMs with 2 to 4 behavioral states. Even in this limited context, fitting HMMs was already much slower than fitting M4. Had we attempted to fit HMMs with 2 to 7 behavioral states, the amount of time required for this would be substantially larger than what we report in Figs. 4 and 5. A similar issue is present in segclust2d, where models are fit with every possible number of track segments and states before comparing via BIC. A different problem is posed by the EMbC model, which imposes four states by default when analyzing step lengths and turning angles. These issues are directly addressed by our framework since we use a mixed-membership model (LDA) with a penalizing prior to cluster track segments, enabling the estimation of the most likely number of states and the state-dependent distributions in a single step. While existing methods can provide useful behavioral inference depending upon the ecological question and dataset, the M4 framework provides a powerful alternative when behaviors are complex, multiple data streams are available and these data are not well characterized by parametric distributions, and/or datasets are large.

### 4.2 Empirical applications

Three behavioral states were clearly estimated for the snail kite dataset, which was supported by biologically relevant distributions of step lengths and turning angles. The ‘encamped’ state likely represents fine-scale behaviors that include resting, feeding, and time spent at the nest (as a fledgling or reproductive adult). On the other hand, the ‘ARS’ state likely includes exploration for nearby suitable habitat as well as foraging bouts (Martin et al. 2006; Pias et al. 2016). Finally, the ‘transit’ state includes fast, directed movements associated with dispersal of snail kites in addition to departure from wetlands experiencing low water levels (Robertson et al. 2017).

The time series of snail kite behavior proportions showed variability in the timing of emigration from natal sites among individuals, but changes in behavior were generally synchronous in the three birds that reached maturity. This variability in the timing of emigration from natal sites could be due to a variety of factors, such as hatching date, body condition, and local environmental conditions (Rodgers and Schwikert 2003; Fletcher et al. 2015; Cattau et al. 2016). The shifts in behavior proportions appeared to show multiple phases of high and low activity, some of which seem to match the phenology of natal dispersal (summer), pre-breeding dispersal (early spring), and post-breeding dispersal (late summer) (Bennetts and Kitchens 2000). While the continued monitoring of these tagged birds should provide greater evidence for the characterization of activity budgets over ontogeny, future research could also explore the primary drivers of snail kite movement and habitat use within each behavioral state through the inclusion of environmental covariates.

### 4.3 Caveats and extensions

In addition to the M4 method proposed by this study, other non-parametric state estimation methods have been previously developed (Nams 2014; Sur et al. 2014; Langrock et al. 2018). In one such example, the behavioral movement segmentation (BMS) model proposed by Nams (2014) uses a combination of direct search optimization, iterative sampling, and k-means clustering to estimate latent states from track segments. BMS is similar to our proposed M4 framework in that both methods are non-parametric, partition multiple data streams into segments, and cluster segments into latent states (Nams 2014). However, M4 differs both technically and conceptually from BMS in that M4 proposes breakpoints using RJMCMC, the number of likely states are estimated within a single model run (instead of using multi-model selection), and track segments are expected to be comprised of multiple states rather than just one. We believe that practitioners should carefully evaluate the properties and assumptions of different methods to determine the best method to properly analyze their data and address their objectives.

Although M4 effectively classified behavioral states from both simulated and empirical tracks, there are some limitations to this approach. The selection of the number and width of bins when discretizing data streams is a subjective choice that impacts the results from the segmentation model and ultimately the estimation of behavioral states. Additionally, our model implicitly assumes that location error is negligible or requires that it be accounted for via another method. Although our model can analyze data streams from regular or irregular time intervals, this will also depend on the inherent properties of the data streams themselves. Since step lengths and turning angles are calculated from multiple successive observations, these values will not be comparable once the data are not close to a regular time interval. However, variables such as net squared displacement (the squared distance from the starting location to all other relocations) can be analyzed over irregular time intervals.

M4 can be extended to analyze other types of data streams and can include prior knowledge on the timing of behavioral shifts. Although only step lengths and turning angles were analyzed for the simulated and empirical tracks, additional ancillary data coming from the sensor (e.g., elevation, salinity, temperature, or accelerometer data) could be used to make behavioral inference. These data streams could come from all types of distributions (i.e., continuous, discrete, bounded between 0 and 1). It is also relatively straightforward to deal with zero-inflated data by including all zeroes in a single bin. Additionally, our segmentation model can be implemented in a semi-supervised fashion, by which practitioners pre-specify breakpoints for the time series based on *a priori* knowledge and these breakpoints will be considered by the RJMCMC algorithm. This may be particularly useful if daily activity patterns are expected or if only one of several possible states can be clearly identified.

## Acknowledgments

This work was supported in part by the University of Florida Biodiversity Institute (UFBI), the US Department of Agriculture’s National Institute of Food and Agriculture (USDA-NIFA) McIntire–Stennis project 1005163, the US National Science Foundation (NSF) award 1458034 and 2040819, the Florida Fish and Wildlife Conservation Commission (FWC), and by the US Army Corp of Engineers (USACE). The tagging of snail kites was approved and conducted under UF IACUC no. 201708334. JAC was supported by a postdoctoral fellowship from the University of Florida Informatics Institute (UFII) Fellowship Program. We would like to thank Tyler Beck for his assistance with field work and Brenda Betancourt for her feedback during model and package development. Additionally, we are grateful to Rocío Joo and Mathieu Basille for their valuable comments on an earlier draft of this paper.

## Authors’ Contributions

J.A.C. and D.V. conceived the ideas and designed methodology; C.L.P. and R.J.F. collected the empirical data; J.A.C., C.L.P., and D.V. analyzed the data; J.A.C. and D.V. led the writing of the manuscript. All authors contributed critically to the drafts and gave final approval for publication.

## Data Availability

All code mentioned here for the Bayesian framework is available within the bayesmove package for R hosted on CRAN at https://CRAN.R-project.org/package=bayesmove. The development version of the package is available on GitHub at https://github.com/joshcullen/bayesmove and a full set of vignettes can be found at https://joshcullen.github.io/bayesmove. The code to generate the simulations and perform method comparison are available on Zenodo (https://doi.org/10.5281/zenodo.4898208). The Everglade snail kite telemetry data have not been made available since this is a federally listed endangered species and the location data are sensitive.

## Notes

https://CRAN.R-project.org/package=bayesmove

https://joshcullen.github.io/bayesmove

https://doi.org/10.5281/zenodo.4898208

### Competing Interest Statement

The authors have declared no competing interest.

### Summary of Updates

Revisions have been made in response to comments made during peer-review. This includes updates to the manuscript and the addition of supplemental files.

## References

Bennetts RE, Kitchens WM. 2000. Factors influencing movement probabilities of a nomadic food specialist: proximate foraging benefits or ultimate gains from exploration? Oikos 91:459–467. doi:10.1034/j.1600-0706.2000.910306.x.

Burnham KP, Anderson DR. 2002. Model selection and multimodel inference: A practical information-theoretic approach. 2nd ed. New York: Springer.

Cattau CE, Fletcher RJ, Reichert BE, Kitchens WM. 2016. Counteracting effects of a non-native prey on the demography of a native predator culminate in positive population growth. Ecological Applications 26:1952–1968. doi:10.1890/15-1020.1/suppinfo.

Cullen JA, Valle D. 2021. bayesmove: Non-Parametric Bayesian Analysis of Animal Movement. R Package version 0.2.0.

Denison DGT, Holmes CC, Mallick BK, Smith AFM. 2002. Bayesian methods for nonlinear classification and regression. Chichester, UK: John Wiley & Sons.

Edelhoff H, Signer J, Balkenhol N. 2016. Path segmentation for beginners: an overview of current methods for detecting changes in animal movement patterns. Movement Ecology 4:21. doi:10.1186/s40462-016-0086-5.

Fletcher RJ, Robertson EP, Wilcox RC, Reichert BE, Austin JD, Kitchens WM. 2015. Affinity for natal environments by dispersers impacts reproduction and explains geographical structure of a highly mobile bird. Proceedings of the Royal Society B: Biological Sciences 282:20151545. doi:10.1098/rspb.2015.1545.

Fraser KC, Davies KTA, Davy CM, Ford AT, Flockhart DTT, Martins EG. 2018. Tracking the conservation promise of movement ecology. Frontiers in Ecology and Evolution 6:150. doi:10.3389/fevo.2018.00150.

Garriga J, Palmer JRB, Oltra A, Bartumeus F. 2016. Expectation-maximization binary clustering for behavioural annotation. PLoS ONE 11:1–26. doi:10.1371/journal.pone.0151984.

Garriga J, Palmer JRB, Oltra A, Bartumeus F. 2019. EMbC: Expectation-Maximization Binary Clustering. R package version 2.0.3.

Green PJ. 1995. Reversible jump Markov chain Monte Carlo computation and Bayesian model determination. Biometrika 82:711–732. doi:10.2307/2337340.

Gurarie E. 2014. bcpa: Behavioral change point analysis of animal movement. R package version 1.1.

Gurarie E, Andrews RD, Laidre KL. 2009. A novel method for identifying behavioural changes in animal movement data. Ecology Letters 12:395–408. doi:10.1111/j.1461-0248.2009.01293.x.

Gurarie E, Bracis C, Delgado M, Meckley TD, Kojola I, Wagner CM. 2016. What is the animal doing? Tools for exploring behavioural structure in animal movements. Journal of Animal Ecology 85:69–84. doi:10.1111/1365-2656.12379.

Hudon SF, Zaiats A, Roser A, Roopsind A, Barber C, Robb BC, Pendleton BA, Camp MJ, Clark PE, Davidson MM, et al. 2021. Unifying community detection across scales from genomes to landscapes. Oikos. doi:10.1111/oik.08393.

Hussey NE, Kessel ST, Aarestrup K, Cooke SJ, Cowley PD, Fisk AT, Harcourt RG, Holland KN, Iverson SJ, Kocik JF, et al. 2015. Aquatic animal telemetry: A panoramic window into the underwater world. Science 348:1255642. doi:10.1126/science.1255642.

John GH, Langley P. 1995. Estimating continuous distributions in Bayesian classifiers. In: Proceedings of the Eleventh Conference on Uncertainty in Artificial Intelligence. Morgan Kaufmann Publishers Inc. p. 338–345.

Jonsen I. 2016. Joint estimation over multiple individuals improves behavioural state inference from animal movement data. Scientific Reports 6:20625. doi:10.1038/srep20625.

Jonsen ID, McMahon CR, Patterson TA, Auger-Méthé M, Harcourt R, Hindell MA, Bestley S. 2019. Movement responses to environment: fast inference of variation among southern elephant seals with a mixed effects model. Ecology 100:e02566. doi:10.1002/ecy.2566.

Joo R, Boone ME, Clay TA, Patrick SC, Clusella-Trullas S, Basille M. 2020. Navigating through the r packages for movement. Journal of Animal Ecology 89:248–267. doi:10.1111/1365-2656.13116.

Joo R, Picardi S, Boone ME, Clay TA, Patrick SC, Romero-Romero VS, Basille M. 2020. A decade of movement ecology. arXiv preprint:2006.00110.

Kays R, Crofoot MC, Jetz W, Wikelski M. 2015. Terrestrial animal tracking as an eye on life and planet. Science 348:aaa2478. doi:10.1126/science.aaa2478.

Kitagawa G. 1987. Non-gaussian state-space modeling of nonstationary time series. Journal of the American Statistical Association 82:1032–1041. doi:10.1080/01621459.1987.10478534.

Langrock R, Adam T, Leos-Barajas V, Mews S, Miller DL, Papastamatiou YP. 2018. Spline-based nonparametric inference in general state-switching models. Statistica Neerlandica 72:179–200. doi:10.1111/stan.12133.

Li M, Bolker BM. 2017. Incorporating periodic variability in hidden Markov models for animal movement. Movement Ecology 5:1. doi:10.1186/s40462-016-0093-6.

Martin J, Nichols JD, Kitchens WM, Hines JE. 2006. Multiscale patterns of movement in fragmented landscapes and consequences on demography of the snail kite in Florida.:527–539. doi:10.1111/j.1365-2656.2006.01073.x.

McClintock BT, Langrock R, Gimenez O, Cam E, Borchers DL, Glennie R, Patterson TA. 2020. Uncovering ecological state dynamics with hidden Markov models. Ecology Letters.doi:10.1111/ele.13610.

McClintock BT, Michelot T. 2018. momentuHMM: R package for generalized hidden Markov models of animal movement. Methods in Ecology and Evolution 9:1518–1530. doi:10.1111/2041-210X.12995.

Michelot T, Langrock R, Patterson TA. 2016. moveHMM: an R package for the statistical modelling of animal movement data using hidden Markov models. Methods in Ecology and Evolution 7:1308–1315. doi:10.1111/2041-210X.12578.

Nams VO. 2014. Combining animal movements and behavioural data to detect behavioural states. Ecology Letters 17:1228–1237. doi:10.1111/ele.12328.

Nathan R, Getz WM, Holyoak M, Kadmon R, Saltz D, Smouse PE. 2008. A movement ecology paradigm for unifying organismal movement research. Proceedings of the National Academy of Sciences 105:19052–19059. doi:10.1021/i360006a005.

Patin R, Etienne M-P, Lebarbier E, Benhamou S. 2019. segclust2d: Bivariate Segmentation/Clustering Methods and Tools. R package version 0.2.0.

Patin R, Etienne MP, Lebarbier E, Chamaillé-Jammes S, Benhamou S. 2020. Identifying stationary phases in multivariate time series for highlighting behavioural modes and home range settlements. Journal of Animal Ecology 89:44–56. doi:10.1111/1365-2656.13105.

Patterson TA, Parton A, Langrock R, Blackwell PG, Thomas L, King R. 2017. Statistical modelling of individual animal movement: an overview of key methods and a discussion of practical challenges. AStA Advances in Statistical Analysis 101:399–438. doi:10.1007/s10182-017-0302-7.

Pias KE, Fletcher RJ, Kitchens WM. 2016. Assessing the value of novel habitats to snail kites through foraging behavior and nest survival. Journal of Fish and Wildlife Management 7:449–460. doi:10.3996/022016-JFWM-008.

Pohle J, Langrock R, van Beest FM, Schmidt NM. 2017. Selecting the number of states in hidden Markov models: Pragmatic solutions illustrated using animal movement. Journal of Agricultural, Biological, and Environmental Statistics 22:270–293. doi:10.1007/s13253-017-0283-8.

Potts JR, Börger L, Scantlebury DM, Bennett NC, Alagaili A, Wilson RP. 2018. Finding turning-points in ultra-high-resolution animal movement data. Methods in Ecology and Evolution 9:2091–2101. doi:10.1111/2041-210X.13056.

Reichert B, Cattau C, Fletcher Jr. R, Sykes Jr. P, Rodgers Jr. J, Bennetts R. 2020. Snail Kite (Rostrhamus sociabilis), version 1.0. In: Poole A, editor. Birds of the World. Ithaca, NY, USA: Cornell Lab of Ornithology.

Robertson EP, Fletcher RJ, Austin JD. 2017. The causes of dispersal and the cost of carry-over effects for an endangered bird in a dynamic wetland landscape. Journal of Animal Ecology 86:857–865. doi:10.1111/1365-2656.12676.

Rodgers JA, Schwikert ST. 2003. Breeding chronology of snail kites (Rostrhamus sociabilis plumbeus) in central and south Florida wetlands. Southeastern Naturalist 2:293–300. doi: 10.1656/1528-7092(2003)002[0293:bcoskr]2.0.co;2.

Sur M, Skidmore AK, Exo KM, Wang T, Ens BJ, Toxopeus AG. 2014. Change detection in animal movement using discrete wavelet analysis. Ecological Informatics 20:47–57. doi:10.1016/j.ecoinf.2014.01.007.

Valle D, Baiser B, Woodall CW, Chazdon R. 2014. Decomposing biodiversity data using the Latent Dirichlet Allocation model, a probabilistic multivariate statistical method. Ecology Letters 17:1591–1601. doi:10.1111/ele.12380.

Valle D, Cvetojevic S, Robertson EP, Reichert BE, Hochmair HH, Fletcher RJ. 2017. Individual movement strategies revealed through novel clustering of emergent movement patterns. Scientific Reports 7:44052. doi:10.1038/srep44052.

Whitford M, Klimley AP. 2019. An overview of behavioral, physiological, and environmental sensors used in animal biotelemetry and biologging studies. Animal Biotelemetry 7:26. doi:10.1186/s40317-019-0189-z.

Williams HJ, Taylor LA, Benhamou S, Bijleveld AI, Clay TA, de Grissac S, Demšar U, English HM, Franconi N, Gómez-Laich A, et al. 2020. Optimizing the use of biologgers for movement ecology research. Journal of Animal Ecology 89:186–206. doi:10.1111/1365-2656.13094.

