## Appendix S1 for "Identifying latent behavioral states in animal movement with M4, a non-parametric Bayesian method"

Appendix S1: Sensitivity Analysis of Binning Method and Thresholds of Breakpoint Accuracy

Identifying latent behavioral states in animal movement with M4, a non-parametric Bayesian method

Joshua A Cullen^1*^, Caroline L Poli^2^, Robert J Fletcher, Jr.^3^, Denis Valle^1^

^1^ School of Forest Resources and Conservation, University of Florida, Gainesville, FL, USA

^2^ School of Natural Resources and Environment, University of Florida, Gainesville, FL, USA

^3^ Department of Wildlife Ecology and Conservation, University of Florida, Gainesville, FL, USA

1. Sensitivity of binning method

The process of selecting a method to discretize the data streams is relatively subjective, but can have a large impact on the results of the non-parametric Bayesian analysis (M4). While the use of equal bin widths for data streams are sensible for distributions that are relatively uniform or are inherently circular (i.e., turning angles), this is not necessarily the case for distributions that are highly skewed (e.g., step lengths). It is possible that the use of quantiles may be better to characterize the overall distribution. Additionally, the number of bins used to discretize the data streams is also expected to influence the results from the segmentation and clustering steps of M4, where a greater number of bins may better characterize the entire distribution. Therefore, we conducted a sensitivity analysis varying the discretization method as well as the number of bins used on the analysis of 10 simulated tracks with lengths of 5000 or 10000 observations and that were generated using uncommon distributions for step lengths (i.e., truncated normal) and turning angles (i.e., beta, uniform, and truncated normal).

This sensitivity analysis compared the accuracy of estimated breakpoints, the number of missed breakpoints, as well as the number of likely behavioral states among four different scenarios. While turning angles were discretized into 8 bins of equal widths from -π to π radians, step lengths were discretized into either 5 or 10 bins using either equal widths or quantiles. When discretizing step lengths into bins using the ‘equal widths’ method, all but the last bin were of equal widths since it was more important to characterize the bulk of the distribution density compared to the tail. Specifically, we set the beginning of the last bin to the 95^th^ quantile of step lengths so that all extreme values were captured in a single bin. This resulted either 4 or 9 equal-sized bins between 0 and 30 (the 95^th^ quantile) (Fig S1.1). When discretizing step lengths into 5 bins of equal widths, bin limits of 0, 7.5, 15, 22.5, 30, and 108 were used. Likewise, the use of 10 bins and equal widths resulted in bin limits of 0, 3.33, 6.67, 10, 13.33, 16.67, 20, 23.33, 26.67, 30, and 108. When discretizing with quantiles for 5 bins, the 0^th^, 25^th^, 50^th^, 75^th^, 90^th^, and 100^th^ quantiles were selected, which resulted in bin limits of 0, 0.72, 1.66, 4.49, 20.05, and 108. When discretizing with quantiles for 10 bins, the 0^th^, 10^th^, 20^th^, 30^th^, 40^th^, 50^th^, 60^th^, 70^th^, 80^th^, 90^th^, and 100^th^ quantiles were selected, which resulted in bin limits of 0, 0.29, 0.57, 0.87, 1.22, 1.66, 2.35, 3.54, 6.47, 20.05, and 108.

Results slightly differed across tracks of 5000 observations (N=5) and 10000 observations (N=5) in length, but both suggested that the use of quantiles resulted in better discrimination of the true simulated states (Tables S1.1). The use of equal bin widths exhibited greater accuracy for breakpoint estimates (87 – 94% on average), but also displayed a 3-5 fold greater number of true breakpoints that were missed. This high proportion of missing breakpoints for the ‘equal bin width’ method likely contributed to the incorrect suggestion of 2 latent behavioral states when 3 were simulated (Tables S1.1). By comparison, the use of quantiles resulted in slightly lower accuracy of breakpoint estimates (80 – 83% on average), but much lower rates of breakpoints that were missed (13 – 21% on average). Additionally, all but one of the simulated tracks discretized using quantiles were able to correctly estimate 3 latent behavioral states from the segmented tracks. There did not appear to be a difference in the results of step lengths discretized with quantiles using 5 bins or 10 bins, so the use of fewer bins will likely be more interpretable and will result in faster model run times. Therefore, we suggest the use of quantiles when discretizing highly skewed distributions for analysis by this non-parametric Bayesian M4 framework.

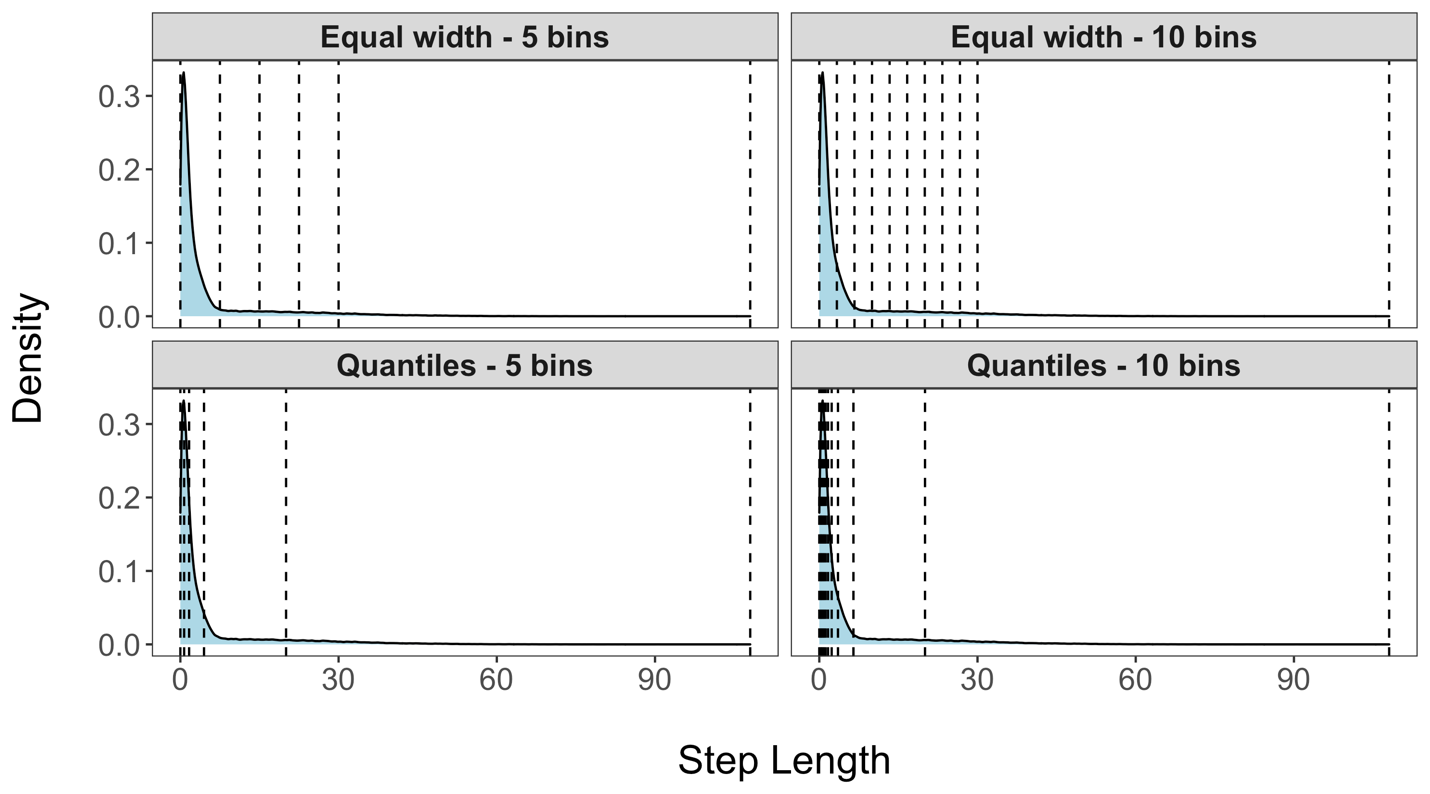

Figure S1.1 Bin limits are shown overlaying a density plot of step lengths for each of the four different methods that were compared for the sensitivity analysis.

Table S1.1. Results of sensitivity analysis on binning method for simulated tracks of 5,000 and 10,000 observations. The mean (min-max) percentage accuracy of breakpoint estimation and the mean (min-max) percentage of missed breakpoints are calculated for each method. Based on clustering of segmented tracks from the LDA model, the mode (% models matching the mode) of the likely number of latent states is also shown.

| Track Length | Method | % Accurate Breakpoints | % Missed Breakpoints | Estimated Number of States |
| --- | --- | --- | --- | --- |
| 5,000 | Equal width – 5 bins | 94.0 (88.2 – 100.0) | 62.0 (52.9 – 74.2) | 2 (100.0) |
|  | Equal width – 10 bins | 93.3 (81.8 – 100.0) | 62.2 (50.0 – 74.2) | 2 (100.0) |
|  | Quantiles – 5 bins | 83.2 (71.4 – 88.2) | 16.3 (10.0 – 22.6) | 3 (100.0) |
|  | Quantiles – 10 bins | 79.5 (70.5 – 88.9) | 13.1 (6.5 – 20.0) | 3 (100.0) |
| 10,000 | Equal width – 5 bins | 94.3 (90.0 – 97.7) | 52.4 (41.1 – 70.7) | 2 (100.0) |
|  | Equal width – 10 bins | 86.9 (79.4 – 93.1) | 52.0 (35.7 – 72.4) | 3 (60.0) |
|  | Quantiles – 5 bins | 83.1 (78.2 – 90.0) | 17.5 (8.9 – 23.6) | 3 (80.0) |
|  | Quantiles – 10 bins | 82.0 (71.8 – 89.3) | 20.7 (10.7 – 36.2) | 3 (100.0) |

2. Sensitivity of thresholds for breakpoint accuracy and missingness

To evaluate the accuracy of the estimated breakpoints from the M4, BCPA, and segclust2d models, thresholds were chosen as to how close an estimate needed to be to the true breakpoints to be considered accurate. Additionally, we wanted to assess how frequently the models completed missed a true breakpoint and did not estimate any breakpoints in the vicinity. To do so, this required relatively subjective decisions as to how many observations away the estimates needed to be for classification as accurate or missing. We addressed this issue by evaluating the 20 simulations generated from unusual distributions (i.e., truncated normal, beta, uniform) as part of a sensitivity analysis.

We considered three sets of different thresholds on which to assess the breakpoint estimates from each model: 1) accurate if within ±5 observations and missing if further than 15 observations, 2) accurate if within ±10 observations and missing if further than 30 observations, and 3) accurate if within ±20 observations and missing if further than 50 observations. While the accuracy of breakpoint estimates and the proportion of missed breakpoints change slightly across different sets of thresholds (Figs S1.2, S1.3), the relationship among models holds constant. Therefore, we opted to use the second set of thresholds as part of the formal analysis as a tradeoff between being too strict or not strict enough in our evaluation.

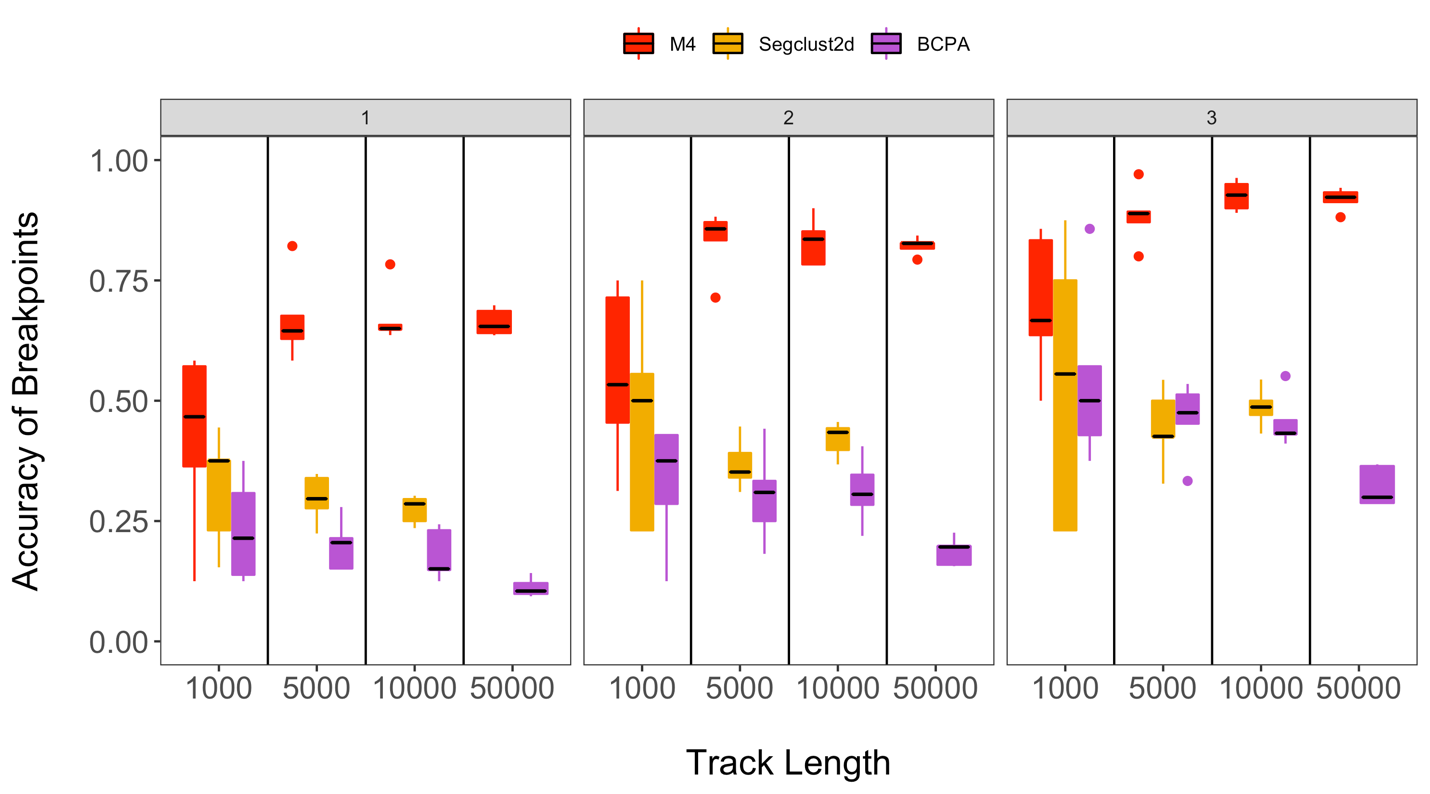

Figure S1.2 The accuracy of breakpoints estimated by each of the three models is compared among different sets of thresholds (1-3) and by track length. The different sets of thresholds include: 1) accurate if within 5 observations and missing if further than 15 observations, 2) accurate if within 10 observations and missing if further than 30 observations, and 3) accurate if within 20 observations and missing if further than 50 observations.

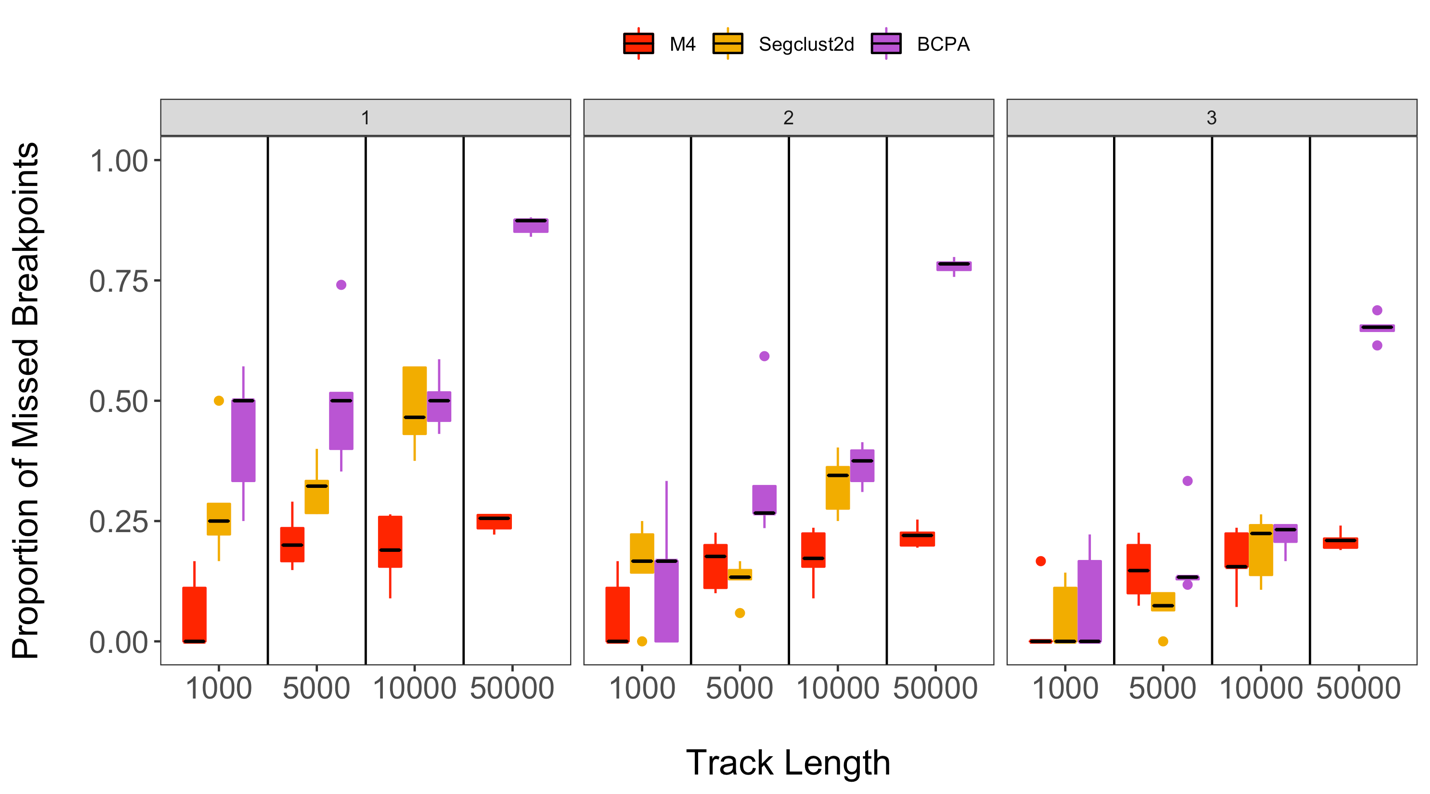

Figure S1.3 The proportion of true breakpoints missed by each of the three models is compared among different sets of thresholds (1-3) and by track length. The different sets of thresholds include: 1) accurate if within 5 observations and missing if further than 15 observations, 2) accurate if within 10 observations and missing if further than 30 observations, and 3) accurate if within 20 observations and missing if further than 50 observations.
