## Appendix S2 for "Identifying latent behavioral states in animal movement with M4, a non-parametric Bayesian method"

Appendix S2: Description of Reversible-Jump Markov Chain Monte Carlo Algorithm

Identifying latent behavioral states in animal movement with M4, a non-parametric Bayesian method

Joshua A Cullen^1*^, Caroline L Poli^2^, Robert J Fletcher, Jr.^3^, Denis Valle^1^

^1^ School of Forest Resources and Conservation, University of Florida, Gainesville, FL, USA

^2^ School of Natural Resources and Environment, University of Florida, Gainesville, FL, USA

^3^ Department of Wildlife Ecology and Conservation, University of Florida, Gainesville, FL, USA

While many questions can be addressed using standard Bayesian methods (i.e., Markov chain Monte Carlo algorithm with Metropolis-Hastings, slice samplers, or Gibbs samplers) due to their fixed number of parameters, others questions are not as tractable. When the number and value of parameters are unknown, this typically requires the fitting of numerous models where the number of parameters are varied. Since this task can become very computationally intensive and take a long time to complete using standard approaches, we rely on a reversible-jump Markov chain Monte Carlo (RJMCMC) to estimate the number and value of these parameters within a single model run.

One standard form of RJMCMC proposes the addition (‘birth’), removal (‘death’), or change (‘swap’) of parameters, which is sometimes referred to as birth-death RJMCMC. In the context of our segmentation model that estimates breakpoints to define track segments, we do not know *a priori* the number or position of these breakpoints. Therefore, the use of a RJMCMC algorithm can prove very useful for this purpose. Each iteration $k$ of the RJMCMC is defined by a model $M_{k}$ that contains a set of *P* breakpoints $\left\{ b_{1k},\ldots,b_{Pk} \right\}$. Each breakpoint is restricted to being an integer between 2 and $T_{i-1}$ across all observations, where $T_{i}$ is the total number of observations for individual $i$. During each of these iterations, there is a proposed ‘birth’, ‘death’, or ‘swap’ of breakpoints for the subsequent model $M_{k^{'}}$ (Fig S2.1). Using the set of current breakpoints, a Gibbs sampler is used to estimate the parameters that define each track segment since the form of the posterior is known due to the use of a conjugate Dirichlet prior with a Categorical distribution (see Appendix S3 for full conditional distribution). Next, the log marginal likelihood is calculated for models $M_{k}$ and $M_{k^{'}}$, where a Bayes factor is calculated to determine whether this proposed change to the breakpoints will be accepted or rejected. This process is repeated for all iterations of the RJMCMC. Additional information regarding birth-death RJMCMC can be found in Denison et al. 2002 and Zhao et al. 2013.

References

Denison DGT, Holmes CC, Mallick BK, Smith AFM. 2002. Bayesian methods for nonlinear classification and regression. Chichester, UK: John Wiley & Sons.

Zhao K, Valle D, Popescu S, Zhang X, Mallick B. 2013. Hyperspectral remote sensing of plant biochemistry using Bayesian model averaging with variable and band selection. *Remote Sensing of Environment*, 132:102-119.

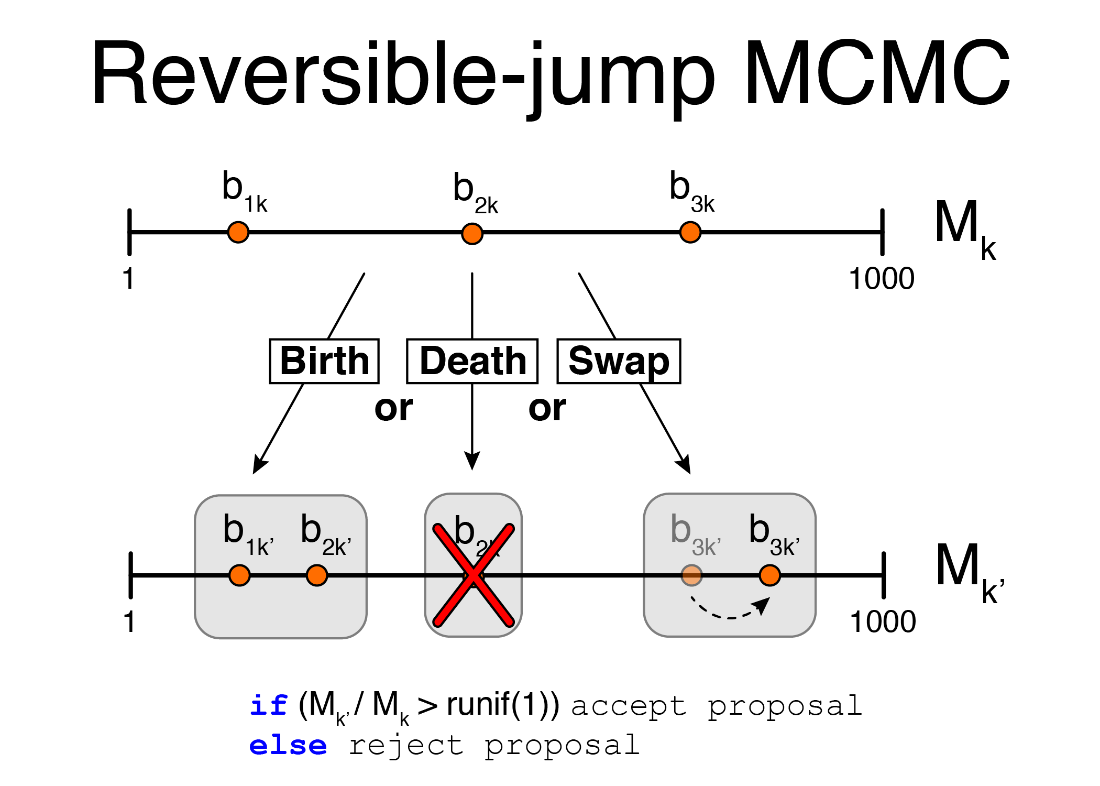

Figure S2.1 A schematic showing how the RJMCMC operates. For a single iteration of the algorithm, there are three possible proposals to change the current set of breakpoints. This proposal can either be accepted or rejected based on a comparison of the log marginal likelihood for each model.
