## Appendix S3 for "Identifying latent behavioral states in animal movement with M4, a non-parametric Bayesian method"

Appendix S3: Full Conditional Distribution of Segmentation Model

Identifying latent behavioral states in animal movement with M4, a non-parametric Bayesian method

Joshua A Cullen^1*^, Caroline L Poli^2^, Robert J Fletcher, Jr.^3^, Denis Valle^1^

^1^ School of Forest Resources and Conservation, University of Florida, Gainesville, FL, USA

^2^ School of Natural Resources and Environment, University of Florida, Gainesville, FL, USA

^3^ Department of Wildlife Ecology and Conservation, University of Florida, Gainesville, FL, USA

1) **Likelihood**

Each potential model is characterized by a set of breakpoints. In other words, model $M_{k}$ consists of breakpoints $\left\{ b_{1k},\ldots,b_{Pk} \right\}$, where $1<b_{pk}<T_{i}$, where $T_{i}$ is the number of observations for individual $i$. Given a particular model $M_{k}$, its breakpoints define time segments that are supposed to be relatively homogeneous.

In this model, we assume that data streams have been discretized into bins. Let $x_{itj}$ be the bin label for individual $i$ at time $t$ for data stream $j$. We assume for any given track segment $c$ and data stream $j$ that:

$$x_{itj}\sim Cat\left( \boldsymbol{\theta}_{\boldsymbol{cj}} \right)$$

$$\boldsymbol{\theta}_{\boldsymbol{cj}}\sim Dirichlet\left( \alpha\right)$$

Our prior for $\boldsymbol{\theta}_{\boldsymbol{cj}}$ (a vector of probabilities that sum to one) is a Dirichlet distribution with hyperparameter $\alpha$ that is equal across all bins. To enable our algorithm to efficiently explore multiple models, we need to integrate out the parameters $\boldsymbol{\theta}_{\boldsymbol{cj}}$ to obtain the marginal likelihood.

2) **Marginal likelihood**

The marginal likelihood for track segment $c$ and data stream *j* is obtained after integrating out $\boldsymbol{\theta}_{\boldsymbol{cj}}$. This integration is given by:

$$\int\left[ \prod_{t\in T_{c}} Cat\left( x_{itj} | \boldsymbol{\theta}_{\boldsymbol{cj}} \right) \right]Dirichlet\left( \boldsymbol{\theta}_{\boldsymbol{cj}} | \alpha\right)d\boldsymbol{\theta}_{\boldsymbol{cj}}$$

where $T_{c}$ is the set of all observations assigned to time segment $c$.

$$=\int\left[ \theta_{cj1}^{n_{cj1}}\times\ldots\times\theta_{cjL}^{n_{cjL}} \right]\left( \frac{\Gamma\left( \sum_{l} \alpha_{l} \right)}{\prod_{l} \Gamma\left( \alpha_{l} \right)} \right)\theta_{cj1}^{\alpha_{1}-1}\times\ldots\times\theta_{cjL}^{\alpha_{L}-1}d\boldsymbol{\theta}_{\boldsymbol{cj}}$$

where $n_{cjl}$ is the number of observations in bin $l$ in track segment $c$ from data stream *j*. If we assume that $\alpha_{1}=\ldots=\alpha_{L}$, then:

$$=\frac{\Gamma\left( L\alpha\right)}{\left[ \Gamma\left( \alpha\right) \right]^{L}}\int\theta_{cj1}^{n_{cj1}+\alpha_{1}-1}\times\ldots\times\theta_{cjL}^{n_{cjL}+\alpha_{L}-1}d\boldsymbol{\theta}_{\boldsymbol{cj}}$$

$$=\frac{\Gamma\left( L\alpha\right)}{\left[ \Gamma\left( \alpha\right) \right]^{L}}\times\frac{\prod_{l} \Gamma\left( a+n_{cjl} \right)}{\Gamma\left( L\alpha+\sum_{l} n_{cjl} \right)}$$

3) **Algorithm**

Based on the marginal likelihood, the posterior probability of model $M_{k}$ is given by:

$$p\left( M_{k} | \ldots\right)\propto\left[ \prod_{j} \prod_{c} \frac{\Gamma\left( L\alpha\right)}{\left[ \Gamma\left( \alpha\right) \right]^{L}}\times\frac{\prod_{l} \Gamma\left( \alpha_{l}+n_{cjl} \right)}{\Gamma\left( L\alpha+\sum_{l} n_{cjl} \right)} \right]p\left( M_{k} \right)$$

In this expression, the prior probability for model *k* (i.e., $p\left( M_{k} \right)$) is given by:

$$p\left( M_{k} \right)=\left( \begin{aligned} T_{i} \\ P \end{aligned} \right)^{-1}\times\frac{1}{T_{i}+1}$$

where $P$ is the number of breakpoints in $M_{k}$.

Our Metropolis-Hastings algorithm iteratively proposes the creation of new breakpoints (birth), deletion of an existing breakpoint (death), or the swap of an existing for a new breakpoint. These proposals are accepted or rejected with probability given by:

$$p_{acceptance}=\min\left( 1,\frac{p\left( M_{k}^{new} | \ldots\right)}{p\left( M_{k}^{old} | \ldots\right)} \right)$$

Additional information regarding how similar algorithms work can be found in (Denison et al. 2002).
