## Appendix S4 for "Identifying latent behavioral states in animal movement with M4, a non-parametric Bayesian method"

Appendix S4: Use of MAP Breakpoint Estimates Compared to Full Posterior

Identifying latent behavioral states in animal movement with M4, a non-parametric Bayesian method

Joshua A Cullen^1*^, Caroline L Poli^2^, Robert J Fletcher, Jr.^3^, Denis Valle^1^

^1^ School of Forest Resources and Conservation, University of Florida, Gainesville, FL, USA

^2^ School of Natural Resources and Environment, University of Florida, Gainesville, FL, USA

^3^ Department of Wildlife Ecology and Conservation, University of Florida, Gainesville, FL, USA

The selection of a single set of breakpoints is important to consistently define a set of track segments on which to make behavioral inference using the LDA model of the M4 framework. Although many sets of breakpoints are estimated within the posterior distribution, we only select those defined by the Maximum a Posteriori (MAP) estimate. In other words, we select the set of breakpoints which results in the greatest log marginal likelihood. Since these breakpoints come from the “best” model, they are also expected to be representative of other “good” posterior estimates from the RJMCMC. We select the MAP estimate since it is inherently difficult to account for uncertainty in breakpoint location estimates as captured via the full posterior distribution.

One possible method to account for breakpoint uncertainty would be to retain a specified proportion of breakpoints that appear across all posterior samples (i.e., only keep breakpoints proposed by 80% of models). However, this does not easily take into account that breakpoints can be moved from model to model of the RJMCMC, where breakpoints may vary across a range of neighboring positions to characterize a single breakpoint. In this case, some breakpoints may be missed if the models do not all agree on a consensus breakpoint position.

Another method that may account for uncertainty in the breakpoint positions is the identification of peaks from density plots of the posterior. These plots help to visualize what locations are most likely to be consensus breakpoints across all posterior estimates, but do not provide any insight with regard to rapid changes in behavioral state. Since the breakpoint densities are calculated using a smoothing parameter for a kernel density function, this may result in one large peak that characterizes multiple breakpoints as opposed to multiple narrow peaks.

A plot of the MAP breakpoint estimates overlaying the densities of all posterior breakpoint estimates (using a small smoothing parameter) from the snail kite analysis shows good agreement between both sets of estimates (Fig S4.1). This indicates that the use of MAP estimates to define track segments appears to be supported by the posterior distribution as a whole and should not pose a problem when making inference.

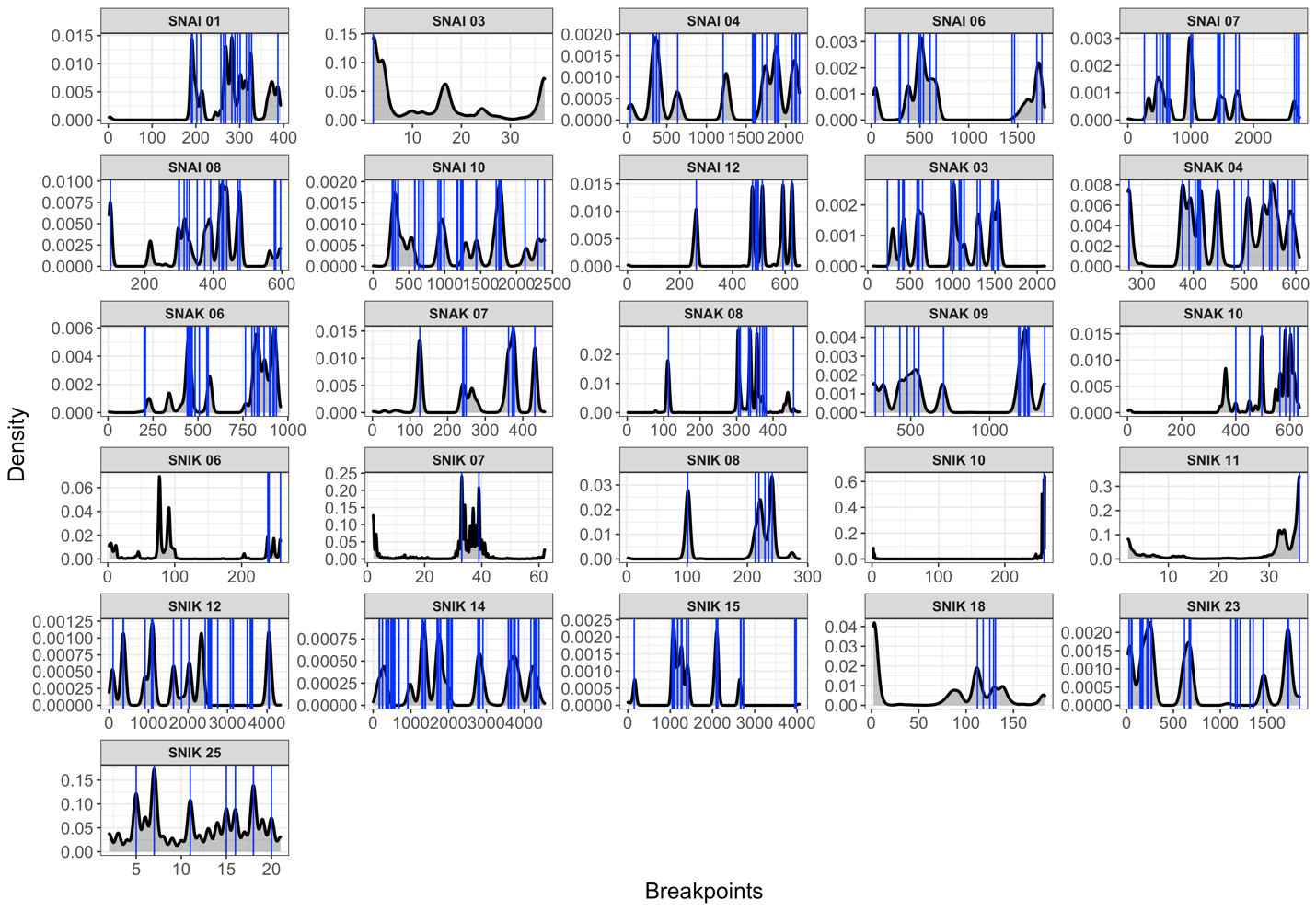

Figure S4.1. Density plots are shown for the breakpoint estimates of the posterior distribution for each individual snail kite in comparison to the MAP estimates (blue lines). MAP estimates show good agreement with most peaks from the density of the posterior estimates, particularly for individuals with > 200 observations.
