## Appendix S5 for "Identifying latent behavioral states in animal movement with M4, a non-parametric Bayesian method"

Appendix S5: Full Conditional Distribution of LDA Clustering Model

Identifying latent behavioral states in animal movement with M4, non-parametric Bayesian method

Joshua A Cullen^1*^, Caroline L Poli^2^, Robert J Fletcher, Jr.^3^, Denis Valle^1^

^1^ School of Forest Resources and Conservation, University of Florida, Gainesville, FL, USA

^2^ School of Natural Resources and Environment, University of Florida, Gainesville, FL, USA

^3^ Department of Wildlife Ecology and Conservation, University of Florida, Gainesville, FL, USA

The Full Conditional Distributions (FCD’s) that are used as part of our Gibbs sampler are given below.

**1)** Parameter $\boldsymbol{\phi}_{\boldsymbol{kj}}$

$$p\left( \boldsymbol{\phi}_{\boldsymbol{kj}} | \ldots\right)\propto\left[ \prod_{i} \prod_{j} \prod_{c} \prod_{t} Cat\left( y_{ijct} | \boldsymbol{\phi}_{\boldsymbol{kj}} \right)^{I\left( z_{ijct}=k \right)} \right]\mathrm{Dirichlet}\left( \boldsymbol{\phi}_{\boldsymbol{kj}} | \alpha\right)$$

$$\propto\left[ \phi_{kj1}^{\sum_{i} \sum_{j} \sum_{c} \sum_{t} I\left( y_{ijct}=1, z_{ijct}=k \right)}\times\ldots\times\phi_{kjB_{j}}^{\sum_{i} \sum_{j} \sum_{c} \sum_{t} I\left( y_{ijct}=B_{j}, z_{ijct}=k \right)} \right]\phi_{kj1}^{\alpha-1}\times\ldots\times\phi_{kjB_{j}}^{\alpha-1}$$

$$\propto\left[ \phi_{kj1}^{n_{kj1}}\times\ldots\times\phi_{kjB_{j}}^{n_{kjB_{j}}} \right]\phi_{kj1}^{\alpha-1}\times\ldots\times\phi_{kjB_{j}}^{\alpha-1}$$

where $B_{j}$ is the total number of bins in $y_{ijct}$ and $n_{bkj}$ is the number of $y_{ijct}$ observations across all individuals and track segments that fall in bin $b$ for data stream $j$ and that were assigned to cluster $k$. More specifically,

$$n_{kjb}=\sum_{i} \sum_{j} \sum_{c} \sum_{t} I\left( y_{ijct}=b, z_{ijct}=k \right)$$

As a result, we have that:

$$p\left( \boldsymbol{\phi}_{\boldsymbol{kj}} | \ldots\right)\propto\phi_{kj1}^{n_{kj1}+\alpha-1}\times\ldots\times\phi_{kjB_{j}}^{n_{kjB_{j}}+\alpha-1}$$

which implies that:

$$\boldsymbol{\phi}_{\boldsymbol{kj}}\sim Dirichlet\left( \left[ n_{kj1}+\alpha,\ldots,n_{kjB_{j}}+\alpha\right] \right)$$

**2)** Parameter $z_{ijct}$

$$p\left( z_{ijct}=k | \ldots\right)\propto Cat\left( y_{ijct}=b | \boldsymbol{\phi}_{\boldsymbol{kj}} \right)\mathrm{Cat}\left( z_{ijct} | \boldsymbol{\theta}_{\boldsymbol{ic}} \right)$$

$$\propto\phi_{kjb}\theta_{ick}$$

We sample $z_{ijct}$ from a categorical distribution with probability proportional to the expression above.

**3)** Parameter $V_{ick}$

$$p\left( V_{ick} | \boldsymbol{\ldots} \right)\boldsymbol{\propto}\left[ \prod_{j} Cat\left( z_{ijct} | \boldsymbol{\theta}_{\boldsymbol{ic}} \right) \right]\mathrm{Beta}\left( V_{ick} | 1,\gamma\right)$$

$$\boldsymbol{\propto}\left[ \prod_{j} \theta_{ic1}^{I\left( z_{ijct}=1 \right)}\times\ldots\times\theta_{icK}^{I\left( z_{ijct}=K \right)} \right]\left( 1-V_{ick} \right)^{\gamma-1}$$

$$\boldsymbol{\propto}\left[ \theta_{ic1}^{\sum_{j} I\left( z_{ijct}=1 \right)}\times\ldots\times\theta_{icK}^{\sum_{j} I\left( z_{ijct}=K \right)} \right]\left( 1-V_{ick} \right)^{\gamma-1}$$

$$\boldsymbol{\propto}\left[ V_{ick}^{\sum_{j} I\left( z_{ijct}=k \right)}\times\left( 1-V_{ick} \right)^{\sum_{j} I\left( z_{ijct}>k \right)} \right]\left( 1-V_{ick} \right)^{\gamma-1}$$

Let $n_{ick}=\sum_{j} I\left( z_{ijct}=k \right)$ and $n_{ic\left( >k \right)}=\sum_{j} I\left( z_{ijct}>k \right)$. Therefore:

$$\boldsymbol{\propto}V_{ick}^{\left( n_{ick}+1 \right)-1}\times\left( 1-V_{ick} \right)^{n_{ic\left( >k \right)}+\gamma-1}$$

This implies that:

$$V_{ick}\sim Beta\left( n_{ick}+1,n_{ic\left( >k \right)}+\gamma\right)$$
