## Appendix S6 for "Identifying latent behavioral states in animal movement with M4, a non-parametric Bayesian method"

Appendix S6: R Functions for Bayesmove Workflow

Identifying latent behavioral states in animal movement with M4, a non-parametric Bayesian method

Joshua A Cullen^1*^, Caroline L Poli^2^, Robert J Fletcher, Jr.^3^, Denis Valle^1^

^1^ School of Forest Resources and Conservation, University of Florida, Gainesville, FL, USA

^2^ School of Natural Resources and Environment, University of Florida, Gainesville, FL, USA

^3^ Department of Wildlife Ecology and Conservation, University of Florida, Gainesville, FL, USA

Table S6.1 List of primary bayesmove functions and their usage within a workflow.

| Stage | Function | Usage |
| --- | --- | --- |
| Prepare | *prep_data()* | Calculates step lengths, turning angles, and net-squared displacement based on the coordinates for each track and calculates time steps based on the date. |
|  | *discrete_move_var()* | Converts data streams from continuous to discrete variables based upon pre-specified bin limits for each data stream. |
| Segment | *segment_behavior()* | Segments individual tracks into relatively homogeneous units of movement by estimating breakpoints. |
|  | *assign_tseg()* | Assigns observations to track segments per individual based upon the breakpoints estimated by the segmentation model. |
| Cluster | *cluster_segments()* | Clusters pooled track segments from all individuals to estimate the likely number of behavioral states, the state-dependent distributions for each data stream, and the proportion of each behavior per track segment. |
|  | *assign_behavior()* | Assigns behavioral state estimates to observations, where all observations within a given track segment are defined by a single set of proportions for each behavioral state as well as the dominant state. |
