## Appendix S7 for "Identifying latent behavioral states in animal movement with M4, a non-parametric Bayesian method"

Appendix S7: Trace-Plots from Non-Parametric Bayesian Models

Identifying latent behavioral states in animal movement with M4, a non-parametric Bayesian method

Joshua A Cullen^1*^, Caroline L Poli^2^, Robert J Fletcher, Jr.^3^, Denis Valle^1^

^1^ School of Forest Resources and Conservation, University of Florida, Gainesville, FL, USA

^2^ School of Natural Resources and Environment, University of Florida, Gainesville, FL, USA

^3^ Department of Wildlife Ecology and Conservation, University of Florida, Gainesville, FL, USA

**1. Simulations**

*1.1 Uncommon distributions*

*1.1.1 M4 segmentation model*

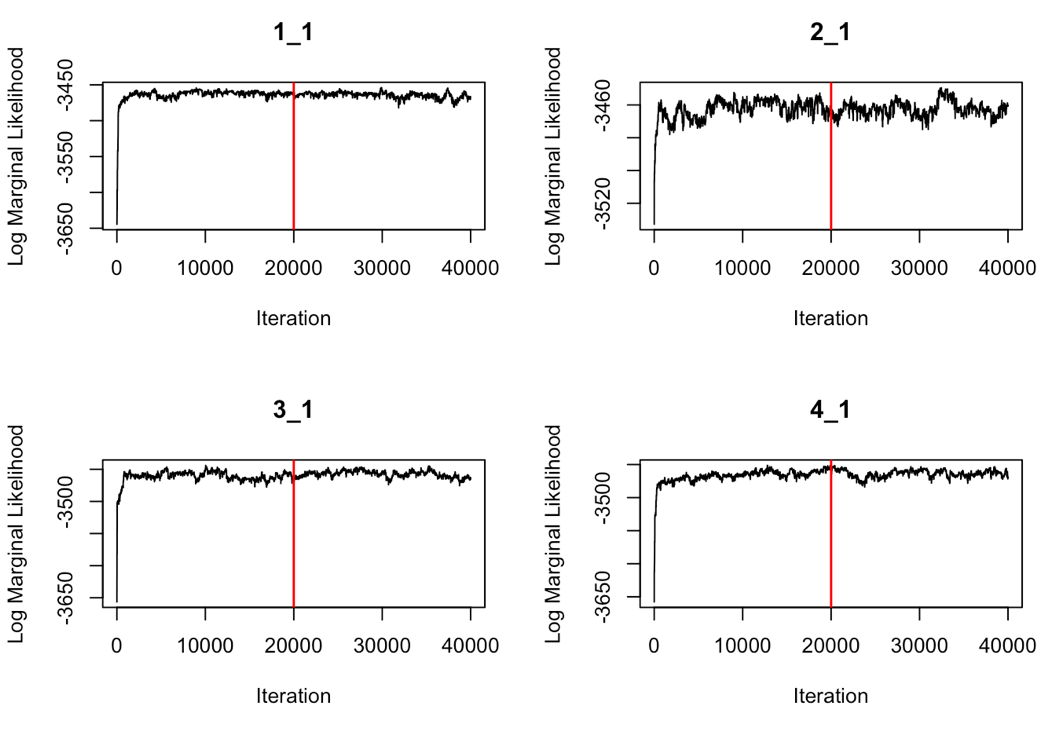

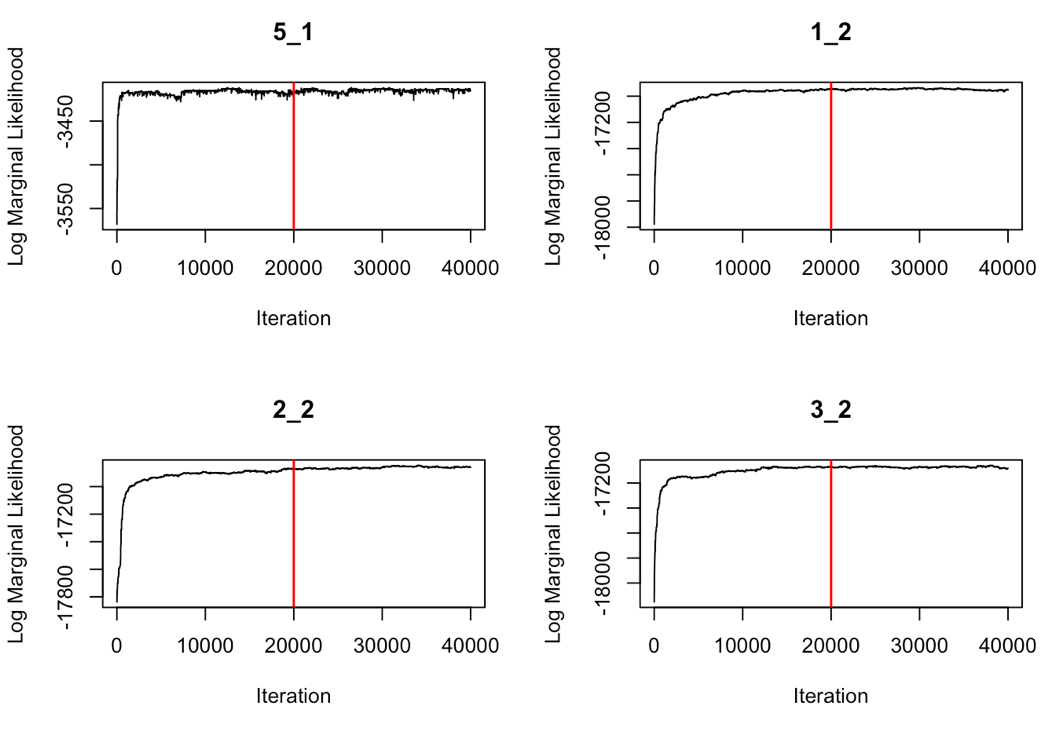

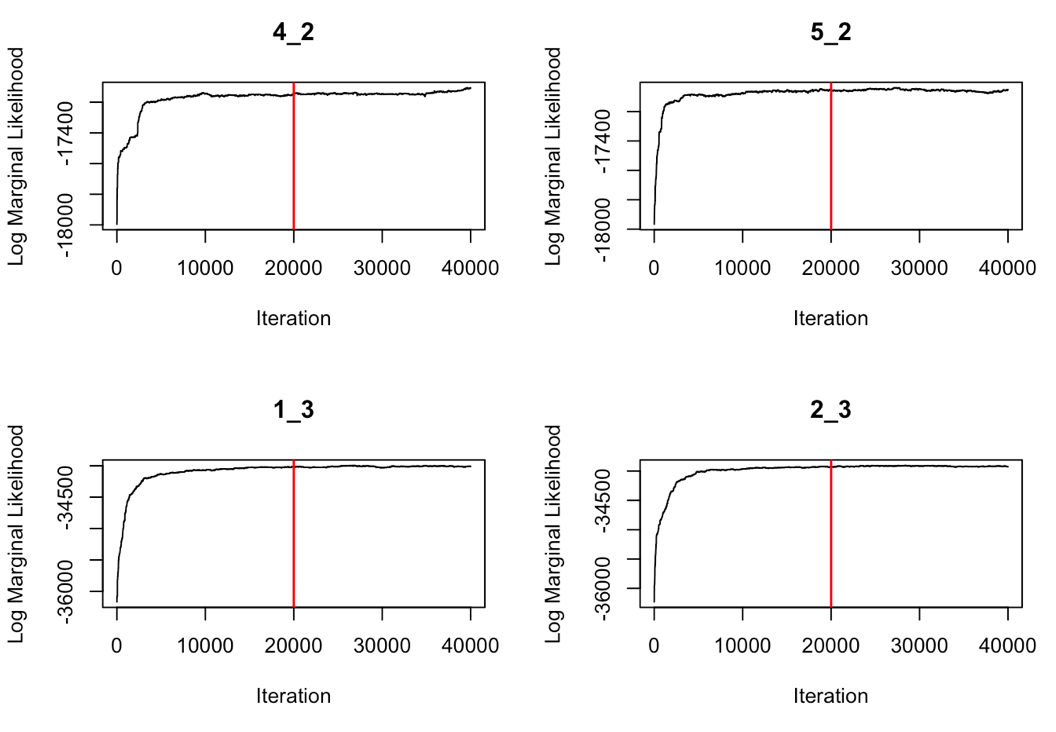

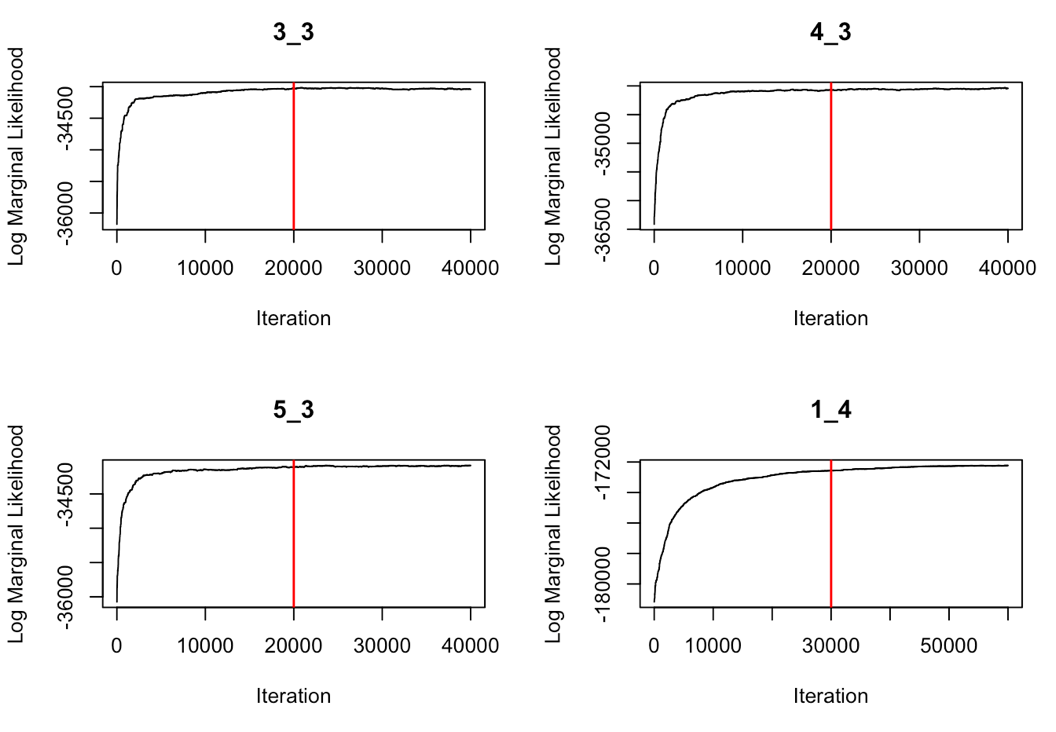

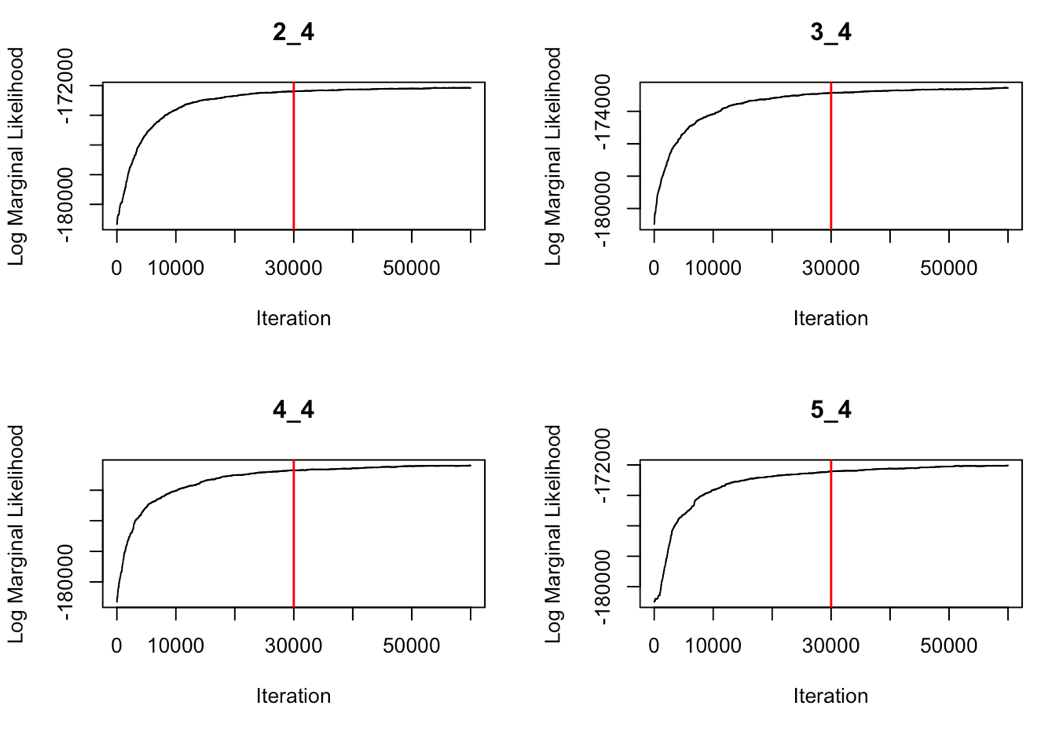

Figure S7.1 Trace-plots from the M4 segmentation model after analyzing simulations generated by uncommon distributions (i.e., truncated normal, uniform, beta). The numbers above each trace-plot refers to the simulation ID, where IDs *_1, *_2, *_3, and *_4 designate tracks of 1000, 5000, 10000, and 50000 observations, respectively. The first part of the ID (i.e., 1_*, 2_*, 3_*, 4_*, 5_*) indicates each of the five simulated tracks for a given track length. The MAP estimate of breakpoints was sampled from the posterior distribution, which is to the right of the burn-in (red line).

*1.1.2 M4 LDA model*

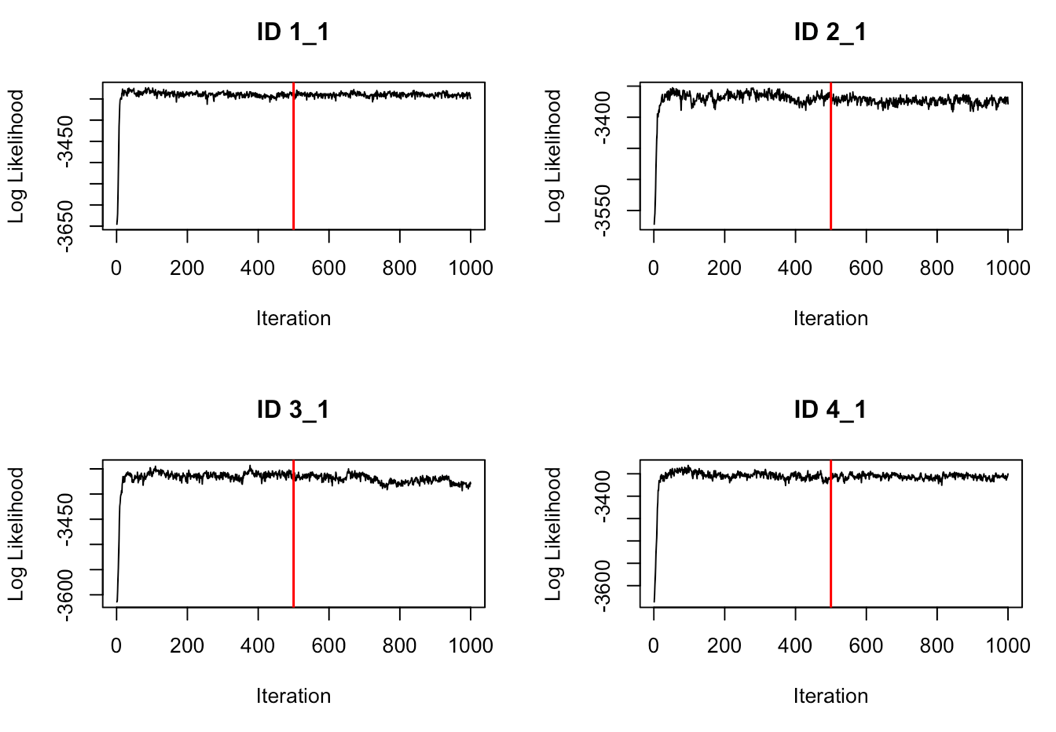

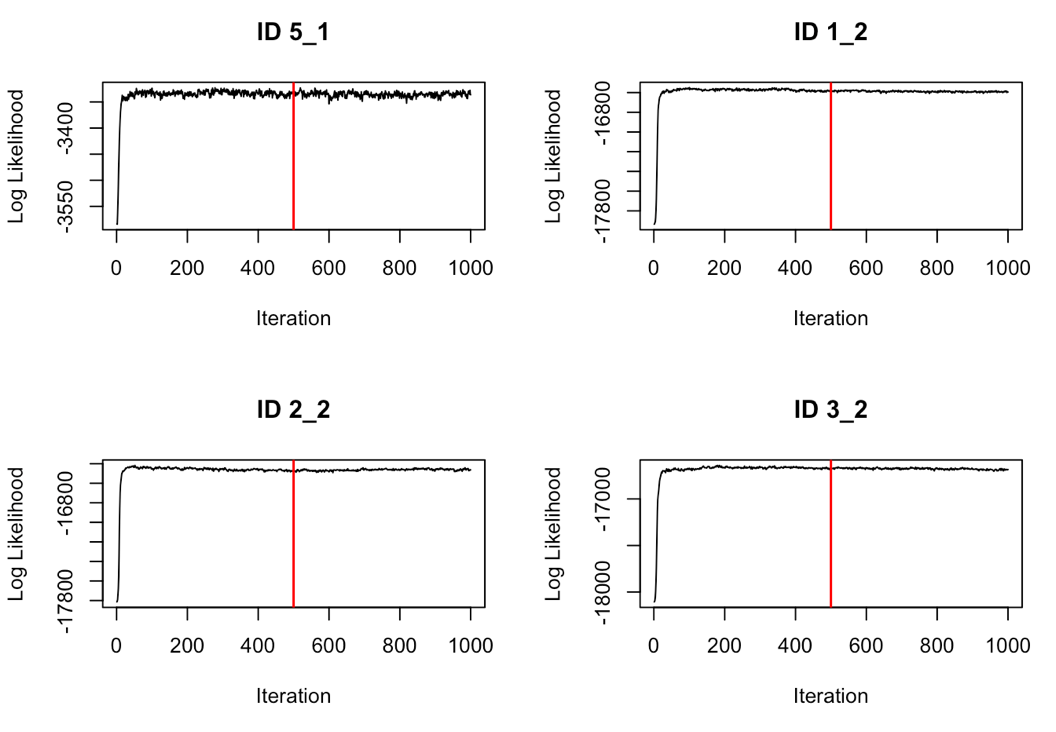

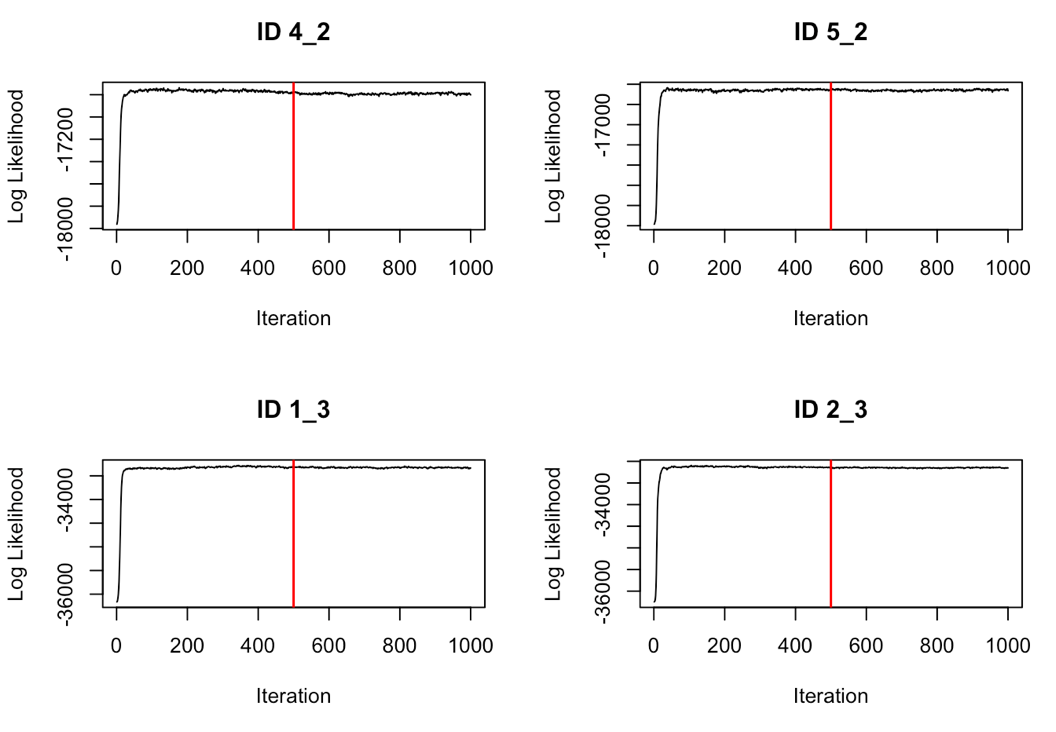

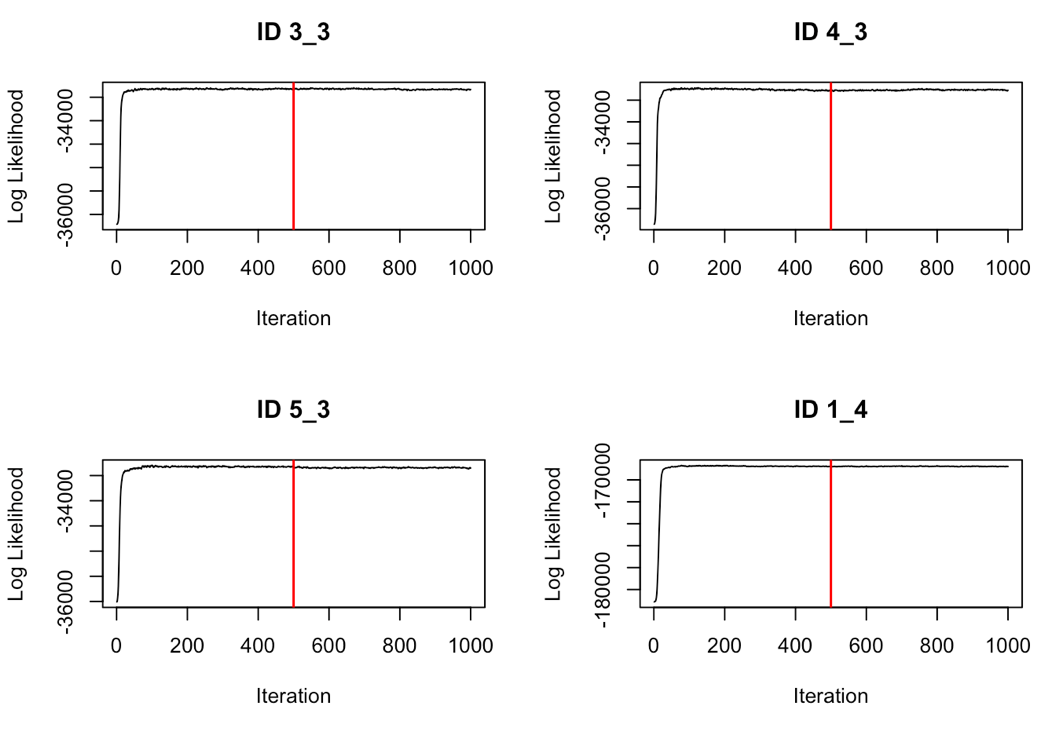

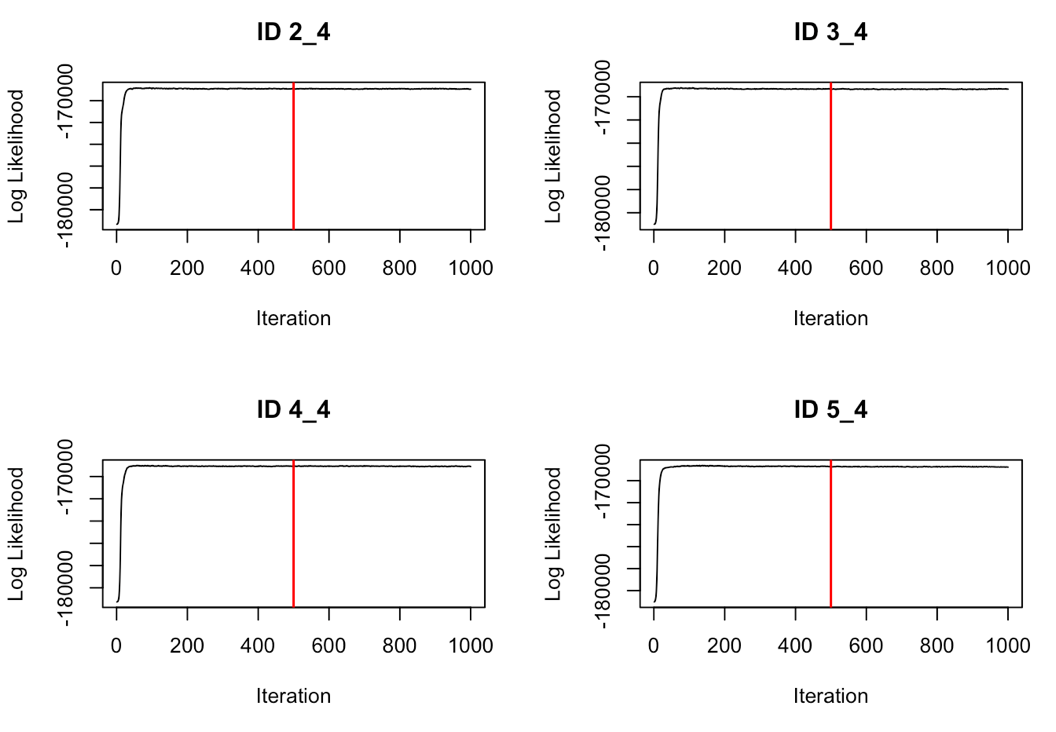

Figure S7.2 Trace-plots from the M4 LDA model after analyzing simulations generated by uncommon distributions (i.e., truncated normal, uniform, beta). The numbers above each trace-plot refers to the simulation ID, where IDs *_1, *_2, *_3, and *_4 designate tracks of 1000, 5000, 10000, and 50000 observations, respectively. The first part of the ID (i.e., 1_*, 2_*, 3_*, 4_*, 5_*) indicates each of the five simulated tracks for a given track length. The posterior distribution is considered to be to the right of the burn-in (red line).

*1.2 Common parametric distributions*

*1.2.1 M4 segmentation model*

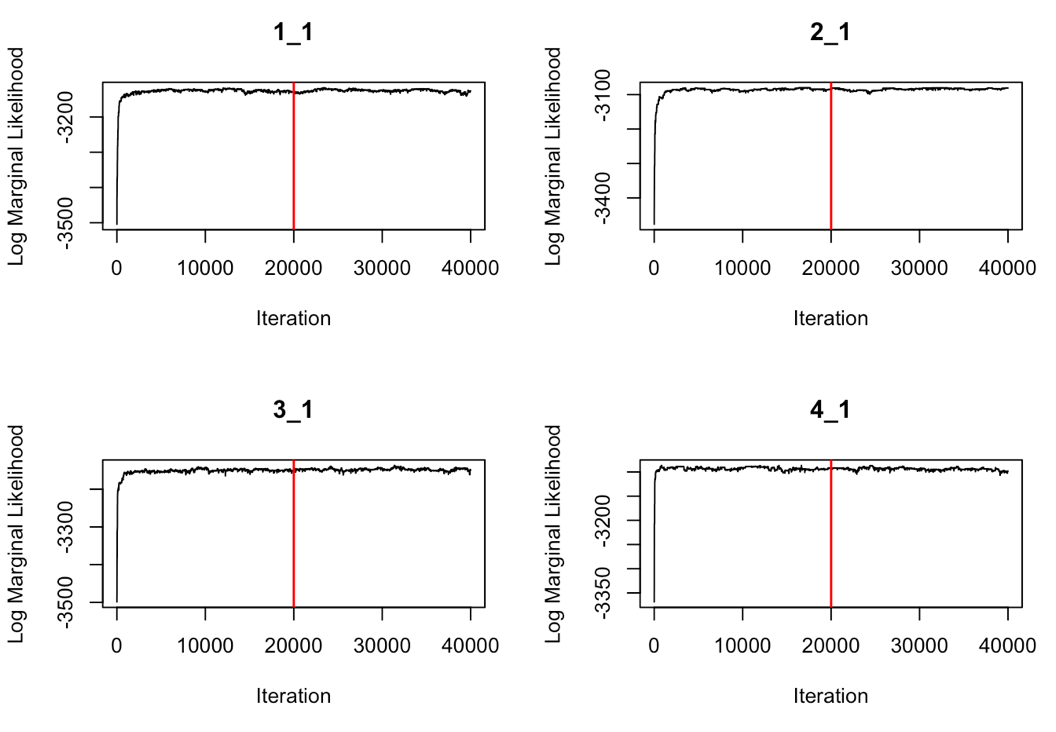

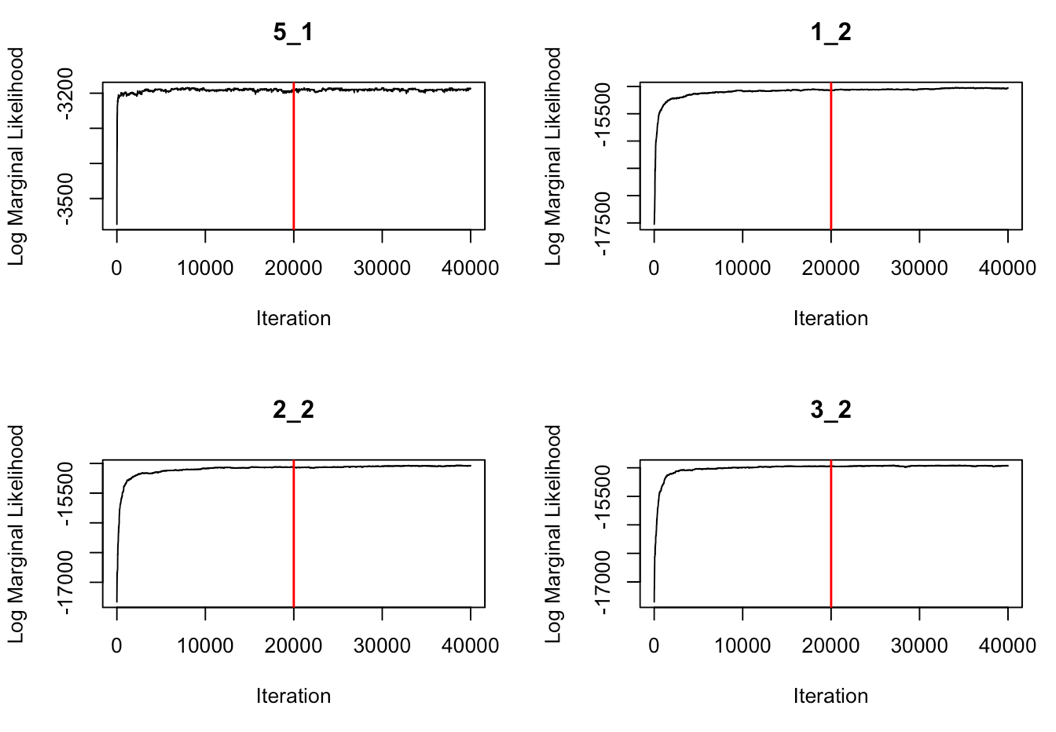

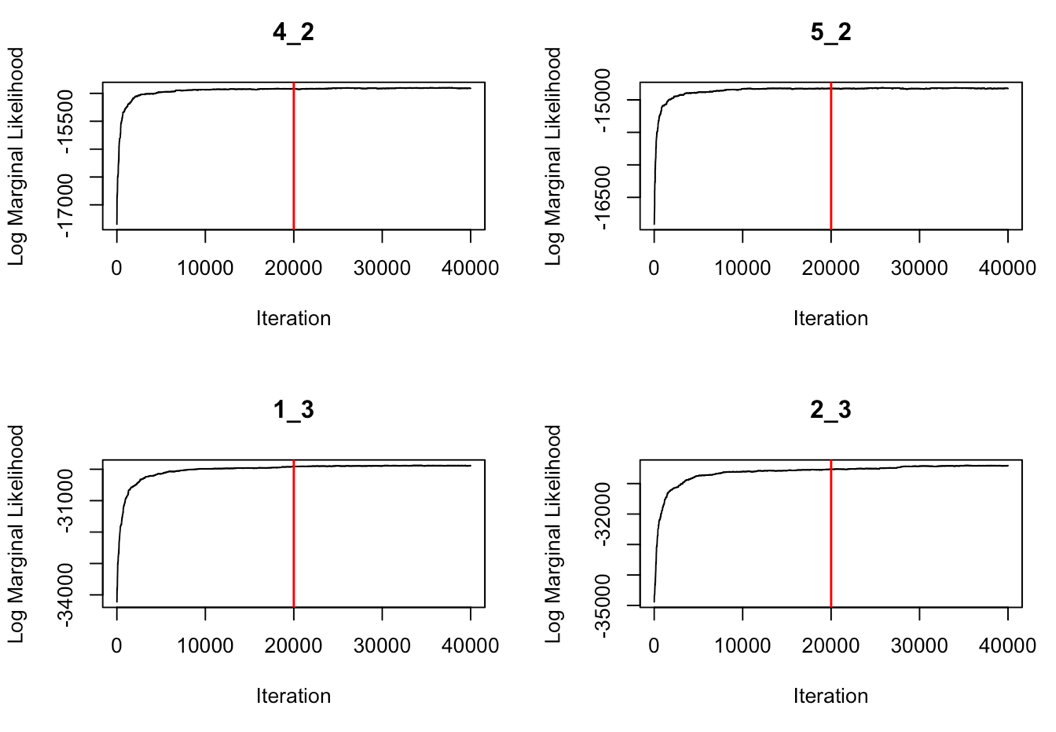

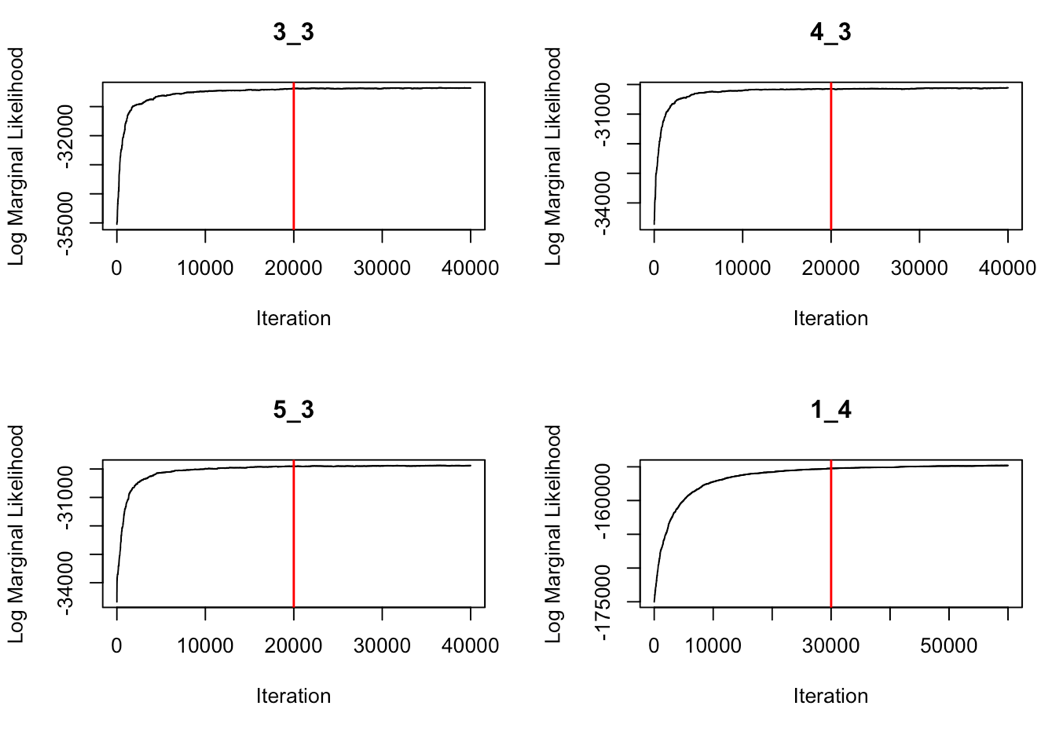

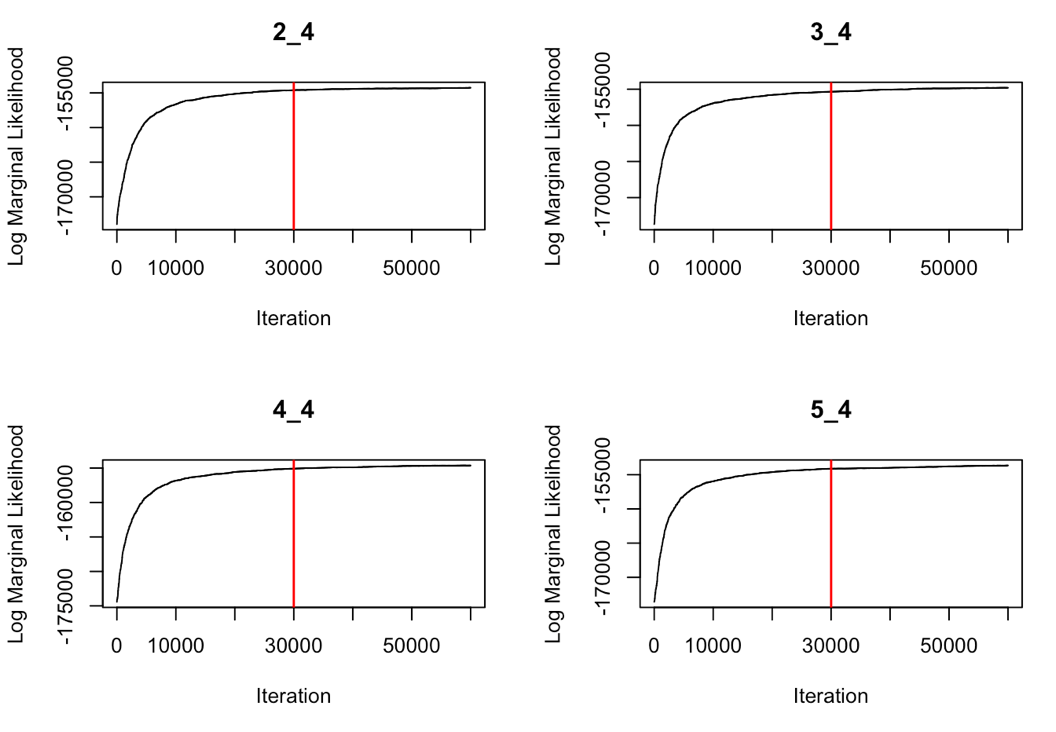

Figure S7.3 Trace-plots from the M4 segmentation model after analyzing simulations generated by common distributions (i.e. gamma, wrapped Cauchy). The numbers above each trace-plot refers to the simulation ID, where IDs *_1, *_2, *_3, and *_4 designate tracks of 1000, 5000, 10000, and 50000 observations, respectively. The first part of the ID (i.e., 1_*, 2_*, 3_*, 4_*, 5_*) indicates each of the five simulated tracks for a given track length. The MAP estimate of breakpoints was sampled from the posterior distribution, which is to the right of the burn-in (red line).

*1.2.2 M4 LDA model*

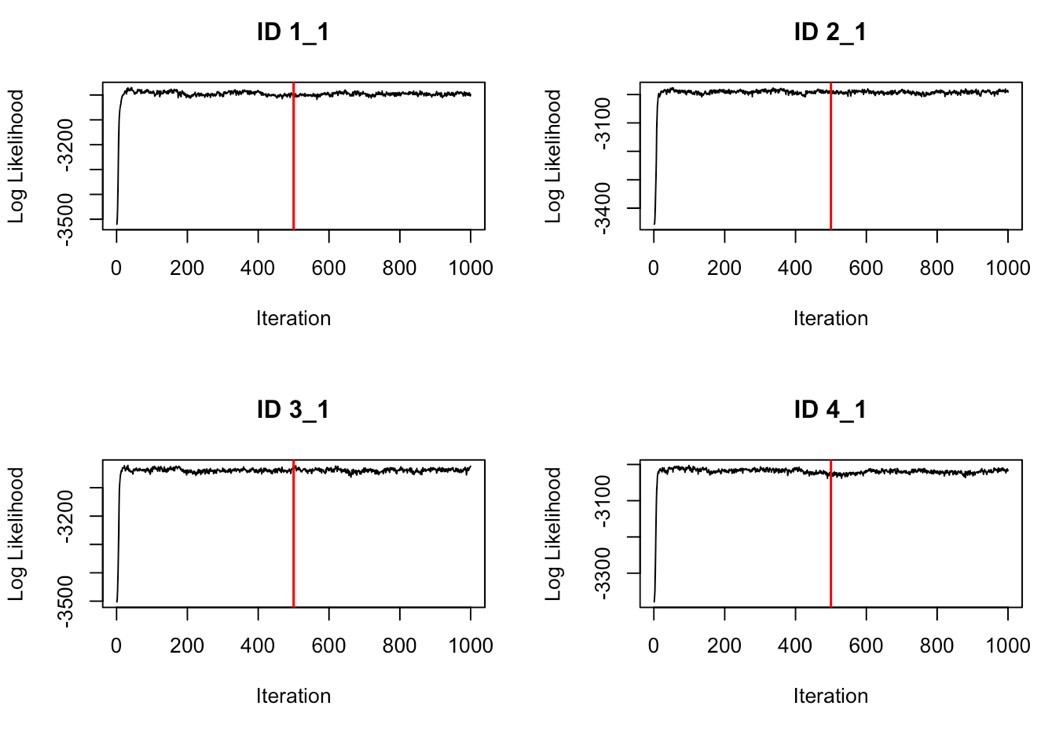

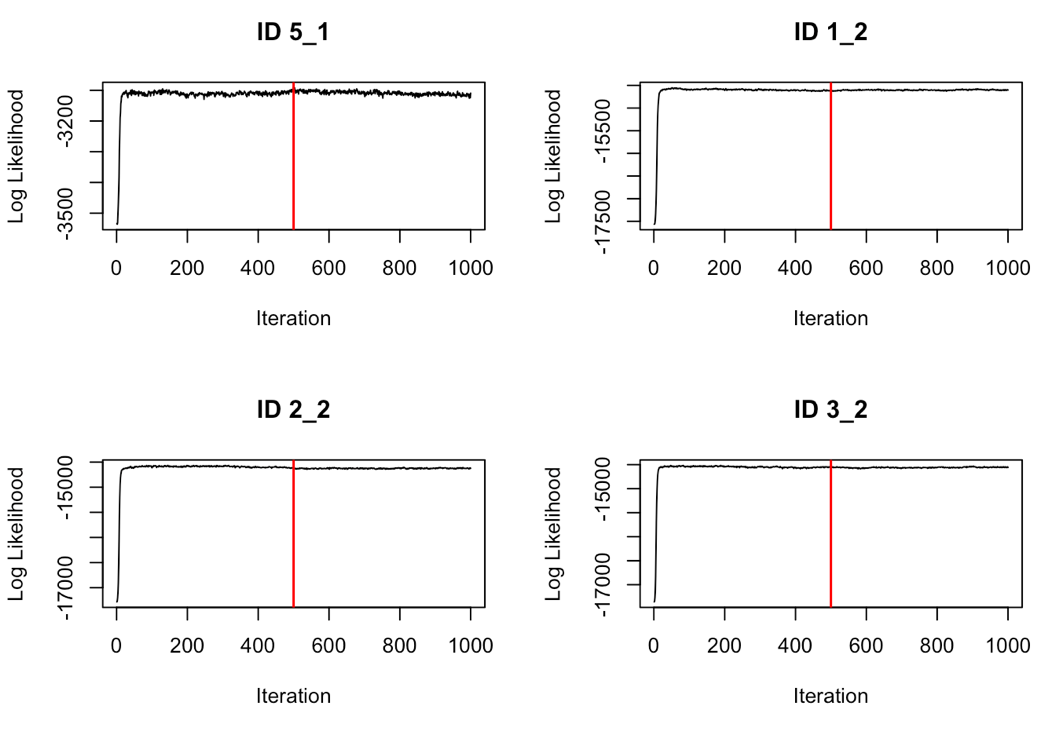

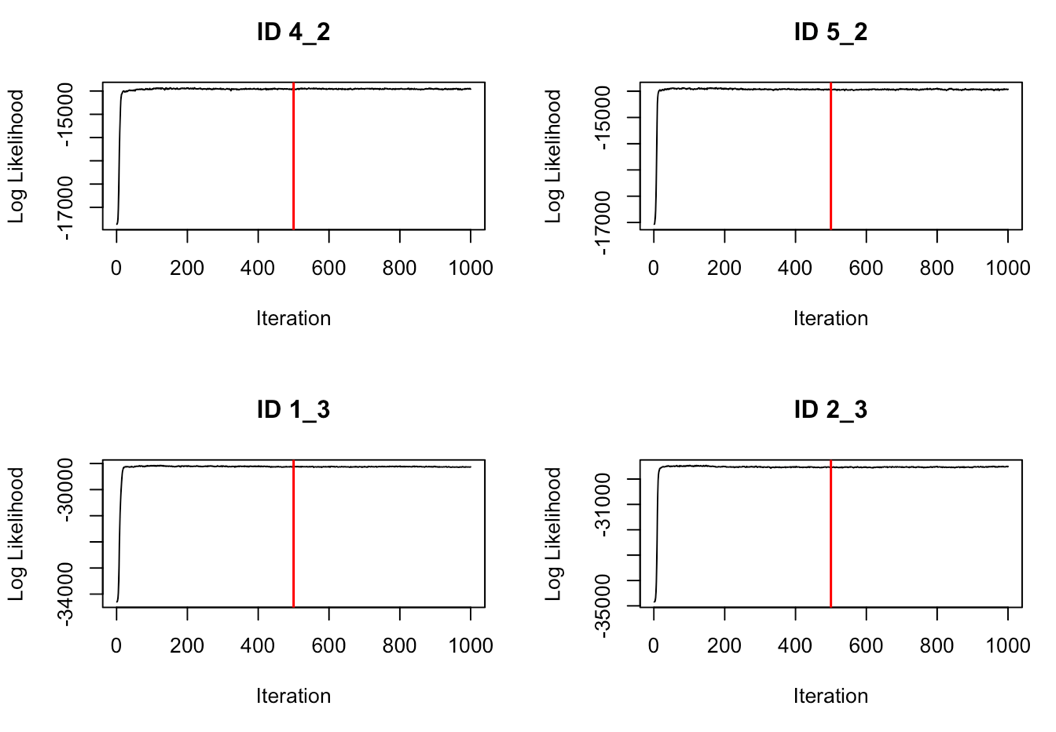

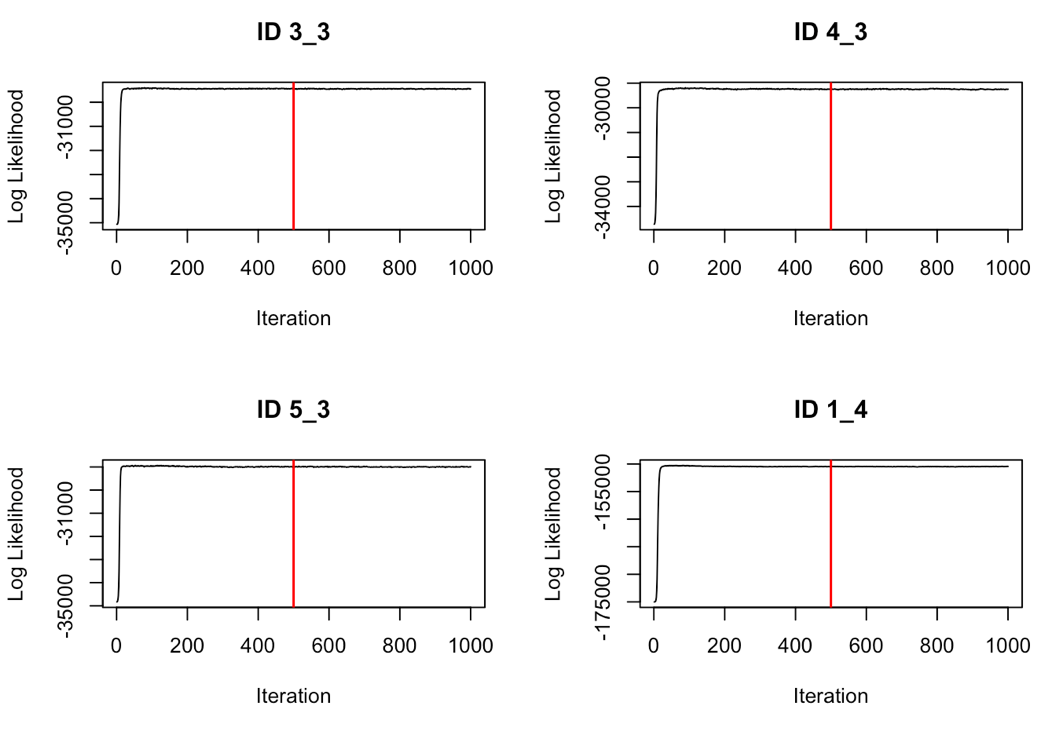

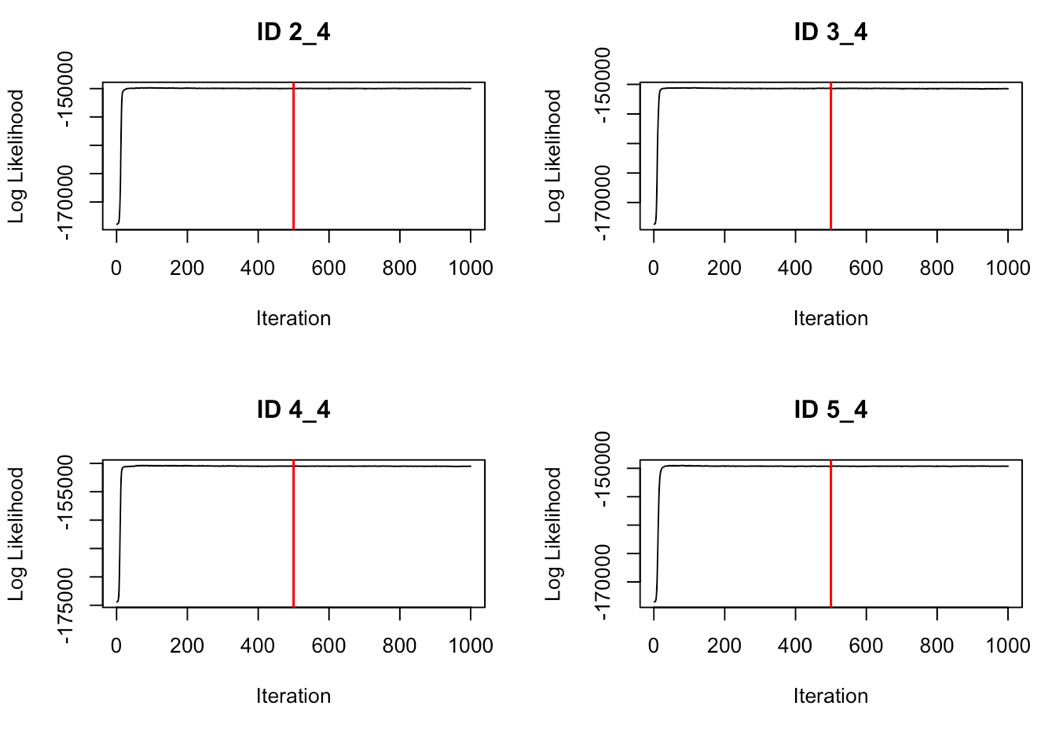

Figure S7.4 Trace-plots from the M4 LDA model after analyzing simulations generated by common distributions (i.e. gamma, wrapped Cauchy). The numbers above each trace-plot refers to the simulation ID, where IDs *_1, *_2, *_3, and *_4 designate tracks of 1000, 5000, 10000, and 50000 observations, respectively. The first part of the ID (i.e., 1_*, 2_*, 3_*, 4_*, 5_*) indicates each of the five simulated tracks for a given track length. The posterior distribution is considered to be to the right of the burn-in (red line).

**2. Empirical analyses**

*2.1 M4 segmentation model*

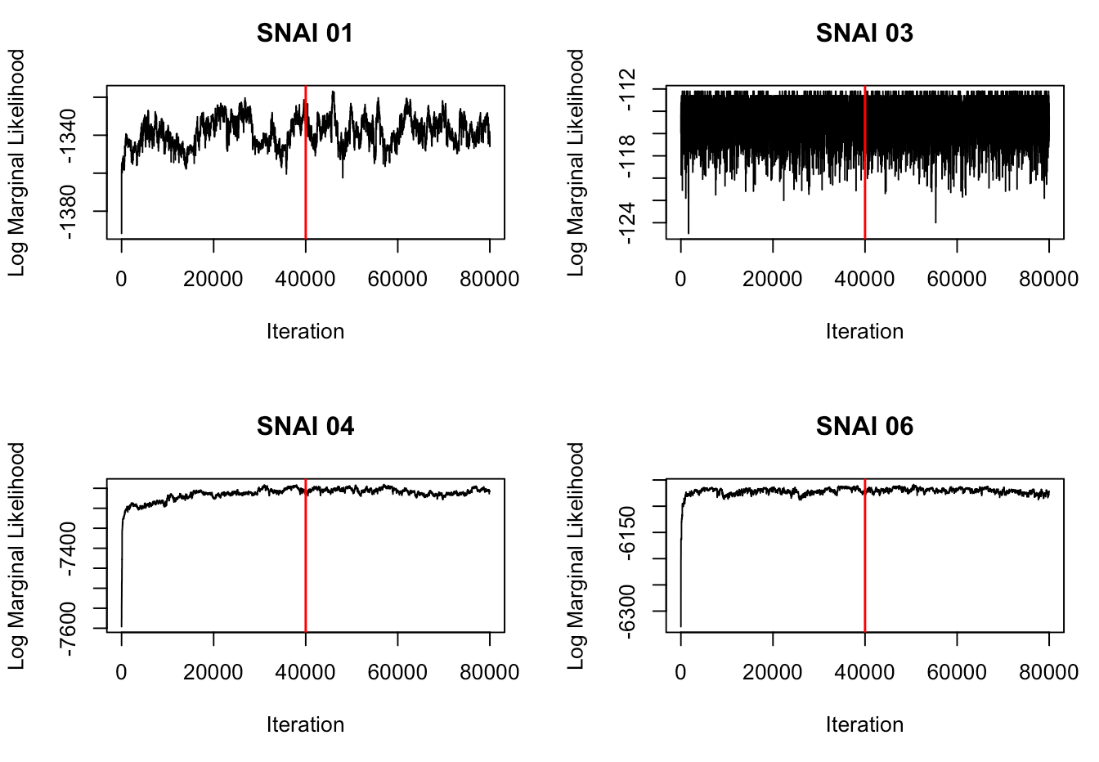

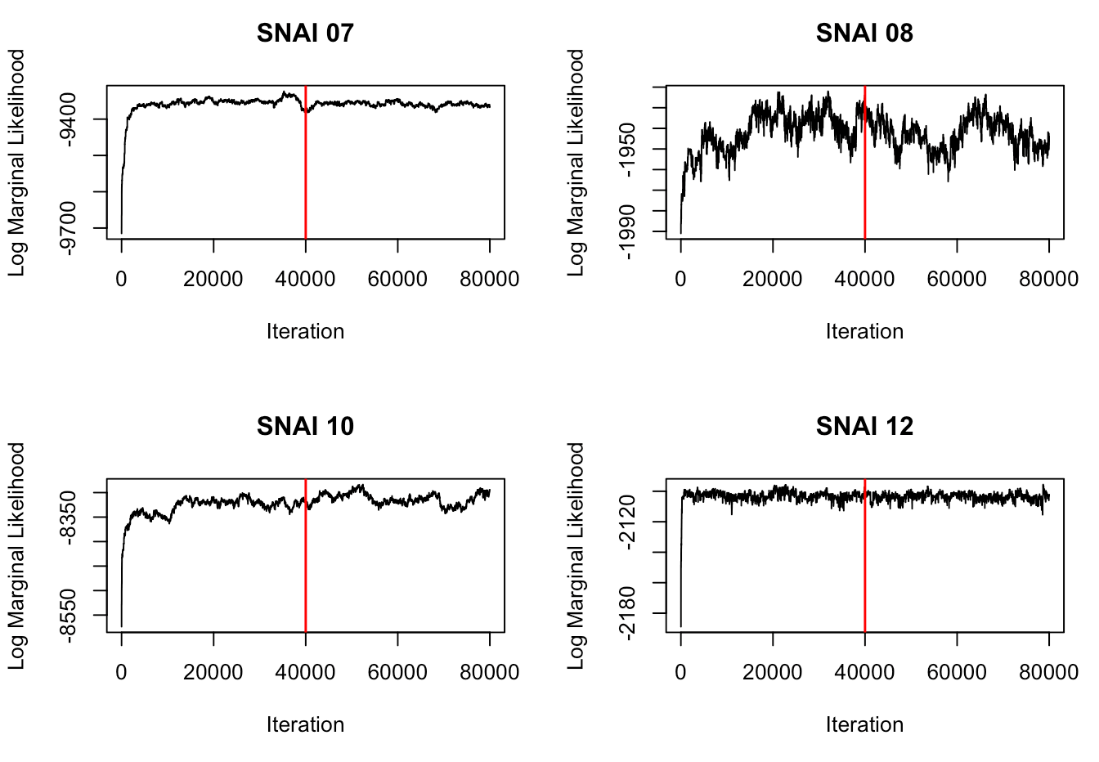

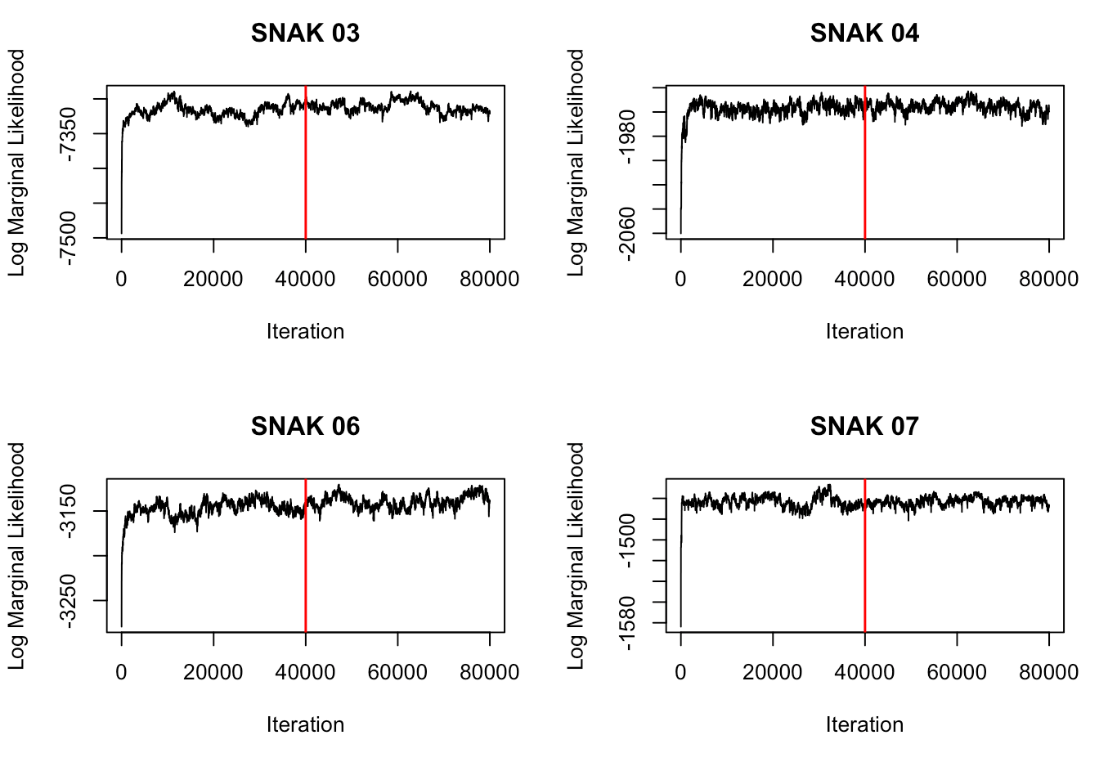

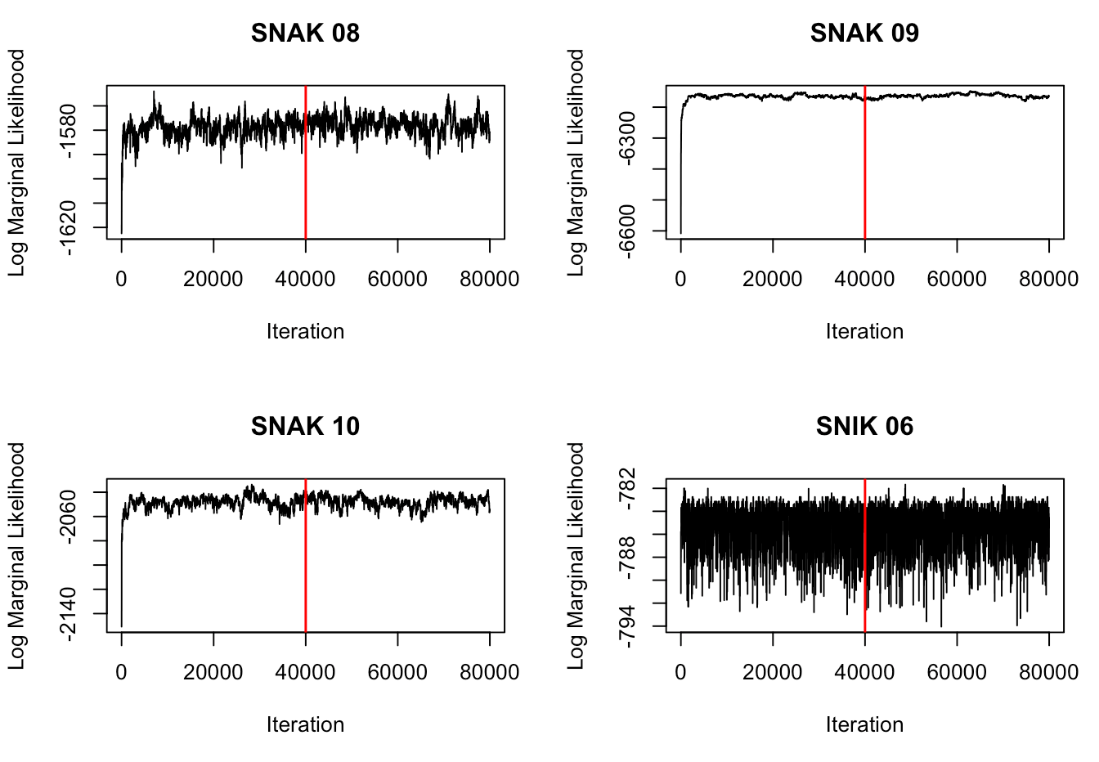

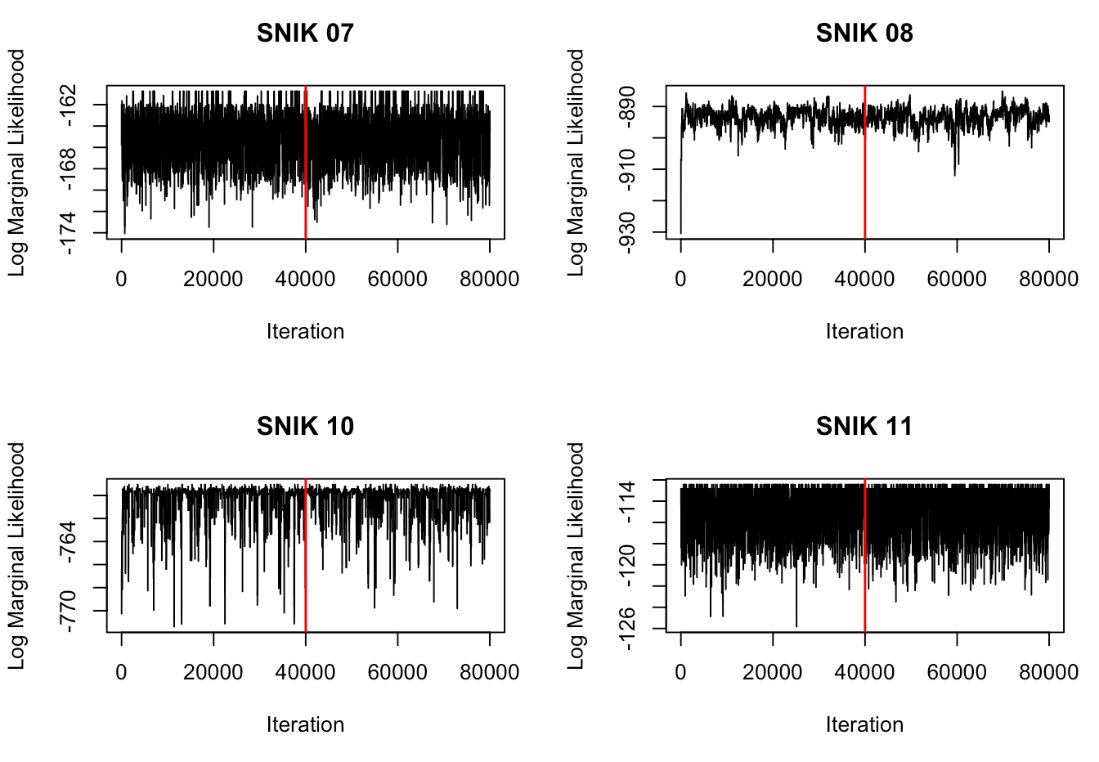

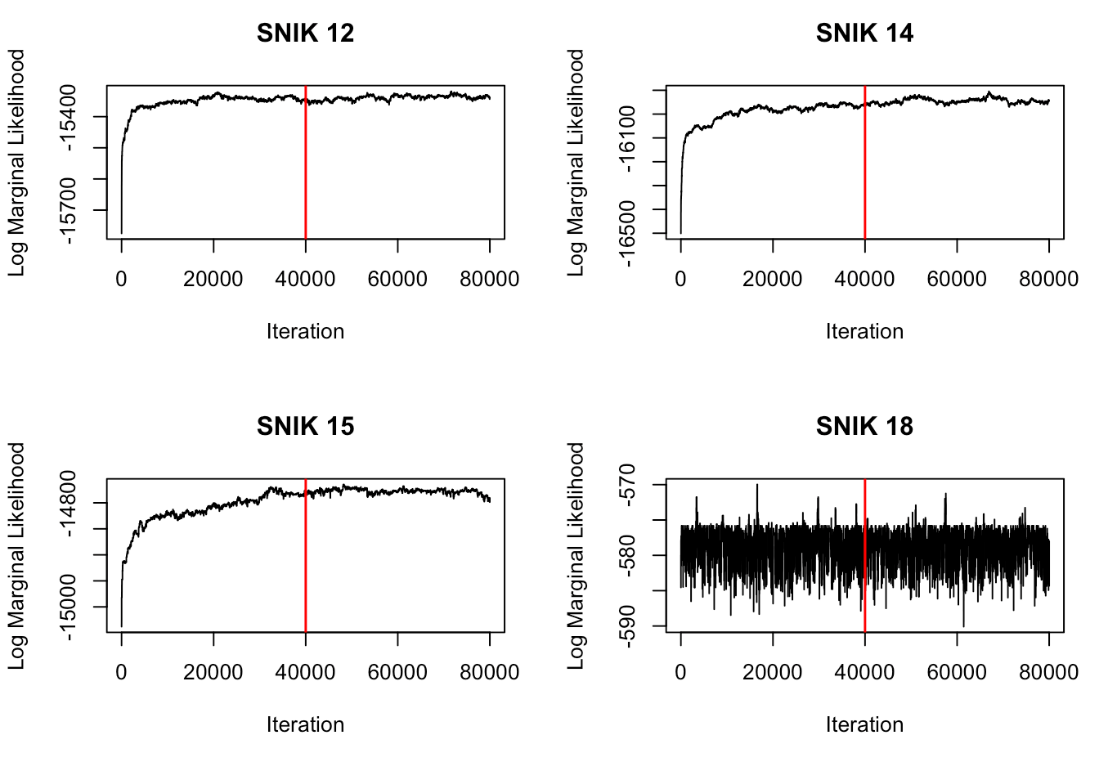

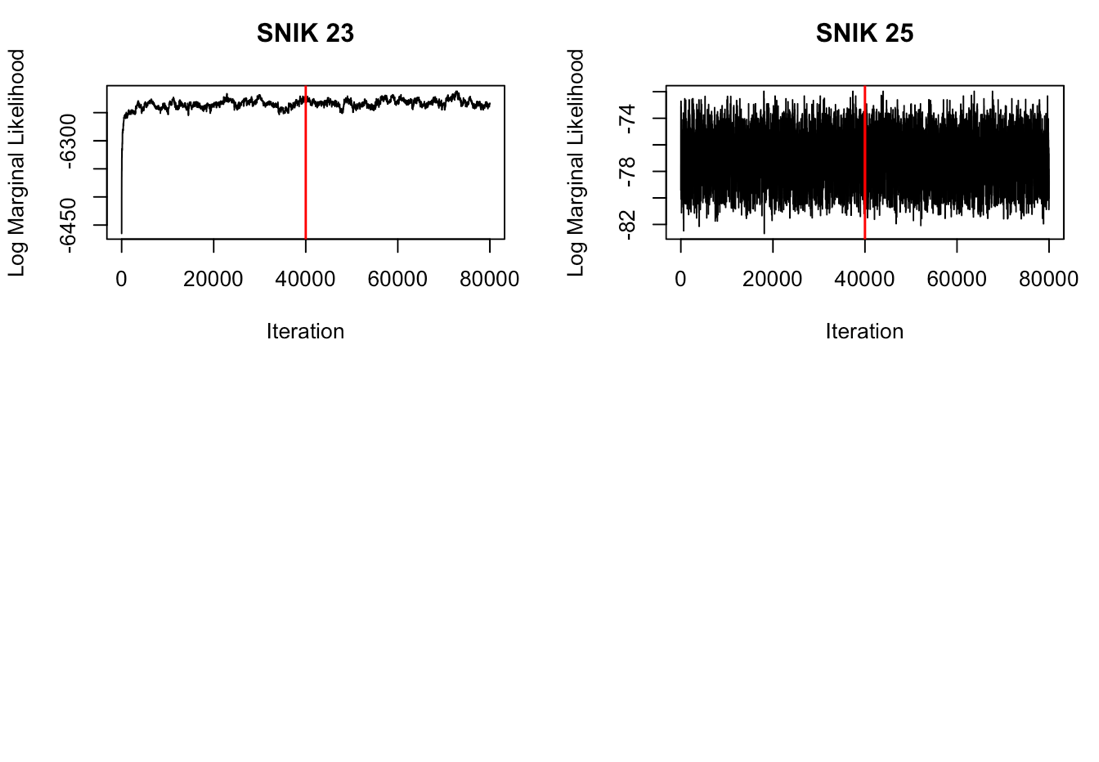

Figure S7.5 Trace-plots of the log marginal likelihood from the M4 segmentation model after analyzing 26 snail kite tracks. The MAP estimate of breakpoints was sampled from the posterior distribution, which is to the right of the burn-in (red line).

*2.2 M4 LDA model*

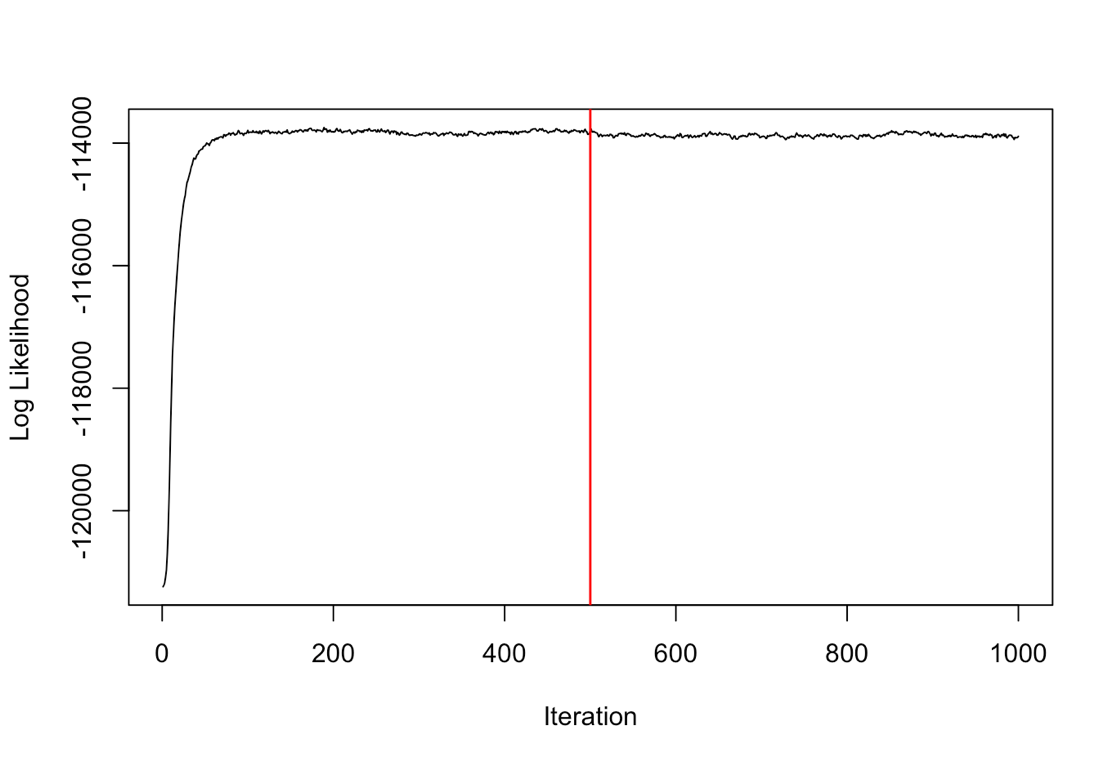

Figure S7.6 Trace-plot of the log likelihood from the M4 LDA model on all snail kite track segments pooled together. The posterior distribution is considered to be to the right of the burn-in (red line).
