## Appendix S8 for "Identifying latent behavioral states in animal movement with M4, a non-parametric Bayesian method"

Appendix S8: Comparison of Properties and Assumptions Among Analyzed Methods

Identifying latent behavioral states in animal movement with M4, a non-parametric Bayesian method

Joshua A Cullen^1*^, Caroline L Poli^2^, Robert J Fletcher, Jr.^3^, Denis Valle^1^

^1^ School of Forest Resources and Conservation, University of Florida, Gainesville, FL, USA

^2^ School of Natural Resources and Environment, University of Florida, Gainesville, FL, USA

^3^ Department of Wildlife Ecology and Conservation, University of Florida, Gainesville, FL, USA

When comparing among methods that are designed to address a given objective, it is important to understand the properties and assumptions of each of these methods. By understanding how each of the methods work, practitioners can select which method works best to analyze the data at hand as well as any specific objectives. Below, we briefly describe the properties and assumptions for each of the four methods that were compared against our proposed Bayesian M4 method in this study.

Hidden Markov models (HMM) can evaluate >2 data streams, which are used to estimate a pre-defined number of discrete states (Table S8.1). This is a discrete-time, process-based model which assumes a regular time series of data streams that are assumed to come from a correlated random walk. A transition matrix governs the probability of changing states or remaining within a given state from time $t$ to $t+1$ based on a Markov assumption. Commonly used HMMs (first order) also assume that the times spent within a state follow a geometric distribution. HMMs also typically use parametric probability distribution functions to fit each of the data streams. The number of states must be defined before running the model and models that use different numbers of states can be compared to determine the most likely number of states. Since the number of estimated parameters continues to grow as more data streams are included and for larger numbers of states, HMMs (maximum likelihood or Bayesian) are not particularly scalable for large sample sizes.

Behavioral change point analysis (BCPA) can only evaluate a single data stream to define change points of activity (Table S8.1). This data stream is typically either persistence velocity ($velocity \times cos(angle))$ or turning velocity $\left( velocity\times sin\left( angle \right) \right)$, but other variables can be used as well. BCPA assumes a continuous space-time, stationary Gaussian process where each segment of the time series is defined by a mean, standard deviation, and value of autocorrelation. BCPA also requires that the practitioner defines the size of a sliding window, which is used to help define a set of change points. Although BCPA is not able to explicitly estimate behavioral states (only track segments), additional methods can be used to cluster these segments into states. This method is scalable since it can estimate the model parameters very quickly.

Segclust2d can evaluate up to two data streams, which are first used to define a set of track segments from change points before clustering these segments into states (Table S8.1). Similar to BCPA, segclust2d relies on the assumption of a Gaussian error process that defines the time series of each segment of the data stream. Each segment is characterized by a given mean and standard deviation based on a multivariate normal distribution and each segment is assumed to belong to state $k$ of $K$ based on a multinomial distribution. To compare among multiple numbers of possible states and segments, a penalized likelihood criterion (i.e. BIC) is calculated and compared across all possible numbers of track segments and numbers of states. Since this model must estimate parameters for all possible numbers of track segments and states, segclust2d is not scalable for large sample sizes.

Expectation-maximization binary clustering (EMbC) can evaluate any number of data streams and clusters observations directly into behavioral states (Table S8.1). This method uses a particular type of Gaussian mixture model where each of *n* data streams is partitioned into a ‘low’ and ‘high’ class, resulting in a total of 2^n^ possible states. This means that the number of possible states is fixed before running the model. Unlike the use of Gaussian distributions to define the error of a stationary time series (as in BCPA and segclust2d), EMbC clusters observations directly (as part of a mixture model) without accounting for time or autocorrelation. Given the properties of this model, it is highly scalable and can quickly evaluate large datasets.

Table S8.1 A comparison of the properties and attributes for each of the models compared by this study. Each of the five methods are broken into the model categories under which they fall, where this proposed Bayesian M4 framework and segclust2d both include segmentation and clustering methods. Additional columns characterize the number of data streams that can be analyzed by each method, the type of probability distribution(s) that is fitted to the data streams, the number of possible states estimated by the model, and whether the model is scalable or not based on the duration it took to run the models on the simulated tracks.

| **Category** | **Method** | **# of data streams** | **Distributions fit to data streams** | **# of estimated states** | **Scalable** | **Reference** |
| --- | --- | --- | --- | --- | --- | --- |
| Segmentation | M4 | any | Categorical | - | yes | this paper |
|  | BCPA | 1 | Gaussian | - | yes | Gurarie et al. 2009 |
|  | Segclust2d | 2 | Gaussian | - | no | Patin et al. 2020 |
| Clustering technique | M4 | any | Categorical | any | yes | this paper |
|  | EMbC | any | Gaussian | 2^# of data streams^ | yes | Garriga et al. 2016 |
|  | Segclust2d | 2 | Gaussian | any | yes | Patin et al. 2020 |
| Process-based model | HMM | any | many | any | no | McClintock and Michelot, 2018; McClintock et al. 2020 |
