## Appendix S9 for "Identifying latent behavioral states in animal movement with M4, a non-parametric Bayesian method"

Appendix S9: LDA Results from Simulations and Snail Kite Tracks

Identifying latent behavioral states in animal movement with M4, a non-parametric Bayesian method

Joshua A Cullen^1*^, Caroline L Poli^2^, Robert J Fletcher, Jr.^3^, Denis Valle^1^

^1^ School of Forest Resources and Conservation, University of Florida, Gainesville, FL, USA

^2^ School of Natural Resources and Environment, University of Florida, Gainesville, FL, USA

^3^ Department of Wildlife Ecology and Conservation, University of Florida, Gainesville, FL, USA

Table S9.1 Mean proportions of total behavior accounted for by the two, three, or four most frequent behaviors in simulations where data streams were generated from uncommon distributions (i.e., truncated normal, beta, uniform). The most likely number of states are shown by bolded proportion values.

| Simulation ID | Track Length  (observations) | Proportion | | |
| --- | --- | --- | --- | --- |
|  |  | 2 states | 3 states | 4 states |
| 1_1 | 1000 | **0.929** | 0.998 | 1.000 |
| 2_1 | 1000 | 0.854 | **0.967** | 0.996 |
| 3_1 | 1000 | 0.830 | **0.961** | 0.997 |
| 4_1 | 1000 | 0.883 | **0.993** | 0.999 |
| 5_1 | 1000 | 0.863 | **0.990** | 0.999 |
| 1_2 | 5000 | 0.821 | **0.997** | 0.999 |
| 2_2 | 5000 | 0.868 | **0.994** | 1.000 |
| 3_2 | 5000 | 0.801 | **0.994** | 0.999 |
| 4_2 | 5000 | 0.831 | **0.989** | 0.998 |
| 5_2 | 5000 | 0.838 | **0.996** | 1.000 |
| 1_3 | 10,000 | 0.654 | 0.886 | **0.995** |
| 2_3 | 10,000 | 0.777 | **0.996** | 1.000 |
| 3_3 | 10,000 | 0.816 | **0.994** | 1.000 |
| 4_3 | 10,000 | 0.861 | **0.992** | 1.000 |
| 5_3 | 10,000 | 0.814 | **0.996** | 1.000 |
| 1_4 | 50,000 | 0.756 | **0.974** | 0.996 |
| 2_4 | 50,000 | 0.794 | **0.976** | 0.998 |
| 3_4 | 50,000 | 0.798 | **0.980** | 0.999 |
| 4_4 | 50,000 | 0.728 | **0.978** | 0.996 |
| 5_4 | 50,000 | 0.751 | **0.982** | 0.998 |

Table S9.2 Mean proportions of total behavior accounted for by the two, three, or four most frequent behaviors in simulations where data streams were generated from common distributions. The most likely number of states are shown by bolded proportion values.

| Simulation ID | Track Length  (observations) | Proportion | | |
| --- | --- | --- | --- | --- |
|  |  | 2 states | 3 states | 4 states |
| 1_1 | 1000 | 0.856 | **0.987** | 0.999 |
| 2_1 | 1000 | 0.767 | **0.991** | 0.999 |
| 3_1 | 1000 | 0.873 | **0.959** | 0.998 |
| 4_1 | 1000 | 0.898 | **0.997** | 0.999 |
| 5_1 | 1000 | 0.802 | 0.897 | **0.964** |
| 1_2 | 5000 | 0.776 | **0.991** | 0.999 |
| 2_2 | 5000 | 0.792 | **0.996** | 1.000 |
| 3_2 | 5000 | 0.749 | **0.972** | 0.998 |
| 4_2 | 5000 | 0.772 | **0.970** | 0.995 |
| 5_2 | 5000 | 0.714 | **0.990** | 0.999 |
| 1_3 | 10,000 | 0.768 | **0.993** | 0.999 |
| 2_3 | 10,000 | 0.797 | **0.992** | 0.999 |
| 3_3 | 10,000 | 0.759 | **0.951** | 0.998 |
| 4_3 | 10,000 | 0.747 | **0.988** | 0.997 |
| 5_3 | 10,000 | 0.766 | **0.995** | 1.000 |
| 1_4 | 50,000 | 0.805 | **0.985** | 1.000 |
| 2_4 | 50,000 | 0.797 | **0.984** | 0.999 |
| 3_4 | 50,000 | 0.802 | **0.983** | 0.998 |
| 4_4 | 50,000 | 0.735 | **0.962** | 0.997 |
| 5_4 | 50,000 | 0.764 | **0.970** | 0.998 |

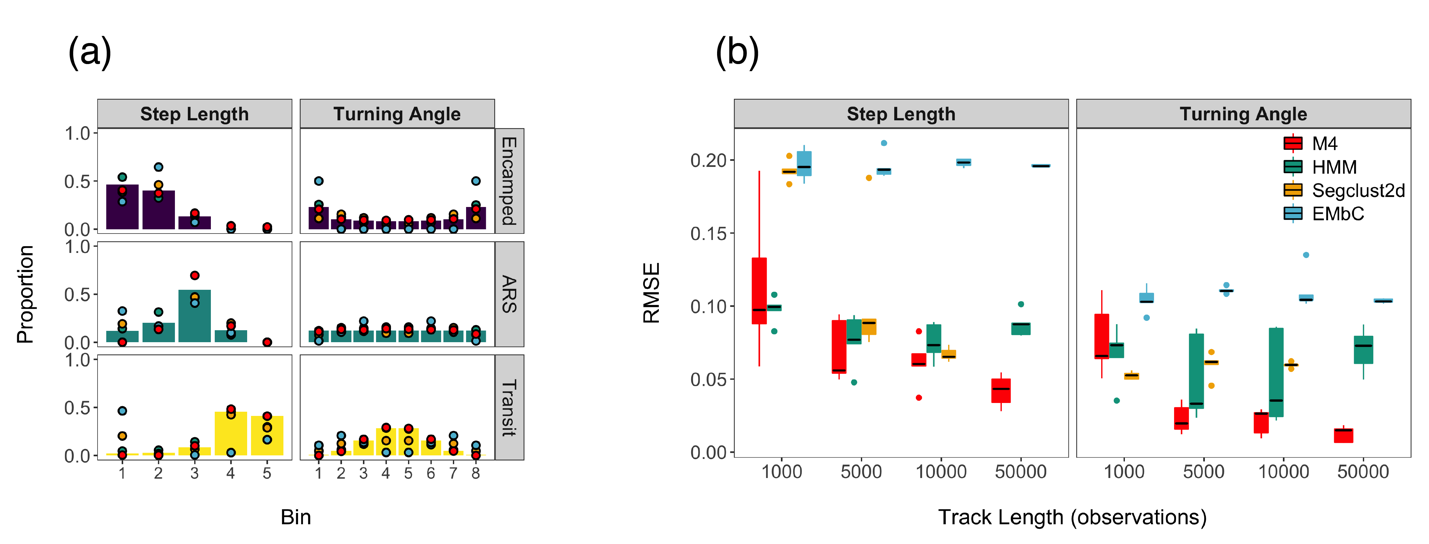

Figure S9.1. Model performance of behavioral state characterization for the M4, HMM, segclust2d, and EMbC models that analyzed simulations generated from uncommon distributions. (a) An example of state-dependent distributions are shown for a single simulated track, where true proportions of each bin are denoted by vertical bars, whereas estimates from each of the models are shown as points. Bin estimates from the HMM, segclust2d, and EMbC models were calculated by integrating the resulting parametric distributions based on the limits used to define step length and turning angle bins. Since the EMbC and segclust2d methods both analyzed the absolute value of turning angles (instead of raw turning angles), these estimates were assumed to be symmetric in the creation of these discretized distributions. (b) The difference between the model estimates from the true values (calculated as root mean square error; RMSE) for the state-dependent distributions are shown for each of the models, where results are aggregated for each data stream.

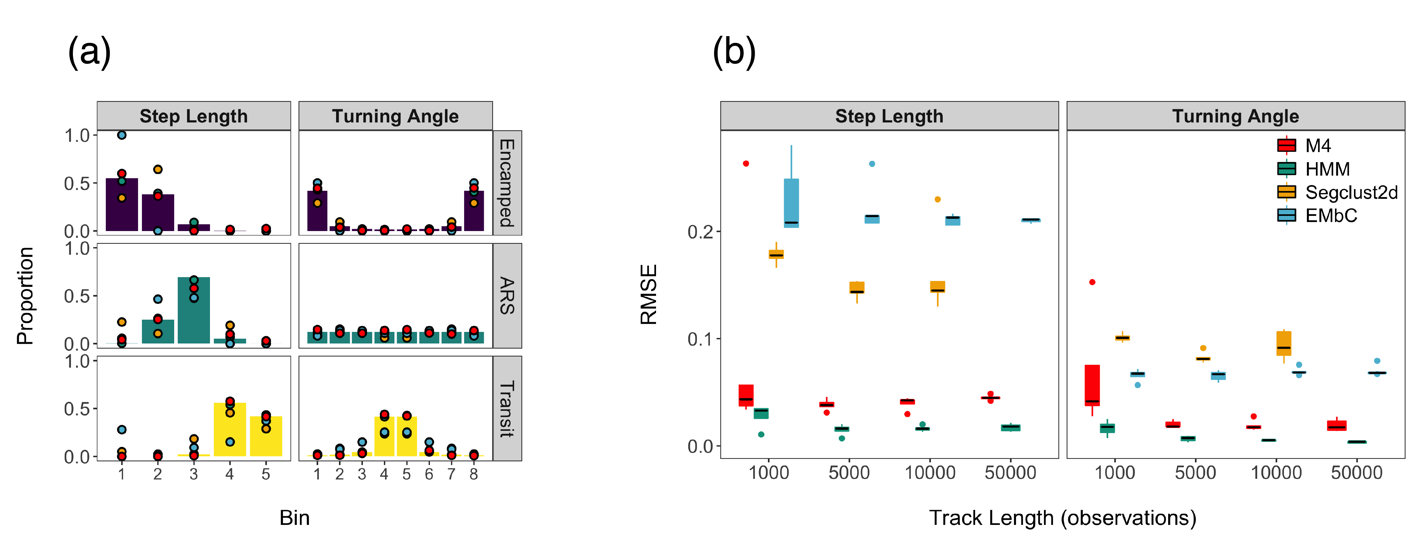

Figure S9.2 Model performance of behavioral state characterization for the M4, HMM, segclust2d, and EMbC models that analyzed simulations generated from common distributions. (a) An example of state-dependent distributions are shown for a single simulated track, where true proportions of each bin are denoted by vertical bars, whereas estimates from each of the models are shown as points. Bin estimates from HMM, segclust2d, and EMbC models were calculated by integrating the resulting parametric distributions based on the limits used to define step length and turning angle bins. Since the EMbC and segclust2d methods both analyzed the absolute value of turning angles (instead of raw turning angles), these estimates were assumed to be symmetric in the creation of these discretized distributions. (b) The difference between the model estimates from the true values (calculated as root mean square error; RMSE) for the state-dependent distributions are shown for each of the models, where results are aggregated for each data stream.

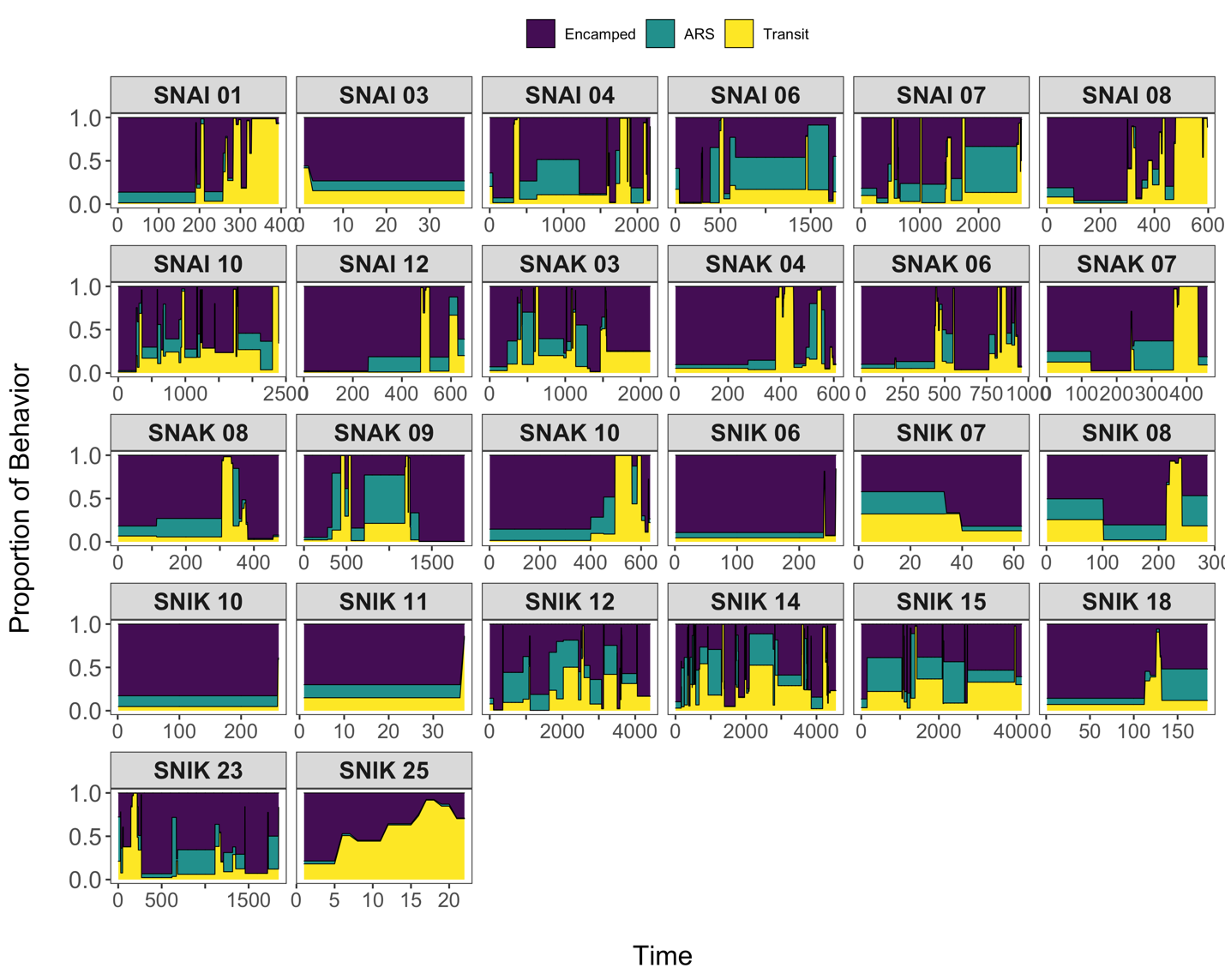

Figure S9.3 Temporal patterns of behavior proportions are shown for each of the 26 analyzed snail kites. While there is high inter-individual variability in behavior patterns, individuals that have been tagged over longer durations exhibit more frequent periods of higher activity behaviors (ARS, transit). Individuals tagged over short durations with most time spent in an encamped state likely have not yet fledged from the nest.
