## Appendix S10 for "Identifying latent behavioral states in animal movement with M4, a non-parametric Bayesian method"

Appendix S10: Estimated State-Dependent Distributions from HMM, EMbC, and Segclust2d

Identifying latent behavioral states in animal movement with M4, a non-parametric Bayesian method

Joshua A Cullen^1*^, Caroline L Poli^2^, Robert J Fletcher, Jr.^3^, Denis Valle^1^

^1^ School of Forest Resources and Conservation, University of Florida, Gainesville, FL, USA

^2^ School of Natural Resources and Environment, University of Florida, Gainesville, FL, USA

^3^ Department of Wildlife Ecology and Conservation, University of Florida, Gainesville, FL, USA

1. Simulations with uncommon distributions (i.e., truncated normal, beta, uniform)

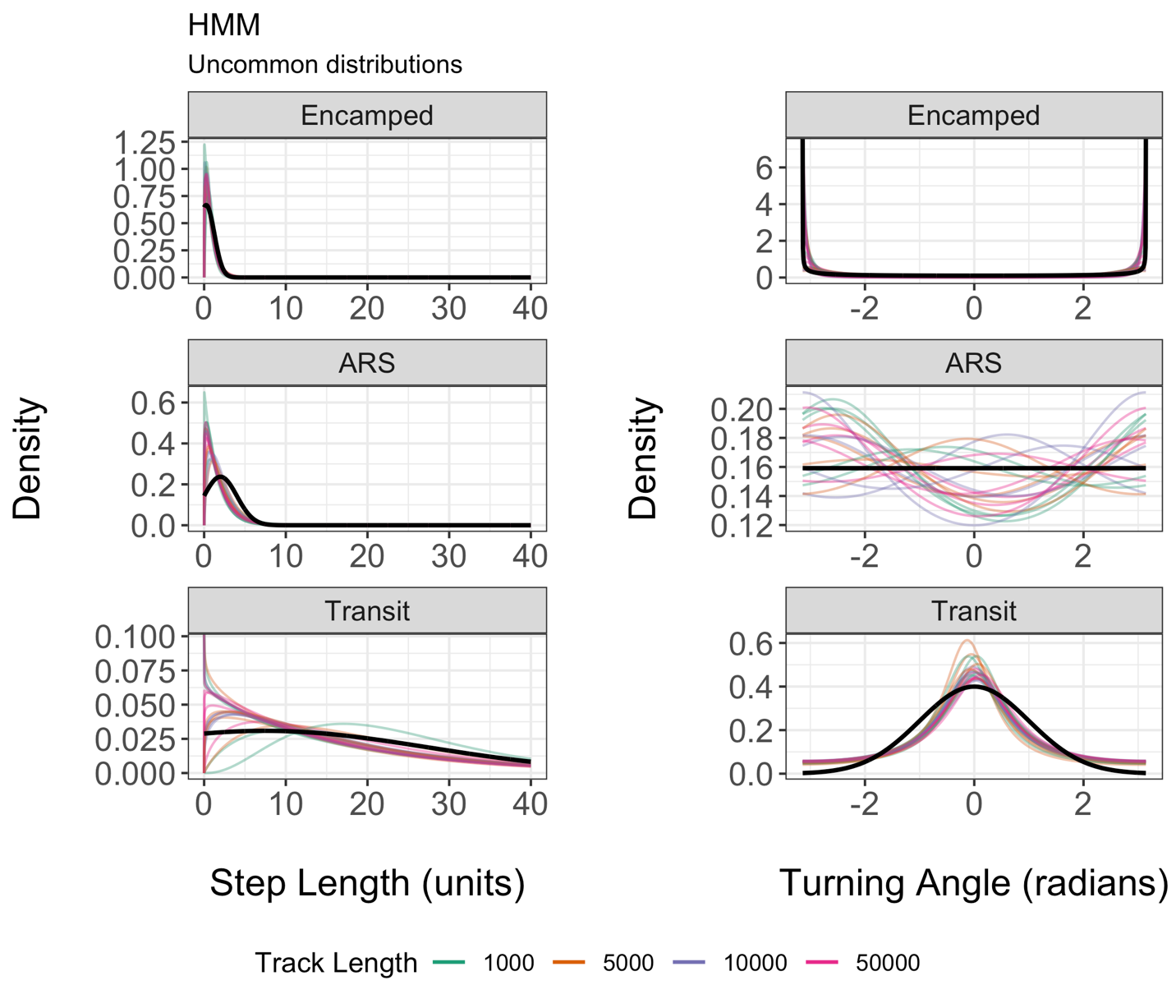

Figure S10.1 True distributions (black) are compared against estimates made across simulated tracks that were analyzed by the HMM for 3 behavioral states. The HMM fitted a gamma distribution to step lengths and a wrapped Cauchy distribution to turning angles.

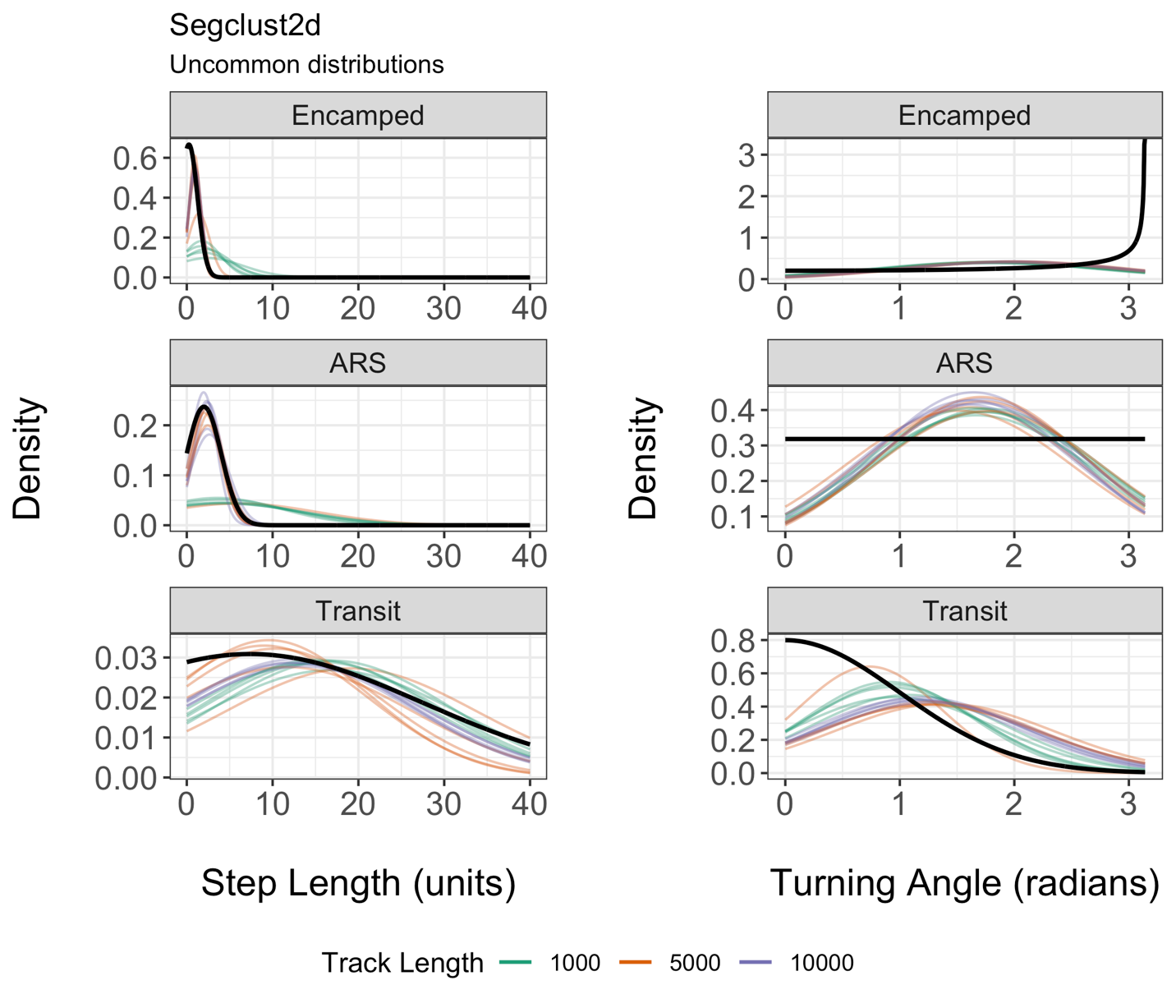

Figure S10.2 True distributions (black) are compared against estimates made across simulated tracks that were analyzed by segclust2d for 3 behavioral states. Segclust2d fitted a normal distribution to step lengths and a normal distribution to the absolute value of turning angles. The range of these estimated distributions covers all real integers, but has been restricted for comparison against the true distributions.

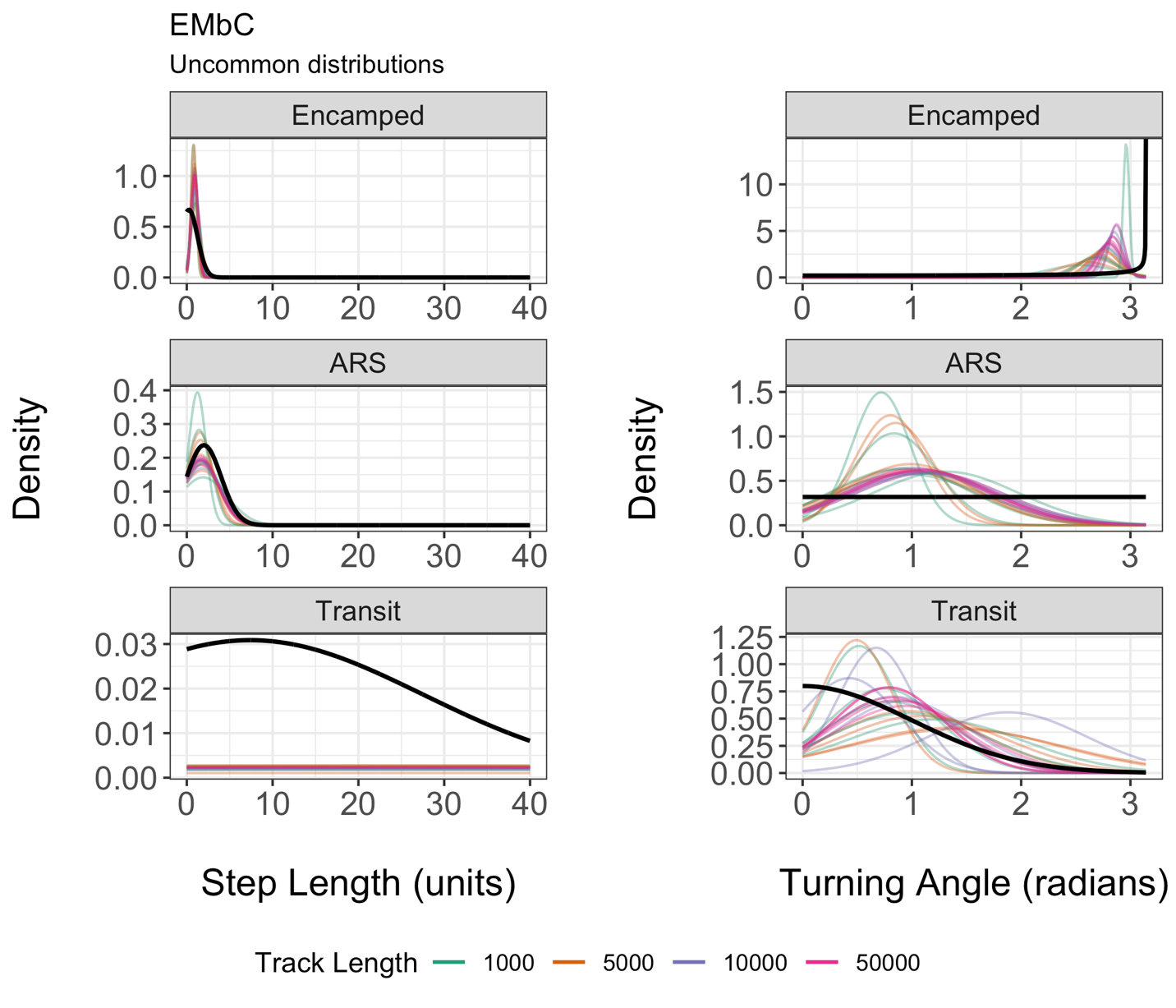

Figure S10.3 True distributions (black) are compared against estimates made across simulated tracks that were analyzed by EMbC for 3 behavioral states (HL and HH states combined into one state). EMbC fitted a normal distribution to step lengths and a normal distribution to the absolute value of turning angles. The range of these estimated distributions covers all real integers, but has been restricted for comparison against the true distributions.

2. Simulations with common distributions (i.e., gamma, wrapped Cauchy)

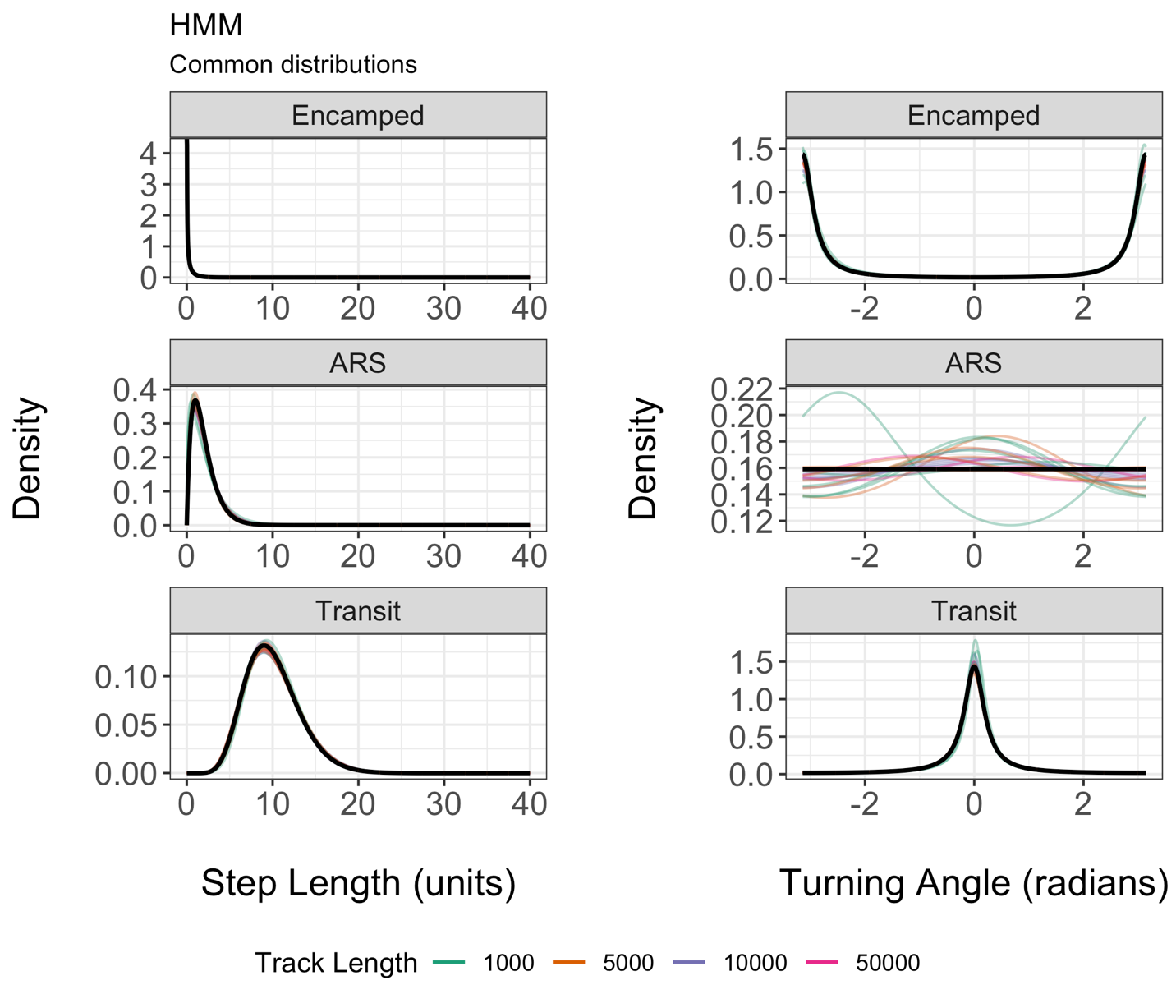

Figure S10.4 True distributions (black) are compared against estimates made across simulated tracks that were analyzed by the HMM for 3 behavioral states. The HMM fitted a gamma distribution to step lengths and a wrapped Cauchy distribution to turning angles.

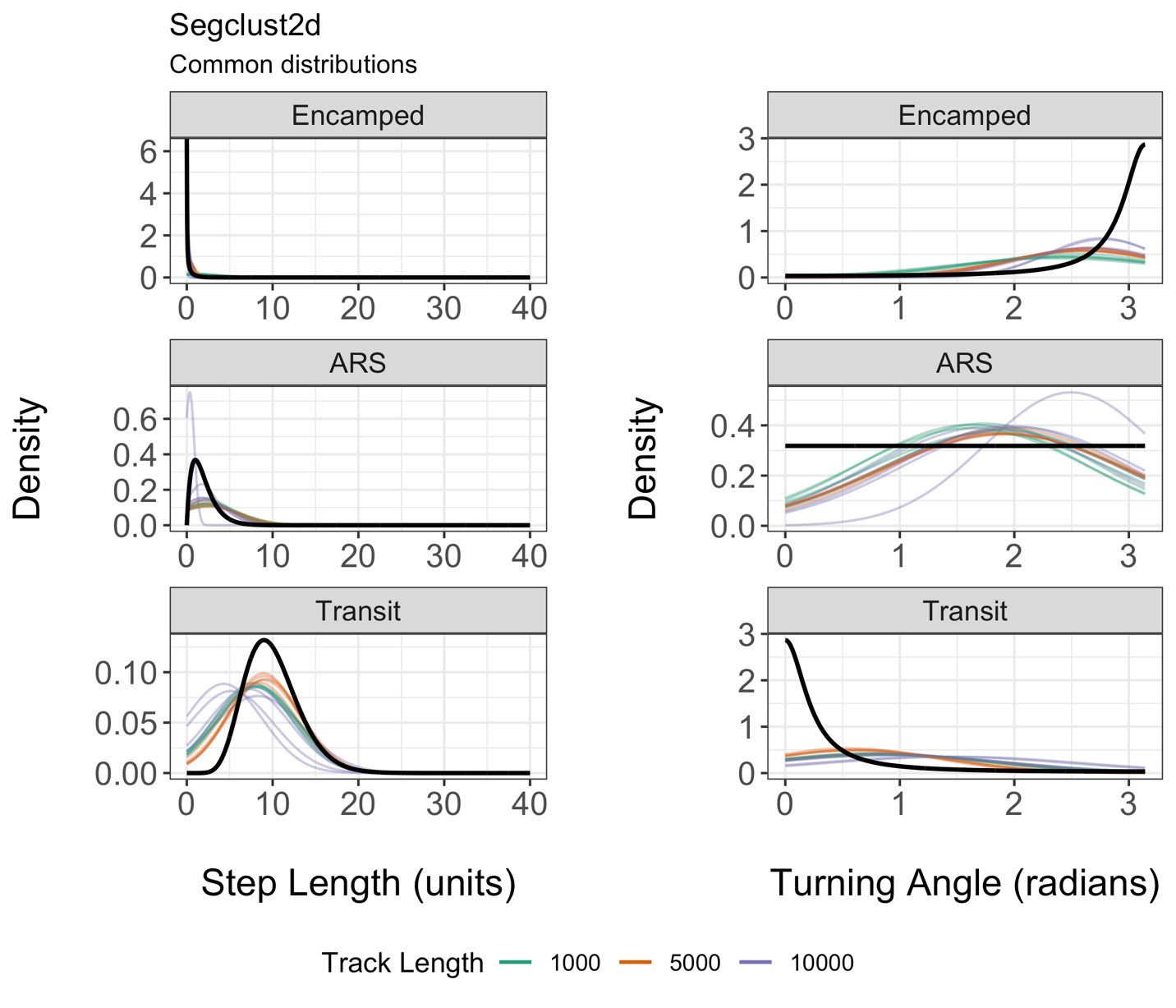

Figure S10.5 True distributions (black) are compared against estimates made across simulated tracks that were analyzed by segclust2d for 3 behavioral states. Segclust2d fitted a normal distribution to step lengths and a normal distribution to the absolute value of turning angles. The range of these estimated distributions covers all real integers, but has been restricted for comparison against the true distributions.

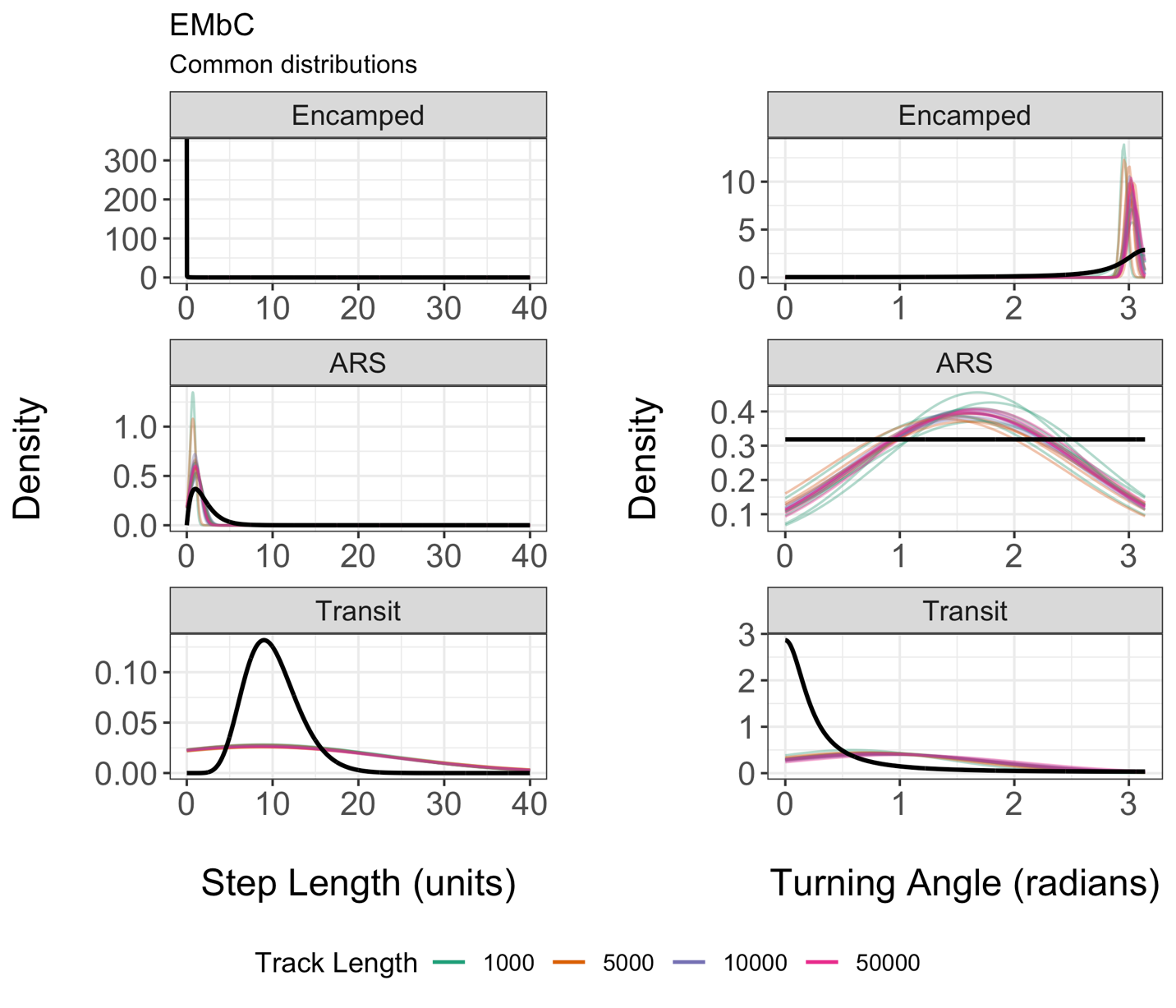

Figure S10.6 True distributions (black) are compared against estimates made across simulated tracks that were analyzed by EMbC for 3 behavioral states (HL and HH states combined into one state). EMbC fitted a normal distribution to step lengths and a normal distribution to the absolute value of turning angles. The range of these estimated distributions covers all real integers, but has been restricted for comparison against the true distributions.
